# Hepatocyte MLKL Drives Obesity-Driven Hepatocellular Carcinoma Progression via Mitochondrial Dysfunction Independent of Necroptosis in MASLD

**DOI:** 10.1101/2025.11.26.690789

**Authors:** Phoebe Ohene-Marfo, Sabira Mohammed, Chao Jiang, Shylesh Bhaskaran, Kara Kneuper, Satoshi Matsuzaki, Bo Hagy, Randal J May, Constantin Georgescu, Megan John, Julianne N. Hoang, Kevin Pham, Albert Tran, Chinthalapally V Rao, Tae Gyu Oh, Michael Kinter, Willard M Freeman, Courtney Houchen, Surendra Shukla, Kenneth Humphries, Jonathan D Wren, Sathyaseelan S. Deepa

**Affiliations:** Department of Biochemistry & Physiology, The University of Oklahoma Health Campus; Stephenson Cancer Center, The University of Oklahoma Health Campus; Aging & Metabolism Research Program, Oklahoma Medical Research Foundation, Oklahoma City, Oklahoma, USA; Section of Digestive Diseases and Nutrition, Department of Medicine, The University of Oklahoma Health Campus; Genes and Human Disease Research Program, Oklahoma Medical Research Foundation, Oklahoma City, Oklahoma, USA; University of Oklahoma, Norman, Oklahoma Medical Research Foundation, Oklahoma City, Oklahoma, USA; Hematology/Oncology Section, Department of Medicine, The University of Oklahoma Health Campus; Department of Oncology Science, The University of Oklahoma Health Campus; The Oklahoma City Veterans Affairs Medical Center, Oklahoma City, Oklahoma, USA; Oklahoma Center for Geroscience & Healthy Brain Aging, The University of Oklahoma Health Campus

**Keywords:** MASLD, HCC, hepatocyte, MLKL, MFN2, mitochondria

## Abstract

**Background and Aims:** Metabolic dysfunction-associated steatotic liver disease (MASLD) is a leading cause of hepatocellular carcinoma (HCC), particularly in obesity, yet mechanisms linking hepatocyte dysfunction to tumorigenesis remain unclear. Mixed lineage kinase domain-like protein (MLKL), the effector of necroptosis, is elevated in MASLD, but its hepatocyte-intrinsic role in obesity-driven MASLD-HCC is unknown.

**Approach and Results:** Using a long-term Western diet (WD)-induced MASLD-HCC model in hepatocyte-specific MLKL knockout (*Mlkl^HepKO^*) mice, we defined MLKL’s hepatocyte-intrinsic function. WD increased hepatocyte MLKL protein expression without detectable necroptosis activation, indicating a necroptosis-independent role. MLKL deficiency did not alter WD-induced inflammation, fibrosis, or liver injury but increased hepatic lipid accumulation while reducing lipotoxic lipid species and preserving mitochondrial function. WD-fed *Mlkl^HepKO^* mice developed fewer and smaller tumors with reduced incidence, multiplicity, proliferation, and stemness. Transcriptomic analysis revealed upregulation of mitochondrial oxidative phosphorylation pathways in *Mlkl^HepKO^* livers. WD suppressed the mitochondrial fusion protein and tumor suppressor MFN2, whereas MLKL deficiency restored MFN2 expression post-translationally. In HCC cells, MLKL deletion reduced proliferation, improved mitochondrial respiration, and decreased glycolysis; these effects were reversed by MFN2 deletion. MLKL localized to nuclear and mitochondrial compartments, consistent with organelle-intrinsic functions. The human MLKL inhibitor necrosulfonamide (NSA) suppressed HepG2 xenograft growth, and elevated MLKL expression in human HCC correlated with poorer overall survival.

**Conclusions:** Hepatocyte MLKL promotes MASLD-associated HCC through a non-necroptotic mechanism involving MFN2 suppression, impaired mitochondrial function, and increased tumor proliferation and stemness. These findings identify MLKL as a potential therapeutic target in MASLD-associated HCC.

## INTRODUCTION

Metabolic dysfunction-associated steatotic liver disease (MASLD), driven primarily by obesity and metabolic disorders, has emerged as the most prevalent liver disease, affecting nearly 38% of adults and 10% of children in the United States (1, 2). MASLD comprises a spectrum of liver diseases from hepatic steatosis, metabolic dysfunction-associated steatohepatitis (MASH), to cirrhosis. Approximately 20-30% of MASLD cases progress to MASH, of which roughly 15-20% develop advanced fibrosis or cirrhosis, with cirrhotic patients showing HCC incidence rates ranging from 0.7-2.6% annually (3, 4). Although most HCC in MASLD arises from cirrhosis, a surprisingly large fraction of MASLD-HCC progression (15-46%) occurs in the absence of cirrhosis, making prediction and surveillance more challenging (5). Despite these observations, the molecular mechanisms driving MASLD-HCC progression remain poorly understood, particularly in the context of obesity, which increases HCC risk nearly 2-fold and HCC-related mortality 4-fold compared to individuals with normal body mass index (6). This knowledge gap continues to limit the development of effective therapies for obesity-driven HCC.

Chronic inflammation is a central driver of MASH pathogenesis and progression to HCC (7). Necroptosis, a regulated form of inflammatory cell death, has been implicated in both human and experimental MASH (8). Necroptosis occurs through the sequential activation of Receptor-Interacting serine/threonine-Protein Kinase 1 (RIPK1), RIPK3, and Mixed Lineage Kinase domain-Like protein (MLKL) through phosphorylation, in response to stimuli such as tumor necrosis factor α (TNFα). Phosphorylated MLKL [Ser345 in mouse (9) or Ser358 in human (10)] undergoes oligomerization and attaches to the plasma membrane to promote cell membrane rupture and release of proinflammatory damage-associated molecular patterns (DAMPs) that activate innate immune cells to promote inflammation (11). Genetic or pharmacological inhibition of necroptosis pathway proteins reduce hepatic inflammation and injury in MASLD and HCC mouse models (12–15), supporting a role of necroptosis in these liver diseases. While hepatocytes, comprising nearly 80% of liver mass, are traditionally viewed as the main cell type undergoing necroptosis, emerging evidence challenges this notion by showing that RIPK3 is epigenetically silenced in hepatocytes and liver cancer cells, preventing MLKL activation and canonical necroptosis (16, 17). Nevertheless, MLKL expression is consistently elevated in the livers of humans and mice with MASH (8, 18, 19), although its functional relevance is unclear.

Although MLKL is classically viewed as the executioner of necroptosis, recent studies show that it also mediates non-necroptotic functions, including extracellular vesicle biogenesis and autophagy, independent of RIPK3 (20, 21). Prior studies linking necroptosis to MASLD and HCC have relied largely on whole-body knockout models, which make it difficult to define the hepatocyte-specific role of MLKL. Because RIPK3 is epigenetically silenced in hepatocytes (16), the functional relevance of MLKL in hepatocytes and liver cancer cells, where canonical necroptosis is often impaired, remains unclear. Addressing these gaps is essential to define MLKL’s context-specific role in MASLD progression and tumorigenesis.

Here, we examined the hepatocyte-specific role of MLKL in obesity-driven MASH-HCC using Western diet (WD), a well-established model of MASLD-associated HCC. Extended WD feeding (∼15 months) induces obesity, MASH, fibrosis, and spontaneous HCC, closely reflecting human disease. Notably, WD-induced HCC in mice arises around 17 months of age, roughly equivalent to 50 years in humans, when HCC incidence begins to increase (22), making this model highly relevant for interrogating hepatocyte-intrinsic MLKL function. Our data show that hepatocyte MLKL accelerates WD-induced HCC by impairing mitochondrial function/dynamics and enhancing cell cycle progression, independent of canonical necroptosis.

## METHODS

### Animals, Diets, and Hepatocyte-Specific *Mlkl* Deletion

All procedures were approved by the University of Oklahoma Health Campus IACUC. Male *Mlkl^fl/fl^* mice (originally generated by Murphy et al. 2013) (9) on a C57BL/6J background were injected with AAV8-TBG-Cre (hepatocyte-specific) or AAV8-TBG-Null (Vector Biolabs, PA, USA) via tail vein at 1.5 months of age to generate hepatocyte-specific *Mlkl* knockout (*Mlkl^HepKO^*) or control mice. Beginning at 2.5 months of age, mice were fed ad libitum either a control diet (CD, Teklad 7013 NIH-31, Research Diets) or a Western diet (WD, Research Diets, D22090208i custom made to match SF-11-078) (23) for 4, 8, or 15 months. Five mice were housed per ventilated cage under a 12-h light/dark cycle at 20 ± 2°C. Diet composition is provided in Supplementary **Table S3**.

### Hepatocyte Isolation

Hepatocytes and non-parenchymal cells were isolated using the Liver Perfusion Kit (130-128-030; Miltenyi Biotec, CA, USA) by following the manufacturer’s instructions. Briefly, left lateral lobe was used for perfusion using the gentleMACS Program 37C_m_LIPK_1, followed by digestion using the gentleMACS Program LIPK_HR_1. The hepatocytes pellet obtained was counted and plated onto collagen (C3867-1VL, Sigma Aldrich) coated plates in HepatoZYME-SFM medium (17705021, ThermoFisher Scientific). The medium was changed to DMEM (11885084, ThermoFisher Scientific) after 3 h of plating. Experiments were performed after 36 h of plating.

### Bulk RNA Sequencing

RNA-seq libraries were prepared by the University of Oklahoma Health Campus Institutional Core Facility and sequenced on an Illumina NextSeq 2000. Differential expression was assessed using limma-voom. Pathway enrichment was performed using fgsea, ReactomePA, and viewPathway.

### Statistics

Data are represented as mean ± SEM. Data normality was assessed using the Shapiro-Wilk test prior to the application of parametric tests; non-parametric alternatives were used for datasets not meeting normality assumptions. Two-tailed unpaired t-test or One-way ANOVA was used to analyze data with GraphPad Prism. P< 0.05 is considered as statistically significant. Proportion of tumor incidence was analyzed using Fisher’s exact test; p<0.05 was considered statistically significant. For bulk RNA sequencing data analysis, moderate t-test p-values were adjusted for multiple testing using the false discovery rate (FDR) method. FDR (q-value) <0.05 and absolute log2 fold change above 1 were used as criteria to filter significantly differentiated genes.

Complete procedural details and reagent lists are provided in Supplementary Methods.

## RESULTS

### WD induces hepatocyte MLKL expression without activating canonical necroptosis

Western blot analysis confirmed efficient hepatocyte-specific deletion of MLKL in *Mlkl^HepKO^* mice. MLKL expression was reduced in liver tissue and absent in isolated hepatocytes from *Mlkl^HepKO^*mice but remained unchanged in extrahepatic tissues and liver non-parenchymal cells (NPC) (**Figures 1A, S1A-C**). To determine the role of hepatocyte MLKL during diet induced MASLD to HCC progression, control and *Mlkl^HepKO^*mice were fed a control diet (CD) or WD beginning at 2.5 months of age and analyzed after 4, 8, or 15 months (**Figure 1B**). WD increased body weight and liver weight to a similar extent in both genotypes (**Figures 1C, D**). WD feeding induced hepatic MLKL expression in control mice, with approximately 2-fold increases at 4 and 8 months and a 4-fold increase at 15 months (**Figure 1E**). MLKL expression was similarly increased in primary hepatocytes isolated from WD-fed mice (**Figure S1D**). Because MLKL oligomerization and necrosome assembly are membrane-associated events, we examined both whole liver lysates (soluble) and the corresponding insoluble (membrane-enriched) fraction; Western blot analysis showed that phosphorylated MLKL (Ser345), MLKL oligomers, total RIPK3, and phosphorylated RIPK1 were undetectable in either fraction from WD-fed livers, whereas all markers were detected in positive controls (**Figure 1E, S1E, S1F**). RIPK1 expression was comparable between groups (**Figure 1E, S1F**). *Ripk3* mRNA was undetectable in primary hepatocytes but detected in liver NPCs, where its expression was not altered by genotype (**Figure S1G**), and hepatic *Ripk3* transcript was not altered by WD feeding, except for an increase at 8 months in *Mlkl^HepKO^* mice (**Figure S1H**). To further confirm that MLKL’s effects occur through a non-necroptotic mechanism given the absence of RIPK3, we restored RIPK3 expression in human HepG2 and mouse AML12 cells. RIPK3 restoration promoted MLKL phosphorylation, reduced cell proliferation, and increased lactate dehydrogenase (LDH) release in HepG2 cells (**Figures S1I**), while *Ripk3* overexpression in AML12 cells sensitized cells to necroptosis inducer-mediated MLKL phosphorylation (**Figure S1J**), confirming restoration of RIPK3-dependent MLKL activation.

**Figure 1.**
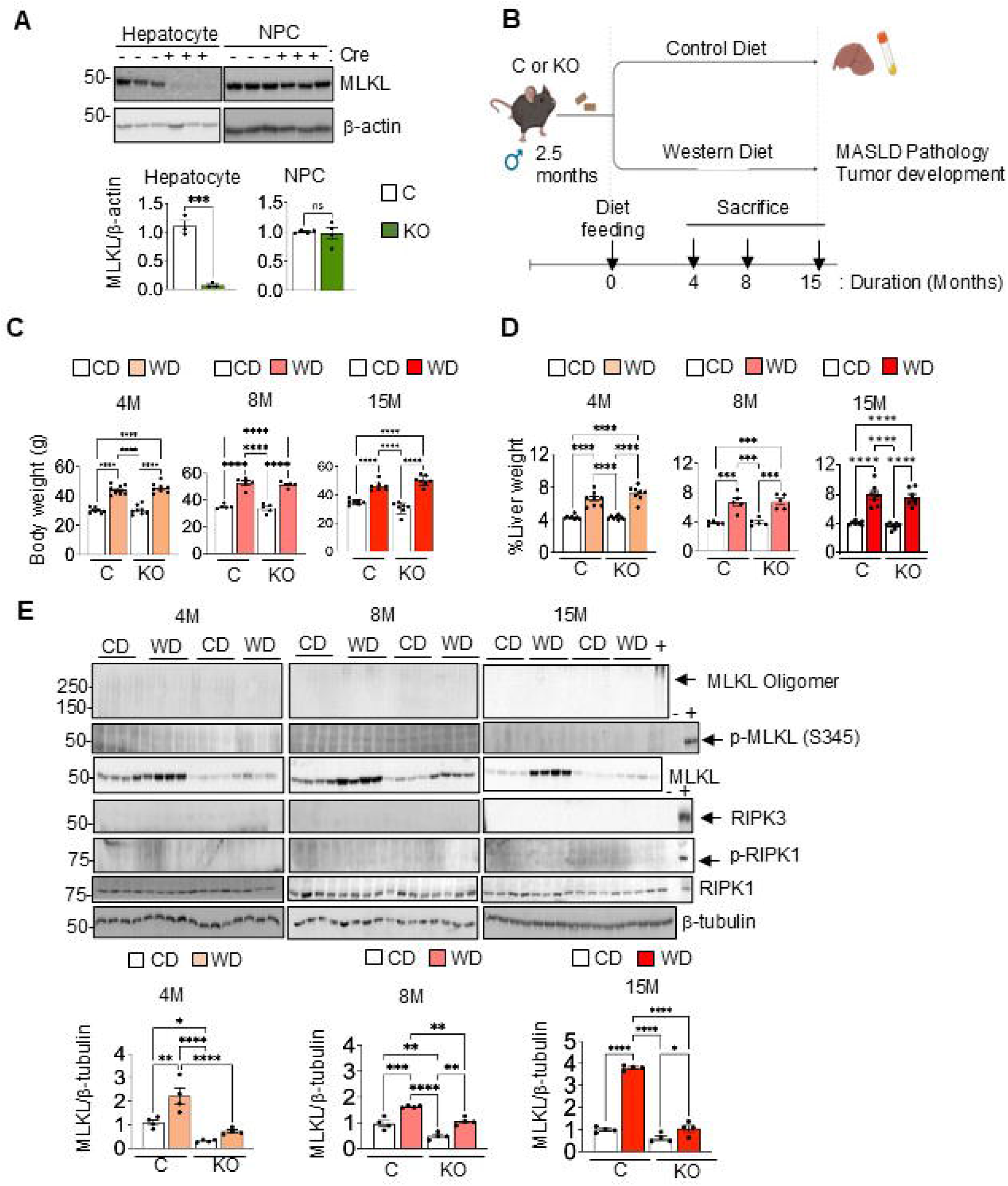
(A) Immunoblots showing the expression of MLKL and β-actin in control (C) or *Mlkl^HepKO^* (KO) hepatocytes or non-parenchymal cells (NPCs) (*Top*); Graphical representation of quantified blots normalized to β-actin (*bottom*). (B) Schematic of experimental design. (C) Body weight of mice (in grams). (D) The ratio of liver weight normalized to the percentage of body weight after 4, 8 and 15 months of control diet (CD) or Western diet (WD) feeding (n=8, 5 or 7 mice/group for 4, 8- or 15-month diet feeding respectively). (E) *Top*: Immunoblots showing expression of MLKL oligomer, p-MLKL (S345), RIPK3, p-RIPK1, RIPK1 and β-tubulin (loading control) in liver tissues of control and *Mlkl^HepKO^* mice fed either control diet (CD) or Western diet (WD) for 4-, 8-, or 15-months. Positive controls (+): *Sod1^−/−^* mouse liver lysate for MLKL oligomer blot, necroptotic BV2 cell lysate for p-MLKL(S345) and p-RIPK1, and *Ripk3*-overexpressing HepG2 cell lysate for RIPK3 blots. Negative controls (−): Liver lysates from *Ripk3^−/−^* or *Mlkl^−/−^* mouse for RIPK3 or p-MLKL(S345) blots respectively. *Bottom:* Graphical representation of quantified immunoblots for MLKL normalized to β-tubulin and expressed as fold change. Each dot represents an individual mouse (n= 4 mice/ group). Data represented as mean SEM. Data points marked with asterisks (*) indicate statistically significant differences comparing the mean of each group with that of every other group. (A) Two-tailed unpaired t-test; (C-E) One-way ANOVA, P< 0.05; *p<0.05, **p< 0.01, ***p<0.001, ****p<0.0001.

To determine whether MLKL deficiency shifts cell death toward alternative pathways, we assessed apoptosis by measuring cleaved caspase-3 via immnuohistochemical staining, pyroptosis by determining the cleaved GSDMD/GSDMD ratio via Western blotting, and ferroptosis by evaluating GPX4, ACSL4, and 4-HNE via Western blotting, and F2-isoprostane levels were quantified by GC/MS analysis. None of these markers differed between WD-fed control and *Mlkl^HepKO^*mice (**Figures S1K-P**). Since ferroptosis markers were unchanged, we next assessed whether redox homeostasis pathways using hepatic mass spectrometry-based proteomic analysis. Although catalase and glutathione-disulfide reductase (GSR) were reduced in WD-*Mlkl^HepKO^*livers, other WD-induced antioxidant changes were similar between genotypes, suggesting changes in specific components of the hepatic antioxidant response (**Figures S1Q**).

### Hepatocyte MLKL deficiency exacerbates hepatic steatosis without worsening WD-induced liver injury or fibrosis

H&E staining demonstrated marked hepatic steatosis in both control and *Mlkl^HepKO^* mice following WD feeding. WD-fed *Mlkl^HepKO^*mice exhibited larger lipid droplets than WD-fed controls at 4 and 8 months (**Figures 2A, S2A**). Although hepatic triglyceride content was comparable at earlier time points, *Mlkl^HepKO^* mice displayed significantly greater triglyceride accumulation after 15 months of WD feeding (**Figure 2B**). Expression of the lipid droplet-associated protein Perilipin-2 (PLIN2) by western blotting was also increased in WD-fed *Mlkl^HepKO^* livers at 8 and 15 months (**Figure S2B**). Analysis of lipid metabolic pathway genes revealed increased expression of cluster of differentiation 36 (*Cd36*), acetyl CoA carboxylase 1 (*Acc1*), stearoyl-CoA desaturase 1 (*Scd1*), and peroxisome proliferator-activated receptor gamma (*Pparγ)* in WD-fed *Mlkl^HepKO^* livers compared with WD-fed controls, whereas fatty acid transport protein 2 (*Fatp2*), carnitine palmitoyltransferase 1a (*Cpt1a*) were similar between the WD-fed groups (**Figure 2C**). Picrosirius red staining, hepatic hydroxyproline content, and fibrosis-related gene expression were similar between genotypes at all time points examined, with the exception of increased fibrosis-related gene expression at 8 months (**Figures 2D, 2E, S2C**). Plasma alanine aminotransferase (ALT) levels were higher in WD-fed *Mlkl^HepKO^* mice at 4 months but were similar to those in WD-fed control mice at 8 and 15 months (**Figure 2F**).

**Figure 2.**
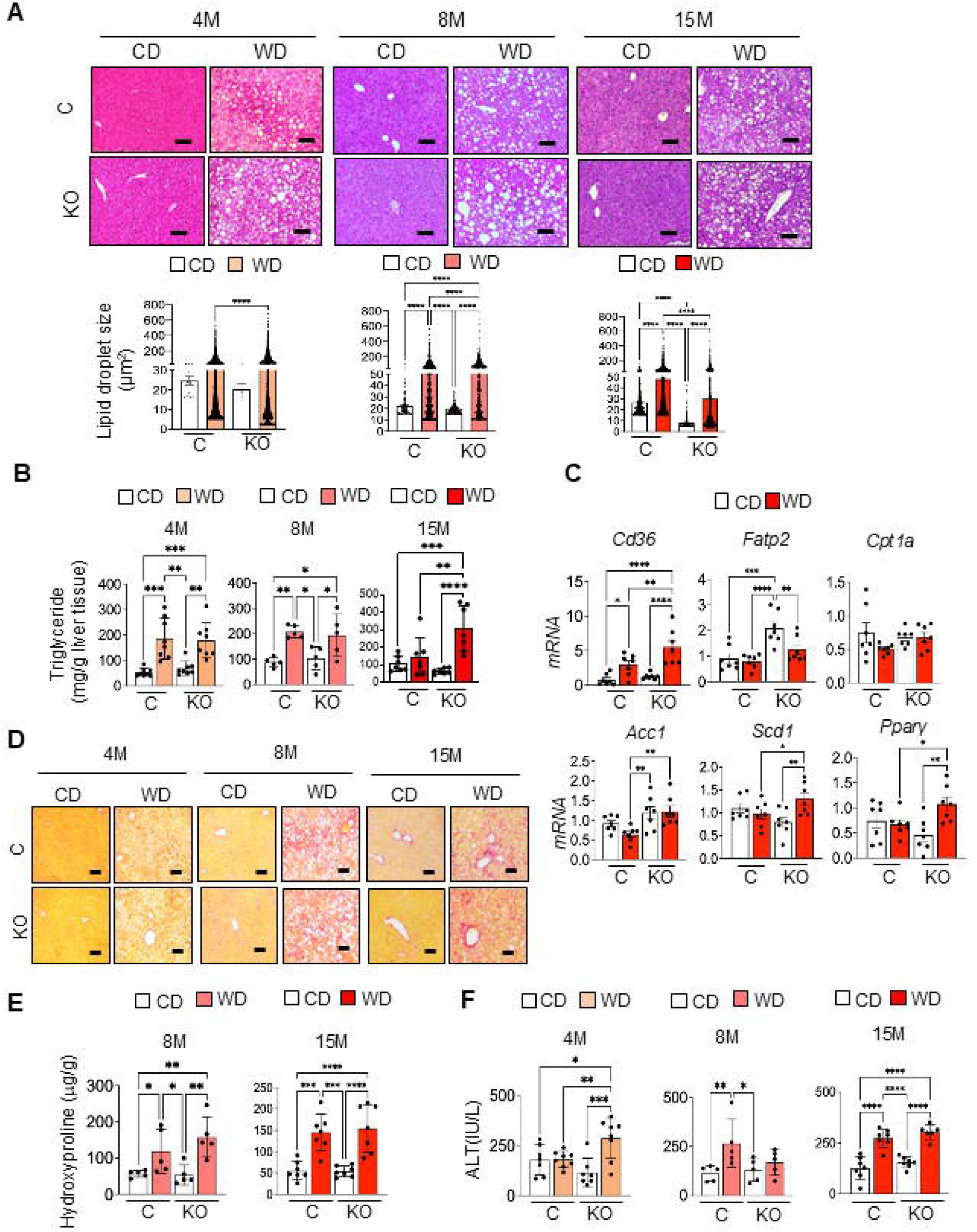
Data from liver tissues of control (C) and *Mlkl^HepKO^* (KO) mice fed Control diet (CD) or Western diet (WD): (A) *Top:* Representative images of H&E-stained liver sections (10X objective; n=3/group), *Bottom:* Mean lipid droplet surface area (μm^2^); Each dot represents an individual lipid droplet measurement (n=3/group). (B) Quantification of triglyceride levels (mg/g) (n=8, 5 or 7 mice /group for 4, 8- or 15-month diet feeding respectively). (C) Transcript levels of genes involved in: Fatty acid transport (*Cd36*, *Fatp2*); Fatty acid oxidation (*Cpt1a*), and Fatty acid synthesis and storage (*Acc1, Scd1, Pparγ)*, 15 months post diet feeding represented as fold change normalized to β-microglobulin. Each dot represents an individual mouse (n=7 mice/group). (D) Representative picrosirius red staining of liver sections [(n=3/group), 10X objective]. (E) Estimation of total hydroxyproline content (μg/g liver). Each dot represents an individual mouse (n=5 or 7 mice/ group for 8- or 15-month diet feeding respectively). (F) Serum ALT levels (IU/L), Each dot represents an individual mouse (n=7, 5 or 7 mice/ group for 4-, 8- or 15-month diet feeding respectively). Data represented as mean SEM. Data points marked with asterisks (*) indicate statistically significant differences comparing the mean of each group with that of every other group. One-way ANOVA, P< 0.05; *p<0.05, **p< 0.01, ***p<0.001, ****p<0.0001.

### Hepatocyte MLKL deficiency promotes lipid storage while preserving mitochondrial function and reducing lipotoxicity

To determine whether the steatotic phenotype was hepatocyte intrinsic, MLKL was silenced in AML12 hepatocytes using siRNA approach (∼80% knockdown; **Figure S3A**) and steatosis was induced with palmitic acid/oleic acid (PA/OA). MLKL knockdown increased intracellular lipid accumulation as assessed by Oil Red O staining (**Figure 3A**). Similar findings were observed in HepG2 cells and primary mouse hepatocytes (**Figures 3B, S3B, S3C**). In AML12 cells, MLKL knockdown increased expression of genes involved in fatty acid uptake (*Cd36*, *Fatp5*), lipid droplet formation (*Plin2*, cell death inducing DFFA-like effector C, *Cidec*), lipid mobilization (Patatin-like phospholipase domain-containing protein 2, *Pnpla2*; hormone-sensitive lipase, *Lipe*), and fatty acid oxidation (*Cpt1a*) following PA/OA treatment (**Figure 3C**), indicating broad remodeling of lipid metabolic pathways. MLKL knockdown also preserved mitochondrial respiration during palmitate induced lipotoxic stress as determined by Seahorse XF Cell Mito Stress Test (**Figure 3D**). LC-MS/MS-based lipidomic analysis of liver tissue revealed that several lipotoxic lipid species, including ceramide (d30:0), diacylglycerol (34:1), and cholesterol ester (17:1), were reduced in WD-fed *Mlkl^HepKO^*livers compared with controls (**Figures 3E, S3D**). Transmission electron microscopy of liver sections revealed reduced lipid droplet-mitochondrial distance, quantified using ImageJ, in WD-fed *Mlkl^HepKO^*livers, suggesting increased proximity (**Figure 3F**).

**Figure 3.**
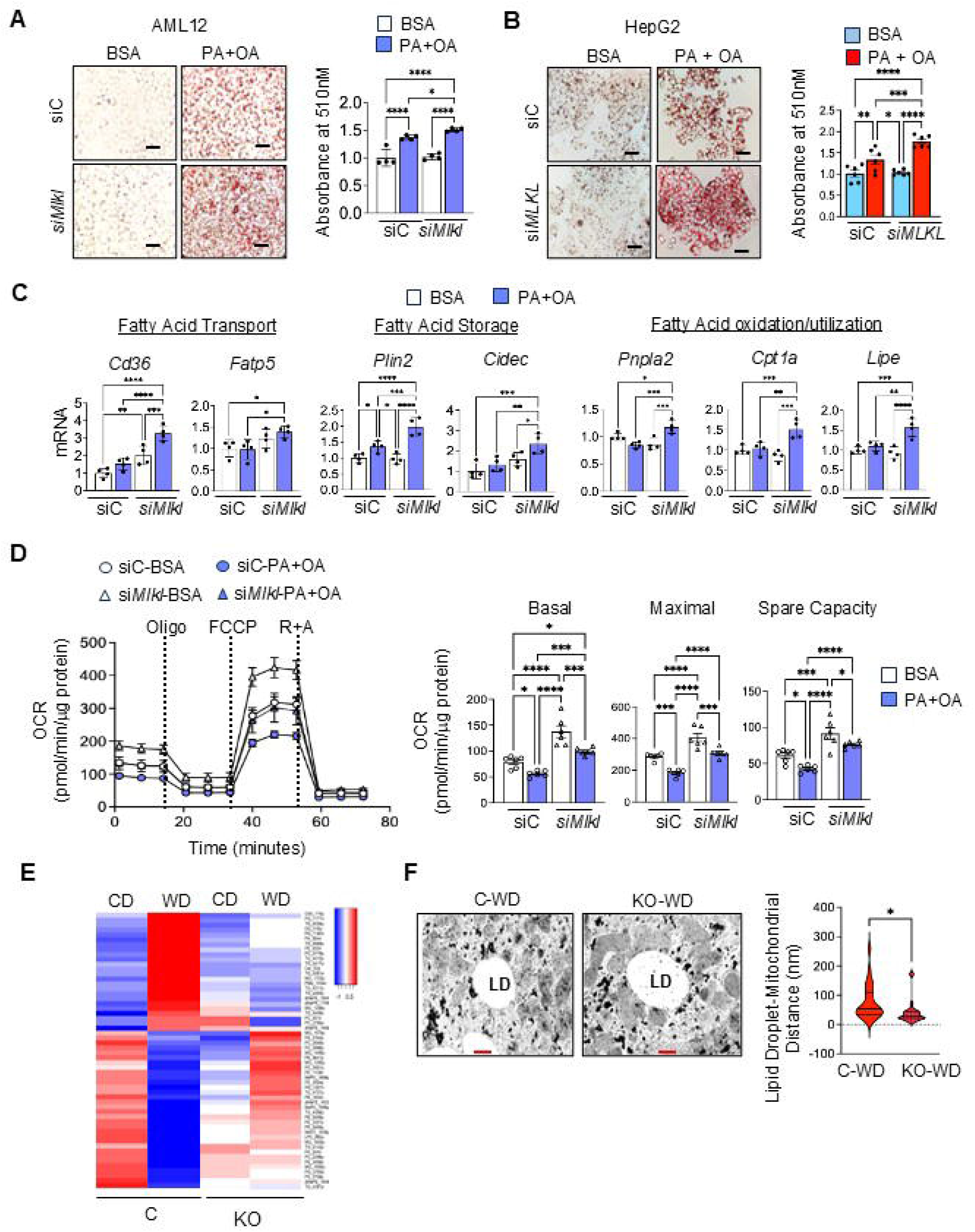
(A-B) *Left:* Images showing Oil red O staining in control (siC) and MLKL knockdown (si*Mlkl,* si*MLKL)* AML12 or HepG2 cells treated with BSA or a mixture of BSA-conjugated palmitic acid (PA) and oleic acid (OA) (BSA-PA:OA, 250:500 μM) for 48 hours (20X objective). *Right*: Graphical representation of the absorbances of quantified Oil red O extracted from AML12 or HepG2 cells expressed as fold change. (C) Transcript levels of genes involved in: Fatty acid transport (*Cd36, Fatp5*); Fatty acid storage (*Plin 2, Cidec*); Fatty Acid oxidation and utilization (*Pnpla2, Cpt1a, and Lipe*) in AML12 cells treated with BSA or BSA-PA:OA (250:500 μM) for 48 h. Each dot represents an individual mouse (n=4/group). (D) Mitostress test in siC or si*Mlkl* AML12 cells treated with either BSA (50 μM) or BSA-PA (50 μM). *Left:* Representative oxygen consumption rate (OCR) curve, with sequential addition of oligomycin, FCCP, and rotenone/antimycin A, and normalized to μg protein. *Right:* Bar graph summarizing the quantified respiratory parameters. Each dot represents an individual experiment (n=6/group). (E) Heat map representing hepatic lipidomic changes in control (C) or *Mlkl^HepKO^* (KO) mice fed either Control diet (CD) or Western diet (WD) for 4 months. Red and blue colors represent lipid species that are significantly upregulated or downregulated respectively. (F) *Left:* Transmission electron microscopy images of liver tissue from control (C) and *Mlkl^HepKO^*(KO) mice fed WD for 4 months; (Magnification: 1000X; Scale bar: 1 μm). *Right:* Graphical representation of the ultrastructure measurement of lipid droplet-mitochondrial distance (nm). LD: lipid droplet. Data represents as mean SEM. Data points marked with asterisks (*) indicate statistically significant differences comparing the mean of each group with that of every other group, (n=3 mice/group). (A-D) One-way ANOVA, (F) Two-tailed unpaired t-test. P 0.05. *p<0.05, **p< 0.01, ***p<0.001, ****p<0.0001.

### Hepatocyte MLKL deficiency reduces WD-induced HCC

Gross examination of liver revealed fewer and smaller tumors in 15-month WD-fed *Mlkl^HepKO^* mice compared with controls (**Figure 4A**). Tumor incidence was significantly reduced in WD-fed *Mlkl^HepKO^* mice compared with WD-fed control mice [3/8 (37.5%) vs 7/8 (87.5%), respectively] (**Figure 4B**). Tumor multiplicity and tumor size were also significantly decreased in WD-fed *Mlkl^HepKO^*mice (**Figures 4C, S4A**). While WD feeding markedly increased hepatic alpha-fetoprotein (AFP) levels in control mice, this increase was significantly attenuated in *Mlkl^HepKO^*mice (**Figure 4D**). Glypican-3 (GPC-3) immunostaining was also reduced in tumors from *Mlkl^HepKO^*mice (**Figure S4B**). Moreover, Ki-67-positive cells were significantly decreased in *Mlkl^HepKO^* tumors compared with controls (**Figure 4E**), however, Ki-67-positive cells were similar in liver tissue at 4, 8, and 15 months (**Figure S4C**). WD-induced increases in stem cell markers doublecortin-like kinase 1 (DCLK1) and octamer-binding transcription factor 4 (OCT4) were attenuated in *Mlkl^HepKO^* livers (**Figure 4F**).

**Figure 4.**
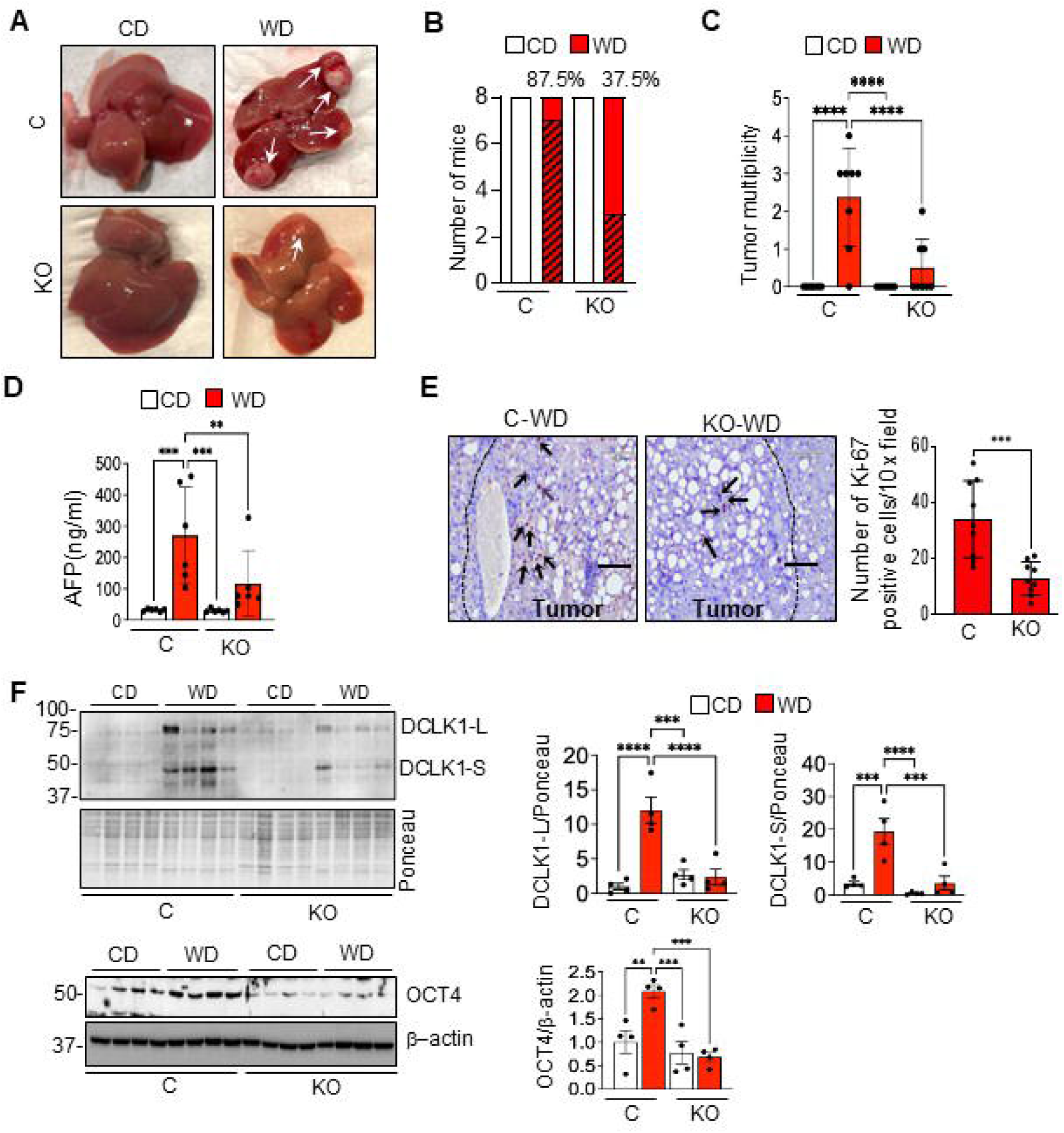
Data from control (C) or *Mlkl^HepKO^* (KO) mice fed Control diet (CD) or Western diet (WD) for 15 months: (A) Representative images of liver tumors (white arrows). Graphical representations of quantifications of: (B) tumor incidence (hatched areas indicate the tumor incidence rate within each WD group), and (C) tumor multiplicity. Each dot represents an individual mouse (n=8 mice/group). (D) Plasma AFP levels; each dot represents an individual mouse (n=6 mice/group). (E) *Left:* Representative IHC images for Ki-67 staining; *Right:* Quantified Ki-67 positive cells per 10X field. Dark brown spots in tumor regions represent positive stains for Ki-67 (Scale Bar: 200 μm); dotted lines separate tumor from non-tumor region, (n=3 mice/group). Each dot represents an individual image taken. (F) *Left:* Immunoblots showing DCLK1 (long and short isoform) and Ponceau (loading control) (*top*) and OCT4 and β-actin (loading control) (*bottom*). *Right*: Graphical representation of quantified blots normalized to Ponceau or β-actin and expressed as fold change. Each dot represents an individual mouse (n=4 mice/group). Data points marked with asterisks (*) indicate statistically significant differences comparing the mean of each group with that of every other group. (E) Two-tailed unpaired t-test. (A, C, D & F) One-way ANOVA, P<0.05. *p<0.05, **p< 0.01, ***p<0.001, ****p<0.0001. (B) Fisher’s exact test, p=0.001

### MLKL promotes proliferation and clonogenicity in HCC cells and is elevated in human HCC

To determine whether the metabolic effects observed in non-transformed AML12 hepatocytes extend to malignant hepatocytes, we next used human HCC cell line, HepG2. While AML12 cells model hepatocyte-intrinsic metabolic responses under steatotic conditions, HepG2 cells provide a clinically relevant model to assess MLKL-dependent effects on HCC cell proliferation, and clonogenicity. MLKL protein expression was increased in HepG2 cells compared with normal human liver THLE-2 cells (**Figure 5A**). Additional human and murine HCC cell lines exhibited variable MLKL expression, whereas RIPK3 remained undetectable (**Figure S5A**). Transcriptomic analysis demonstrated downregulation of cell cycle pathways and upregulation of tissue organization and immune-related pathways following MLKL depletion (**Figures S5B, S5C**). Cell cycle analysis by flow cytometry showed accumulation in the G1 phase with a concomitant reduction in S-phase entry in *MLKL*-deficient cells (**Figure 5B**). Consistently, *MLKL* knockdown increased p21 protein expression while reducing RB and CDC2 phosphorylation, assessed by western blotting (**Figure 5C**). Functionally, MLKL depletion significantly reduced cell proliferation, colony formation, and spheroid formation. Similar results were observed following treatment with the human MLKL inhibitor necrosulfonamide (NSA)(10) (**Figures 5D, 5E**, **S5D-F**). Analysis of TCGA-LIHC datasets demonstrated elevated *MLKL* expression in primary HCC tumors compared with normal liver tissue (**Figure S5G**). Immunohistochemical analysis confirmed increased MLKL protein abundance in human tumors (**Figure 5F**), and elevated MLKL expression was associated with reduced overall survival in Kaplan-Meier plotter analysis (**Figure S5H**).

**Figure 5.**
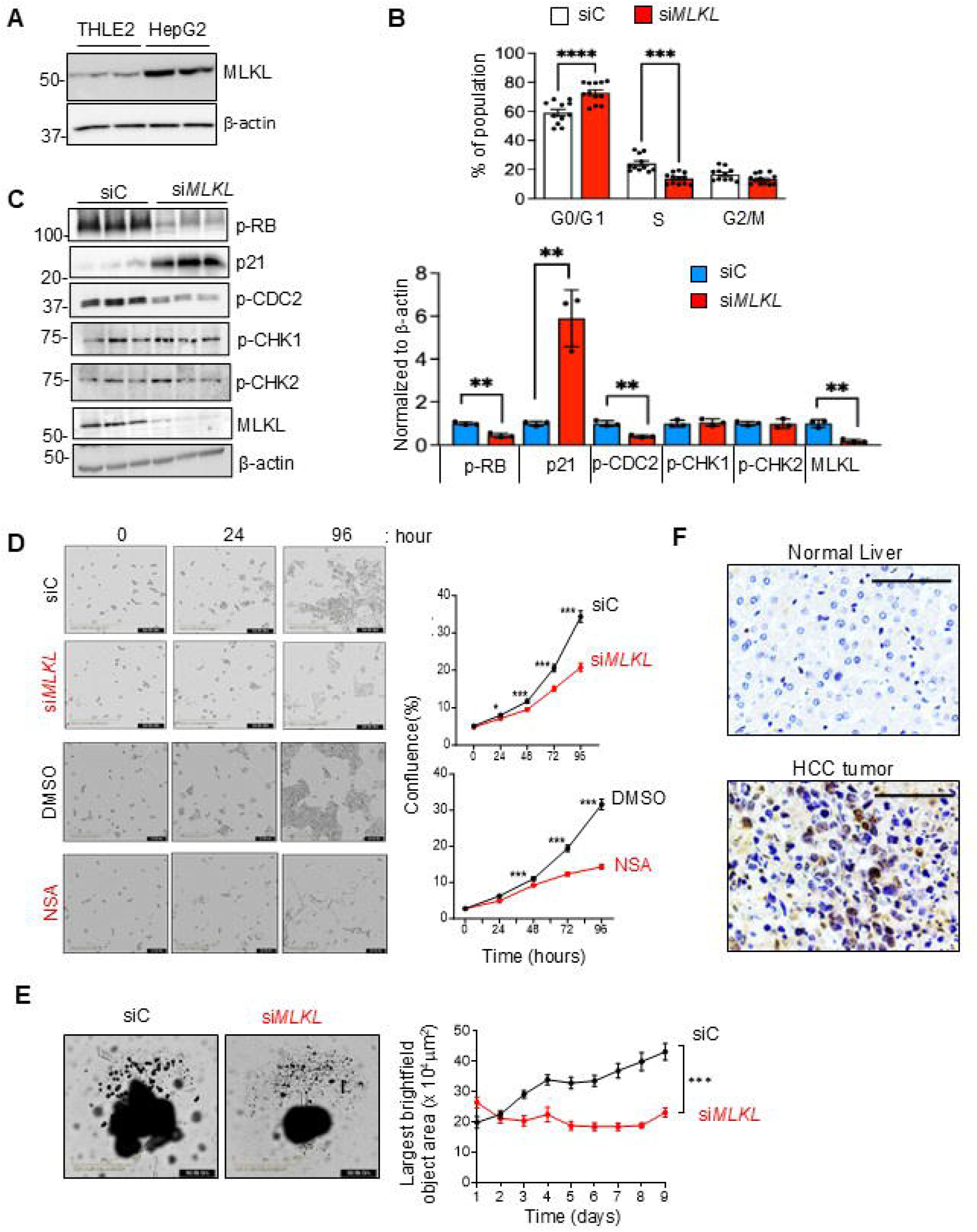
(A) Immunoblots showing MLKL and β-actin (loading control) in normal human liver cell line (THLE2) and HCC cell line (HepG2). (B-E) Data from control siRNA (siC) or *MLKL* siRNA (si*MLKL*) treated HepG2 cells: (B) Cell cycle analysis determined by flow cytometry. Graph shows the percentage of cells in the G0/G1, S, and G2/M phases. Each dot represents individual samples/group. (C) *Left*: Immunoblot showing the changes in p-RB, p21, p-CDC2, p-CHK1, p-CHK2, and MLKL. *Right:* Graphical representation of quantified blots normalized to β-actin and expressed as fold change. Each dot represents an individual experiment (n=3/group). (D) *Left*: Representative images from Incucyte live-cell imaging showing changes in cell confluence from 0-96 hours, (n=12/group). *Top*: siC or si*MLKL* cells, and *Bottom*: DMSO (vehicle) or necrosulfonamide (NSA, 5 μM) treated HepG2 cells, (n=12/group). *Right*: Proliferation curves representing cell confluency (in percentage) from time 0-96 h. siC or DMSO (black line) or si*MLKL* or NSA (red line). (E) *Left*: Representative images of spheroids formed by siControl or si*MLKL* HepG2 cells, (n=12/group). *Right*: Real-time spheroid growth monitoring curve representing spheroid area (μm^2^) over 9 days. (F) Representative IHC images of human liver tissue microarray showing MLKL expression in normal liver tissues *(top)* or HCC tumors *(bottom)*. Scale bar: 100 μm. Data represented as mean ± SEM. Two-tailed unpaired; P< 0.05. p<0.05, **p< 0.01, ***p<0.001, ****p<0.0001.

### Loss of MLKL restores the mitochondrial fusion and tumor-suppressor protein mitofusin 2 (MFN2) and enhances mitochondrial respiratory capacity

Bulk RNA sequencing of livers from mice fed WD for 15 months demonstrated distinct transcriptional profiles between WD-fed control and *Mlkl^HepKO^*mice (**Figure S6A**). Pathway enrichment analysis revealed increased oxidative phosphorylation, respiratory electron transport, mitochondrial translation, and complex I biogenesis pathways in *Mlkl^HepKO^* livers, whereas inflammatory and immune signaling pathways were reduced (**Figure 6A**). Mitochondrial DNA content and electron transport chain protein abundance were similar between genotypes (**Figures S6B, S6C**). However, transmission electron microscopy (TEM) demonstrated that WD-induced reductions in mitochondrial size were preserved in *Mlkl^HepKO^* livers, with mitochondrial area quantified using ImageJ (**Figures 6B, S6D**). Western blot analysis showed that WD increased the phospho-dynamin-related protein 1 (DRP1)/DRP1 ratio and reduced fusion protein MFN2 in control livers, indicating a shift toward mitochondrial fission and impaired mitochondrial fusion. In contrast, WD-fed *Mlkl^HepKO^* mice exhibited reduced phospho-DRP1/DRP1 and increased MFN2 abundance (**Figure 6C**). WD did not alter fission mitochondrial 1 (FIS1), or fusion proteins MFN1 and optic Atrophy 1 (OPA1), whereas hepatocyte MLKL deficiency reduced FIS1 and MFN1 levels (**Figure 6C**). In fatty acid treated AML12 hepatocytes, MLKL knockdown preserved MFN2 protein expression, reduced the p-DRP1/DRP1 ratio, and increased OPA1 expression (**Figures 6D, S6E**). *Mfn2* transcript levels were unchanged in WD-fed control and *Mlkl^HepKO^*livers but were significantly reduced in fatty acid-treated MLKL-knockdown AML12 cells (**Figure S6F**). Measurement of hepatic NADH oxidase, an overall assessment of electron transport chain activity, in isolated liver mitoplasts demonstrated impaired mitochondrial bioenergetics in WD-fed control mice, whereas this decline was attenuated in *Mlkl^HepKO^* mice (**Figure 6E**). Analysis of 4 and 8-month diet fed liver tissues showed that while WD significantly reduced MFN2 expression at both time points, the p-DRP1/DRP1 ratio was similar between WD-fed control and *Mlkl^HepKO^* mice (**Figure S6G**). Additionally, bulk RNA sequencing of livers from mice fed WD for 8 months demonstrated downregulation of pathways associated with DNA synthesis and cell cycle control of chromosomal replication in *Mlkl^HepKO^* mice (**Figures S6H-J)**.

**Figure 6.**
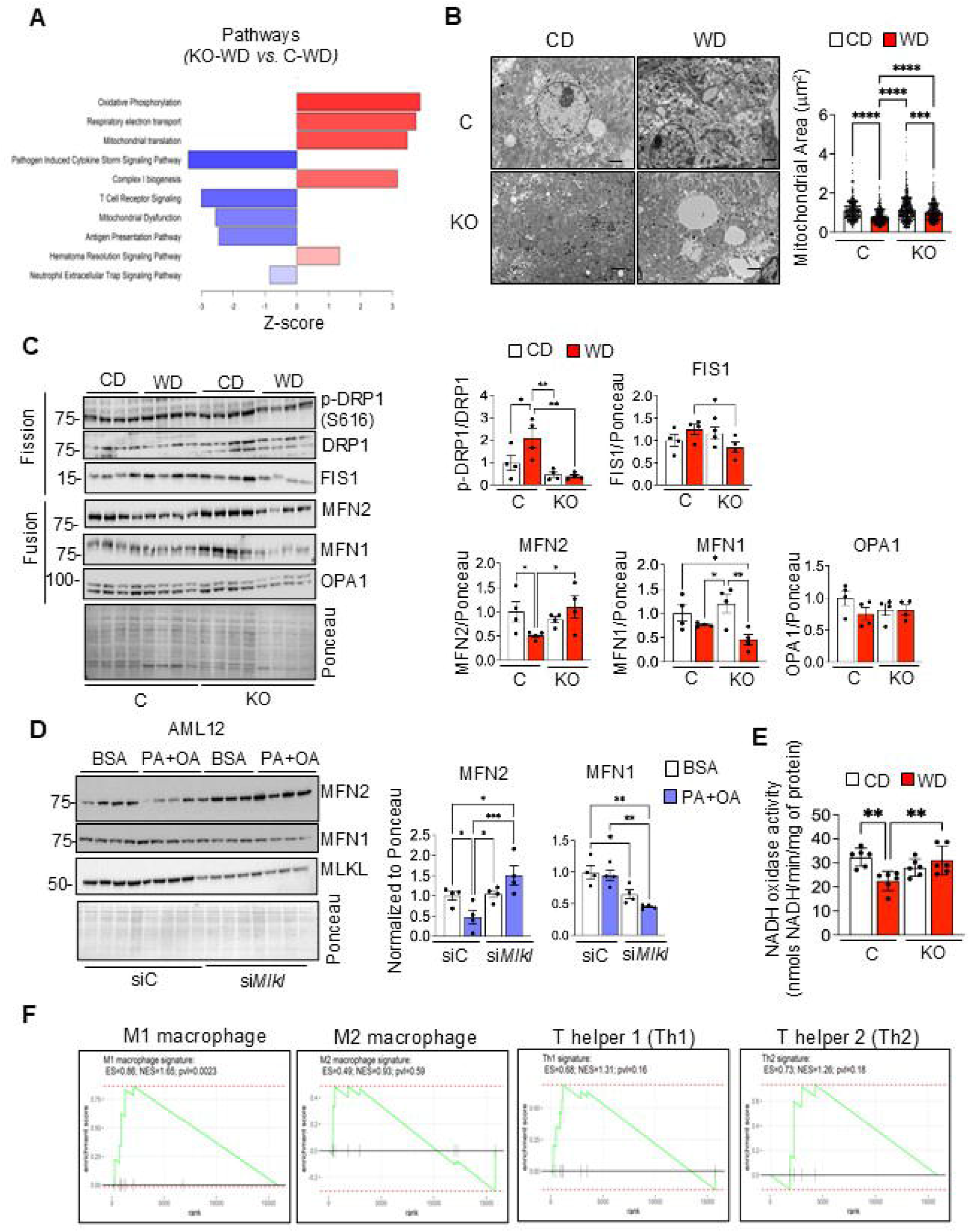
(A-C) Data from livers of control (C) or *Mlkl^HepKO^* (KO) mice fed either Control diet (CD) or Western diet (WD) for 15 months. (A) Graphical representation of canonical pathways identified by bulk RNA sequencing that were significantly upregulated (red) or suppressed (blue), (n=5 mice/group). (B) *Left:* Transmission electron microscopy images of liver tissue (n=3 mice/group; Magnification: 1000X; scale bar= 1µm). *Right:* Graphical representation of mitochondrial area (μm^2^). Each dot represents an individual mitochondrion measurement. (C) *Left*: Immunoblots showing changes in mitochondrial dynamics proteins p-DRP1 (Ser616), DRP1, FIS1, MFN2, MFN1, OPA1 and Ponceau (loading control). *Right:* Graphical representation of immunoblots as fold change normalized to Ponceau. Each dot represents an individual mouse (n=4 mice/group). (D) *Left*: Immunoblots showing MFN2, MFN1, and Ponceau (loading control) in siC and si*Mlkl* AML12 cells treated with BSA (control) or BSA-PA:OA (250:125 μM) for 48 h (n=4). *Right:* Quantification of MFN2 and MFN1 expression represented as fold change normalized to Ponceau. Each dot represents an individual experiment. (E) NADH oxidase activity in mitoplasts isolated from the livers of control (C) or *Mlkl^HepKO^* (KO) mice fed either Control diet (CD) or Western diet (WD) for 15 months. Each dot represents an individual mouse (n=6 mice/group). (F) Gene Set Enrichment Analysis (GSEA) enrichment plots evaluating: M1 or M2 macrophage signature, and T-Helper: Th1/2 signature profiles in 15 months diet fed *Mlkl^HepKO^* WD vs. Control WD. Data are represented as mean ± SEM. Statistical significance was determined by One-way ANOVA, P<0.05; *p<0.05, **p< 0.01, ***p<0.001, ****p<0.0001.

Since bulk RNA-sequencing revealed downregulation of inflammatory and immune signaling pathways in WD-fed *Mlkl^HepKO^* mice, we performed gene set enrichment analysis (GSEA), which demonstrated enrichment of M1 macrophage-associated gene signatures, with no significant enrichment of M2 macrophage-, Th1-, or Th2-associated gene signatures (**Figure 6F**). Consistently, immunohistochemical analysis of the M1-associated macrophage polarization marker iNOS (24) showed significantly increased iNOS^+^ staining in tumors from WD-fed *Mlkl^HepKO^* mice compared with WD-fed control mice (**Figures S6K)**. Analysis of hepatic inflammatory mediators revealed increased *Il1β*, *Ccl2*, *Cxcl9*, and *Cxcl10* gene expression following WD feeding, particularly at 15 months (**Figure S6L**). Hepatocyte-specific MLKL deletion had minimal effects on these responses, although *Tnf*α and *Ccl2* expression were modestly increased in *Mlkl^HepKO^* mice. Similar findings were observed after 8 months of WD feeding both at the transcript and protein levels (**Figures S6M, S6N**).

### MLKL deficiency enhances mitochondrial respiration through MFN2 and suppresses HCC cell proliferation

To establish the relevance of the MLKL-MFN2-mitochondrial axis in tumor cells and validate findings from hepatocyte models, we examined the effect of siRNA mediated *MLKL* depletion in HepG2 cells and in primary hepatocytes isolated from *Mlkl^−/−^* mice. *MLKL* knockdown increased MFN2 expression and oxygen consumption rate (OCR) in HepG2 cells (**Figures 7A, S7A**). Similar effects were observed in primary hepatocytes isolated from male, not female, *Mlkl^−/−^* mice (**Figures 7B, S7B**). Additionally, MLKL knockdown increased OCR in AML12 cells (**Figures S7C).** To determine whether MLKL regulates MFN2 through direct interaction, we assessed MLKL-MFN2 association by reciprocal co-immunoprecipitation in WD-fed liver and proximity ligation assays in fatty acid treated AML12 cells. These analyses revealed minimal association between MLKL and MFN2 (**Figures S7D-F**), suggesting that MLKL regulates MFN2 indirectly. Because *Mfn2* mRNA levels did not explain the increased MFN2 protein abundance, we examined whether MLKL regulates MFN2 stability through proteasomal degradation. In AML12 cells, MG132 treatment increased MFN2 abundance under metabolic stress conditions (**Figure 7C**), whereas in HepG2 cells, MG132 treatment restored MFN2 protein levels, supporting proteasome-dependent regulation of MFN2 turnover in both cell lines (**Figure S7H**). Next, to determine whether MFN2 is required for the metabolic consequences of MLKL deficiency, MFN2 was silenced in *MLKL*-knockdown HepG2 cells. MFN2 depletion reversed the increased OCR, reduced ECAR/glycolysis as assessed by Seahorse extracellular flux analysis, suppressed cell proliferation, and altered mitochondrial morphology, as assessed by MitoTracker Red staining (**Figures 7D-F, S7I, S7G**). Consistent with reduced glycolytic activity observed with *MLKL* knockdown, transcriptomic analysis of MLKL-deficient HepG2 cells revealed downregulation of glycolysis-associated genes (**Table S7**). Similarly, proteins involved in glycolysis elevated in WD-fed control livers were reduced in WD-fed *Mlkl^HepKO^* livers (**Figure S7J**).

**Figure 7.**
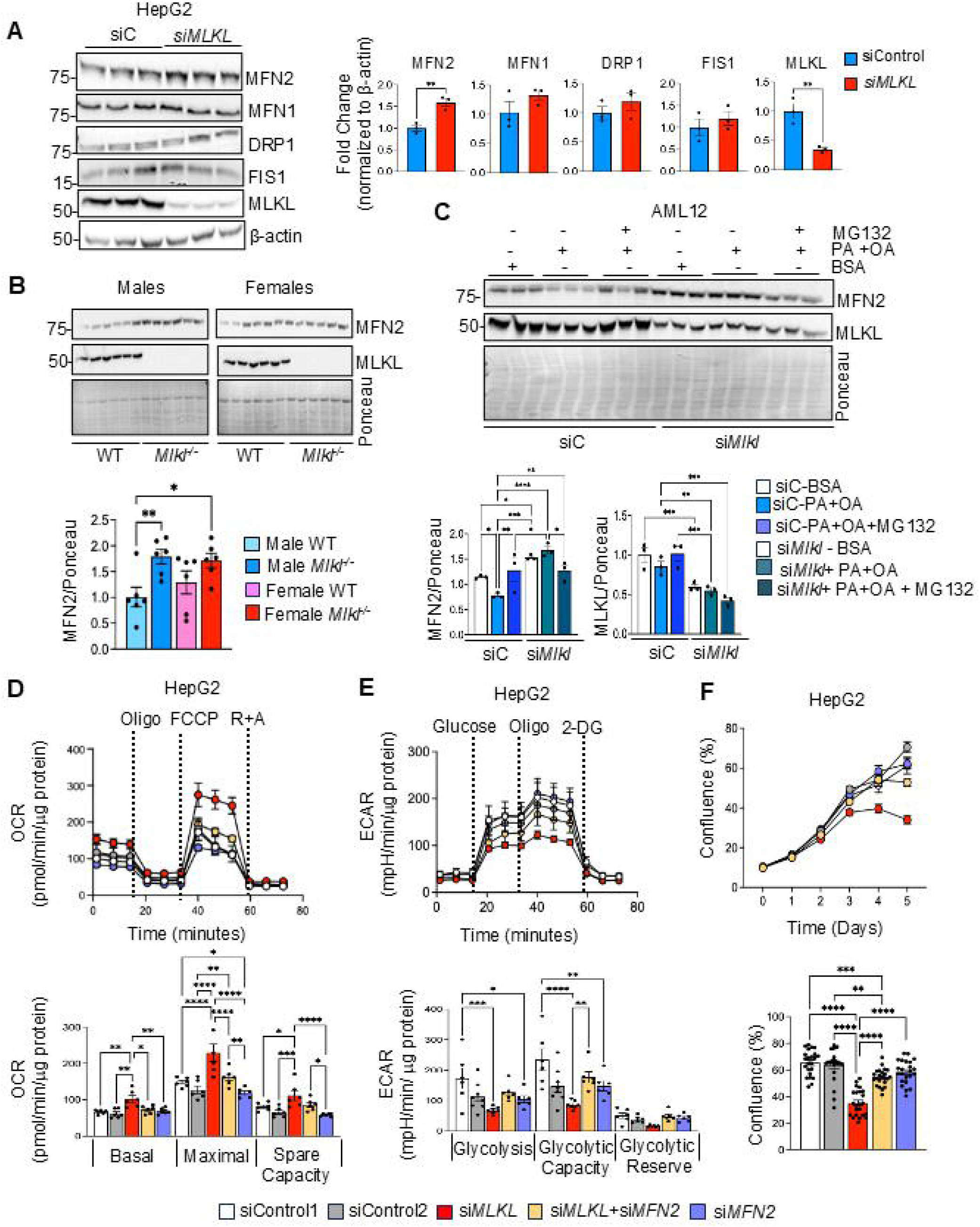
(A) *Left*: Immunoblots showing MFN2, MFN1, DRP1, FIS1 and MLKL in control (siC) or *MLKL* knockdown (si*MLKL)HepG2* cells. *Right:* Graphical representation of quantified blots as fold change normalized to β-actin. (B) *Top*: Immunoblot showing MFN2, MLKL and Ponceau in primary hepatocytes isolated from 3-month-old male or female wild type (WT) or *Mlkl* knockout (*Mlkl^−/−^*) mice, (n= 6/group). *Bottom:* Quantified MFN2 immunoblot as fold change normalized to Ponceau. Each dot represents an individual mouse. (C) *Top:* Western blots showing MFN2 and MLKL expression in siC or si*Mlkl* AML12 hepatocytes treated with BSA (375 µM) or BSA-PA:OA (250:125 µM), with (+) or without (−) 10 µM MG132 for 4 h. Ponceau staining served as the loading control. *Bottom:* Quantified MFN2 and MLKL immunoblots as fold change normalized to Ponceau. Each dot represents an individual experiment (n= 3/group). (D-F) Data from HepG2 cells treated with control siRNA [(siControl 1=50 nM or siControl 2=100 nM)], *MLKL* siRNA (50 nM), *MFN2* siRNA (50 nM) or a combination of *MLKL* (50 nM) and *MFN2* (50 nM) siRNA for 48 h. (D) *Top:* Representative oxygen consumption rate (OCR) curve, with sequential addition of oligomycin, FCCP, and rotenone/antimycin A (R+A). *Bottom:* Bar graph summarizing the quantified respiratory parameters. (E) *Top:* Extracellular acidification rate (ECAR) analysis in treated cells at basal level and after sequential addition of glucose, oligomycin and 2-deoxy glucose. *Bottom:* Bar graph summarizing the quantified glycolytic parameters. (F) *Top*: Proliferation curves from Incucyte live-cell imaging showing changes in cell confluence over 5 days in treated cells. *Bottom*: Quantification of percentage cell confluency in treated cells, (D-F: Each dot represents an individual sample). Data points marked with asterisks (*) indicate statistically significant differences comparing the mean of each group with that of every other group. (A) Two-tailed unpaired t-test. (B-F) One-way ANOVA, P<0.05. *p<0.05, **p< 0.01, ***p<0.001, ****p<0.0001.

### MLKL localizes to nuclear and mitochondrial compartments and represents a potential therapeutic target in HCC

Canonical MLKL function involves plasma membrane translocation to execute necroptosis; however, its localization under non-necroptotic conditions remains poorly defined. Confocal microscopy demonstrated MLKL localization in both the nucleus and cytoplasm of HepG2 and AML12 cells, with partial co-localization with mitochondrial markers (**Figures 8A, 8B**). Subcellular fractionation followed by immunoblotting also showed presence of MLKL in nuclear and mitochondrial fractions (**Figure S8A**). In tumors from WD-fed control mice, MLKL localized predominantly to the nucleus and cytosol (**Figure 8C**), supporting a compartment-specific role for MLKL in hepatocytes and HCC cells beyond its classical membrane-associated necroptotic function.

**Figure 8.**
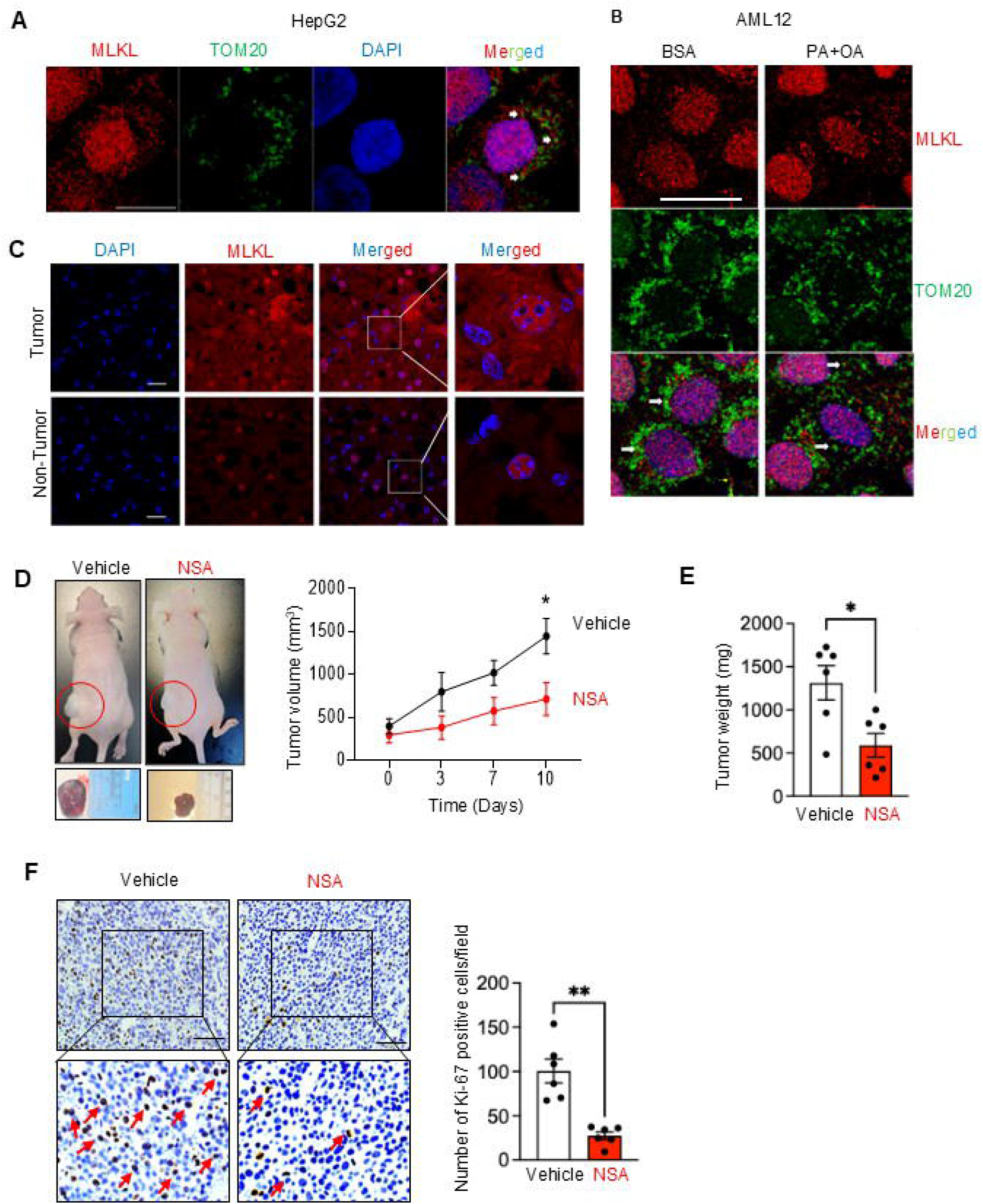
(A-B) Representative immunofluorescence images showing MLKL (red), TOM-20 (green) and DAPI (blue) staining in HepG2 cells (Scale bar: 10 μm) (A), and AML12 cells treated with BSA (control) or BSA-PA:OA (Scale bar: 25 μm) (B). (C) Immunofluorescence images showing MLKL protein stain (red) in the livers of control (C) or *Mlkl^HepKO^*(KO) mice fed WD for 15 months (Scale Bar: 25 μm). (D) *Left:* Representative gross morphology images of xenograft mice *(top)* and tumors *(bottom)* showing tumor size (red circle) differences between vehicle or NSA treated groups (n=6 mice/group). *Right:* Line graph tracking the tumor volume (mm^3^) over a 10-day monitoring period. (E) Quantitative evaluation of tumor weight (mg) at the experimental end point comparing vehicle control group against NSA treatment group, (n=6 mice/group). (F) *Left:* Representative IHC staining images of proliferation marker Ki-67 in tumor sections from vehicle and NSA-treated mice (n=6 mice/group) (Scale bar: 60 μm). *Right:* Graphical representation of the Ki-67 positive proliferating cells per field. Each dot represents an individual mouse. Data represented as mean ± SEM. Two-tailed unpaired; P< 0.05 *p<0.05, **p< 0.01.

Finally, to evaluate the therapeutic potential of MLKL inhibition, nude mice bearing subcutaneous HepG2 xenografts were treated with NSA or vehicle for 10 days. NSA significantly reduced tumor volume and tumor growth compared with vehicle-treated controls (**Figures 8D-F**). NSA treatment did not affect body weight, serum ALT and AST levels, or hematological parameters (**Figure S8B**, **Table S9**). These findings identify MLKL as a potential therapeutic target in HCC.

## DISCUSSION

The present study identifies a hepatocyte-intrinsic, non-canonical role for MLKL in MASLD-associated HCC. Despite marked WD-induced MLKL expression, canonical necroptosis was undetectable. Hepatocyte-specific MLKL deletion did not alter liver injury, fibrosis, or inflammation but increased hepatic lipid accumulation while markedly reducing HCC incidence, multiplicity, tumor size, and stemness and proliferation markers. These findings identify hepatocyte MLKL as a tumor promoter under metabolic stress, although a role in early tumorigenesis cannot be excluded. Elevated MLKL expression in human HCC correlated with poorer survival, whereas genetic or pharmacological MLKL inhibition suppressed HCC cell proliferation and clonogenicity. Together, these findings establish MLKL as a hepatocyte-intrinsic, non-necroptotic promoter of metabolic liver cancer, independent of MASLD severity.

Our findings support a non-necroptotic role for hepatocyte MLKL in WD-induced MASLD-HCC progression. Canonical necroptosis, assessed by p-MLKL and MLKL oligomerization, was undetectable throughout disease progression. These findings are consistent with a recent study using a streptozotocin/high-fat, high-sugar diet-induced MASLD-HCC mouse model, in which robust HCC developed despite undetectable hepatic necroptosis, whereas parenchymal-specific *Mlkl* knockout mice exhibited reduced HCC incidence (25). In our study, RIPK3 overexpression restored p-MLKL expression in hepatocytes, suggesting that the absence of necroptosis during WD feeding reflects the lack of RIPK3, which is epigenetically silenced in hepatocytes and HepG2 cells (16, 17). This is consistent with our recent finding that primary hepatocytes from aged mice lack detectable RIPK3 and are resistant to necroptosis induction (26). Supporting a necroptosis-independent role, MLKL promotes hepatocarcinogenesis by inhibiting AMPK-dependent autophagy, impairing mitophagy, and suppressing oxidative stress-induced parthanatos, as demonstrated in DEN-induced HCC, hepatic ischemia-reperfusion injury, and RIPK3-deficient HCC models (17, 27, 28). Together, these findings highlight emerging non-necroptotic functions of MLKL in liver disease, including regulation of autophagy and vesicular trafficking (29, 30).

A notable finding of our study is the dissociation between MASLD severity and HCC development. Despite comparable or greater steatosis, fibrosis, and liver injury in WD-fed *Mlkl^HepKO^* mice, hepatocyte-specific MLKL deficiency markedly reduced HCC incidence, supporting a hepatocyte-intrinsic role for MLKL in hepatocarcinogenesis. Similar dissociation has been reported by Imerzoukene et al. (25) and in our prior studies using whole-body *Mlkl^−/−^* mice (13). Notably, hepatocyte MLKL deficiency increased triglyceride accumulation and lipid droplets while reducing HCC, a phenotype recapitulated in hepatocyte models. Although *Mlkl* deletion promotes lipid accumulation in macrophages (31), systemic or adipocyte-specific *Mlkl* deletion reduces steatosis (32, 33), suggesting that systemic effects may mask hepatocyte-specific MLKL functions. Despite increased lipid storage, MLKL deficiency reduced lipotoxic ceramide and diacylglycerol species, preserved mitochondrial respiration, and enhanced lipid droplet–mitochondria interactions. Thus, lipid composition and metabolic consequences, rather than total lipid burden, influence WD-driven hepatocarcinogenesis.

Transcriptomic, functional, and biochemical analyses identified mitochondrial metabolism as a major pathway regulated by MLKL. RNA sequencing revealed enrichment of oxidative phosphorylation and mitochondrial respiratory programs in 15-month WD-fed *Mlkl^HepKO^* livers. Consistently, MLKL deficiency preserved mitochondrial morphology, maintained hepatic NADH oxidase activity, and enhanced mitochondrial respiration in hepatocyte models that is consistent with prior reports (26, 34). MLKL deficiency also reduced glycolysis, suggesting a shift away from a glycolytic, tumor-supportive metabolic state. Given the role of aerobic glycolysis in HCC progression and tumor growth (35, 36), these findings support a metabolically less permissive environment for hepatocarcinogenesis in WD-fed *Mlkl^HepKO^* mice.

MFN2 appears to be a key mediator of MLKL-dependent metabolic reprogramming. In HepG2 cells, MFN2 depletion abolished the increased mitochondrial respiration, reduced glycolysis, and decreased proliferation associated with MLKL deficiency, indicating that MFN2 is required for these effects. Consistent with its tumor-suppressive role in HCC, MFN2 promotes mitochondrial fusion and oxidative phosphorylation while limiting glycolysis and is frequently downregulated in HCC correlating with poor clinical outcomes (37–40). In our model, WD feeding reduced MFN2 protein levels, whereas hepatocyte-specific MLKL deficiency preserved MFN2 expression. Proteasome inhibition restored MFN2 levels in AML12 and HepG2 cells, suggesting a role of MLKL in MFN2 degradation under lipotoxic stress. Since MFN2 is regulated by the ubiquitin-proteasome system (41), MLKL may control MFN2 stability through this pathway. Although a direct MLKL-MFN2 interaction was not detected, future studies are needed to identify upstream regulators of MFN2 turnover.

Beyond mitochondrial metabolism, we identified a previously unrecognized role for MLKL in regulating cell cycle progression. MLKL localized to the nucleus in tumor and non-tumor hepatocytes, suggesting a functional nuclear pool. Consistent with a non-necroptotic role, MLKL nuclear translocation has been reported in lung cancer cells through a ROS-dependent, RIPK3- and oligomerization-independent mechanism (42). Although MLKL contains a nuclear localization signal and RIPK3 can promote nuclear translocation during necroptosis (43–45), the mechanisms regulating RIPK3-independent nuclear localization remain unclear and may involve non-canonical phosphorylation sites (43, 44). Functionally, MLKL deficiency reduced cell cycle progression and proliferation, suggesting that nuclear MLKL may promote tumor-supportive programs. In addition, our findings identify mitochondrial MLKL in hepatocytes and HCC cells and implicate it in MFN2-dependent mitochondrial metabolism and tumor progression, consistent with emerging evidence for mitochondrial functions of non-necroptotic MLKL (26, 45).

Temporal analyses support a tumor-promoting role for hepatocyte MLKL in MASLD-associated HCC. Although WD feeding reduced MFN2 expression at 4-8 months, mitochondrial fission (p-DRP1/DRP1) and hepatocyte proliferation (Ki-67) remained comparable between genotypes in non-tumor liver, indicating that early MFN2 loss occurs independently of altered fission or proliferation. Bulk RNA sequencing of WD-fed *Mlkl^HepKO^* livers showed reduced DNA synthesis and cell cycle pathways at 8 months and enhanced oxidative phosphorylation programs at 15 months, consistent with preserved MFN2 and mitochondrial function. Reduced Ki-67 positivity was observed specifically in *Mlkl^HepKO^*tumors but not adjacent non-tumor liver. Thus, hepatocyte MLKL appears to promote tumor progression by establishing a tumor-permissive metabolic state rather than initiating early metabolic stress responses. Although hepatocyte MLKL deficiency did not broadly suppress WD-induced inflammatory mediators and modestly increased *Tnf*α and *Ccl2* expression, the M1-associated transcriptional signature suggests that MLKL deficiency promotes an M1-enriched anti-tumor microenvironment rather than quantitatively altering hepatic inflammation. This is consistent with prior work showing that MLKL loss in RIPK3-deficient HCC enhances antitumor immunity and improves checkpoint blockade responses (17). In HCC, iNOS+ macrophages are recognized as tumor suppressive through NO/ROS-mediated cytotoxicity and apoptosis (46). The mechanisms by which hepatocyte MLKL deficiency promotes M1 polarization remain to be defined.

Because HCC incidence is higher in men, our long-term WD-HCC studies were performed in male mice (47). Although our prior work showed higher HCC incidence in males, whole-body *Mlkl* deficiency protected both sexes, supporting a sex-independent role for MLKL in hepatocarcinogenesis (13). However, our short-term studies revealed that MLKL deletion preserved MFN2 and mitochondrial respiration in male but not female hepatocytes, suggesting sex-dependent regulation of this pathway. Whether hepatocyte MLKL deletion protects female mice from WD-induced HCC remains to be determined. Human relevance is supported by increased MLKL expression in HCC and its association with poor survival in TCGA-LIHC and patient samples. The human MLKL inhibitor NSA recapitulated MLKL knockdown effects in HepG2 cells, reducing proliferation, clonogenic growth, and xenograft growth without overt toxicity. These findings support MLKL as a potential biomarker and therapeutic target in HCC, while further studies are needed to define sex specificity and therapeutic safety.

In summary, hepatocyte MLKL functions predominantly as a tumor promoter in MASLD-associated hepatocarcinogenesis through a RIPK3-independent mechanism, while a role in early tumorigenic events cannot be excluded. MLKL deficiency preserves MFN2, enhances mitochondrial respiration, remodels lipid metabolism, limits cell cycle progression and stemness-associated programs, and promotes an antitumorigenic immune microenvironment. Genetic deletion and pharmacological inhibition of MLKL attenuate HCC development, supporting the MLKL-MFN2 axis as a potential therapeutic target in metabolic liver cancer.

Limitations of the study: The long-term WD-HCC cohort included only male mice, and future studies in females are needed to define sex-dependent mechanisms. While MFN2 was supported as a downstream mediator, the mechanism of MLKL-dependent MFN2 proteasomal degradation remains unclear, and contributions from non-parenchymal cells was not explored. Future studies should define MLKL-MFN2 regulation, clarify nuclear MLKL functions, and determine whether MFN2 restoration rescues tumor phenotypes.

## Supporting information

Supplementary methodology

## AUTHOR CONTRIBUTIONS

Phoebe Ohene-Marfo, Sabira Mohammed, Chao Jiang, Shylesh Bhaskaran, Kara Kneuper, Megan John, Julianne N. Hoang and Albert Tran performed the experiments, analyzed the data, and helped with manuscript preparation. Jonathan D Wren, Constantin Georgescu and Tae Gyu Oh analyzed the lipidomics and bulk RNA sequencing data. Willard M Freeman and Kevin Pham performed and analyzed data for the mitochondrial DNA copy number assay. Michael Kinter performed and analyzed the targeted mitochondrial proteomics data. Randal J May performed immunostaining in liver and tumor tissues. Surendra Shukla assisted with xenograft mouse model studies. Satoshi Matsuzaki, Bo Hagy, and Kenneth Humphries performed mitochondrial functional and oxidative stress marker assays and Chinthalapally V Rao and Courtney Houchen edited the manuscript. Sathyaseelan S. Deepa designed the study, supervised the research, wrote, reviewed and edited the manuscript.

## LIST OF ABBREVIATIONS

AAV8: Adeno-associated virus serotype 8
ACC: Acetyl-CoA carboxylase
ACSL4: Acyl-CoA synthetase long-chain family member 4
AFP: Alpha-fetoprotein
ALT: Alanine transaminase
AST: Aspartate aminotransferase
AML12: Alpha mouse liver 12
AMPK: AMP-activated protein kinase
BAX: Bcl-2-associated X protein
BSA: Bovine Serum Albumin
CCL: C-C motif chemokine ligand
CXCL: C-X-C motif chemokine ligand
CD: Control diet
CD31/36/206: Cluster of differentiation 31/36/206
Cdc2: Cyclin-Dependent Kinase 1
Cidec: Cell Death Inducing DFFA Like Effector C
Clec4f: C-type lectin domain family 4 member F
CPT1A: Carnitine palmitoyltransferase 1a
DAMPs: Damage-associated molecular patterns
DCLK1: Doublecortin Like Kinase 1
DEN: Diethylnitrosamine
2-DG: 2-deoxy-D-glucose
DMEM: Dulbecco’s modified eagle medium
DMSO: Dimethyl sulfoxide
DRP1: Dynamin-related protein 1
ECAR: Extracellular acidification rate
EMEM: Eagle’s Minimum Essential Medium
ETC: Electron transport chain
FATP: Fatty Acid Transport Protein
FBS: Fetal Bovine Serum
FCCP: Carbonyl cyanide p-trifluoromethoxyphenyl hydrazine
FDR: False discovery rate
FIS1: Mitochondrial fission 1 protein
GPC-3: Glypican-3
GPX4: Glutathione peroxidase 4
GSDMD: Gasdermin D
GSEA: Gene Set Enrichment Analysis
GSR: Glutathione Reductase
HCC: Hepatocellular carcinoma
4-HNE: 4-hydroxynonenal
HPRT: Hypoxanthine phosphoribosyltransferase 1
IL: Interleukin
iNOS: Inducible nitric oxide synthase
LDH: Lactate dehydrogenase
LIPE: Hormone-sensitive lipase E
MASH: Metabolic dysfunction-associated steatohepatitis
MASLD: Metabolic dysfunction-associated steatotic liver disease
Mfn: Mitofusin
MLKL: Mixed lineage kinase domain-like protein
MOPS: 3-(N-morpholino) propanesulfonic acid
NADH: Nicotinamide Adenine Dinucleotide + Hydrogen
NPC: Non-parenchymal cells
NSA: Necrosulfonamide
OA: Oleic acid
OCR: Oxygen Consumption Rate
OCT4: Octamer-binding transcription factor 4
OHP: Hydroxyproline
OPA1: Optic Atrophy 1
PA: Palmitic acid
PARP: Poly(ADP-ribose) polymerase
PLIN2: Perilipin 2
Pnpla: Patatin-like phospholipase domain containing 2
Pol E: Polymerase E
PPAR: Peroxisome proliferator-activated receptor
PSR: Picro-Sirius Red Stain
PTM: Post-Translational Modifications
Rb: Retinoblastoma
RIPK: Receptor interacting serine/threonine kinase
Scd1: Stearoyl-CoA desaturase 1
SOD2: Superoxide Dismutase 2
TBG: Thyroxin Binding Globulin
TCGA-LIHC: The Cancer Genome Atlas Liver Hepatocellular Carcinoma
TEM: Transmission electron microscopy
TNFα: Tumor necrosis factor alph
TXN1: Thioredoxin 1
TXNRD1: Thioredoxin Reductase 1
VDAC: Voltage-Dependent Anion Channel
WD: Western diet

## ACKNOWLEDGMENTS

The authors would like to acknowledge Prof. James Murphy, Walter and Eliza Hall Institute of Medical Research, Australia for generously providing the *Mlkl^flox/flox^* mice model used in the study. The H&E staining and PSR staining service provided by the Stephenson Cancer Center tissue pathology core, Incucyte live cell imaging service, XF analyzer provided by the Stephenson Cancer functional genomics core are supported partly by the National Institute of General Medical Sciences Grant P30GM154635 and National Cancer Institute Grant P30CA225520 of the National Institutes of Health. The flow cytometry and confocal microscopy imaging service was provided by the Institutional Research Core Facility The authors thank imaging core facility at Oklahoma Medical Research Foundation for the electron microscopy imaging.

