## Supplementary figures for "Hepatocyte MLKL Drives Obesity-Driven Hepatocellular Carcinoma Progression via Mitochondrial Dysfunction Independent of Necroptosis in MASLD"

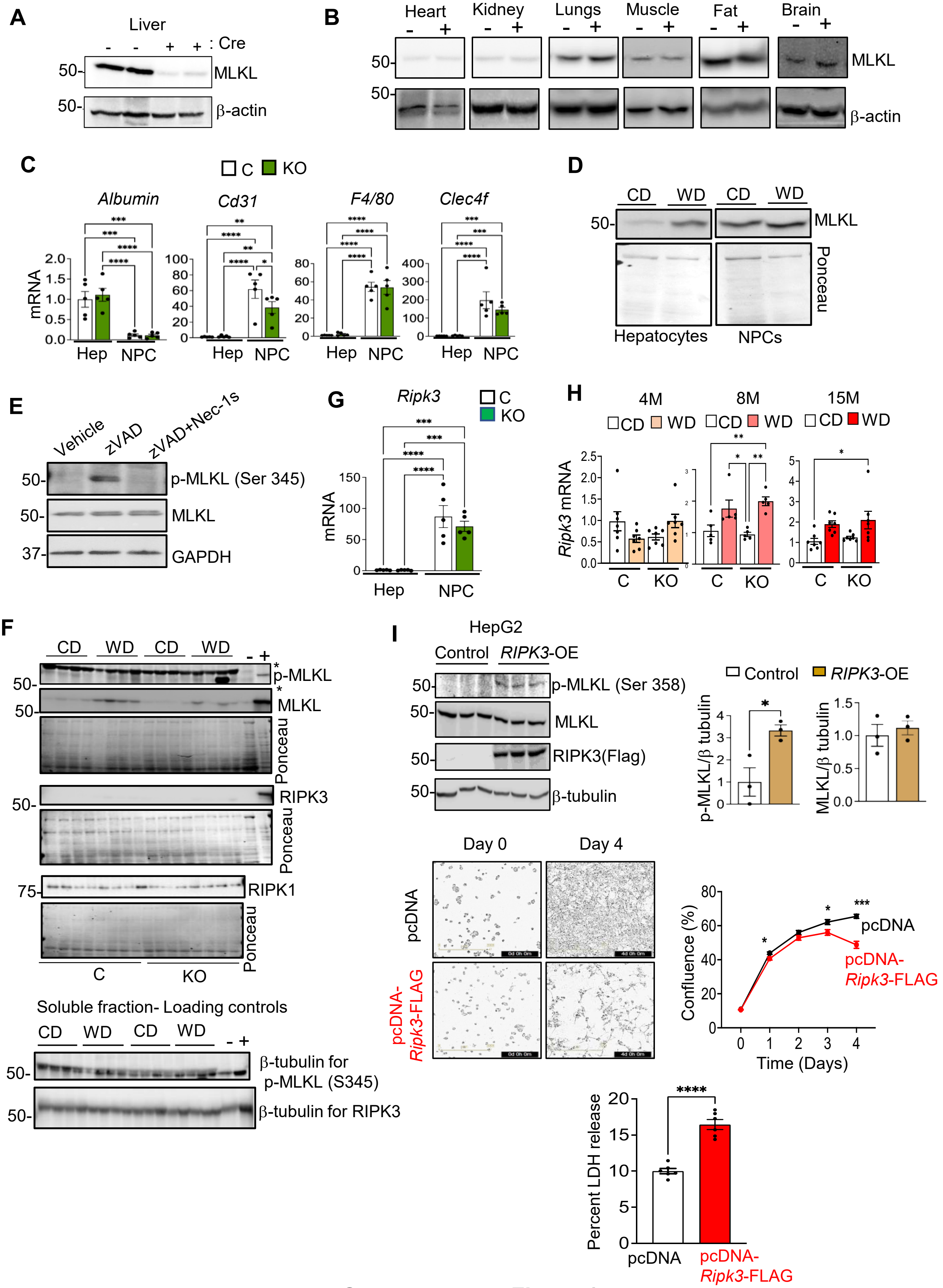


**Supplementary Figure 1**

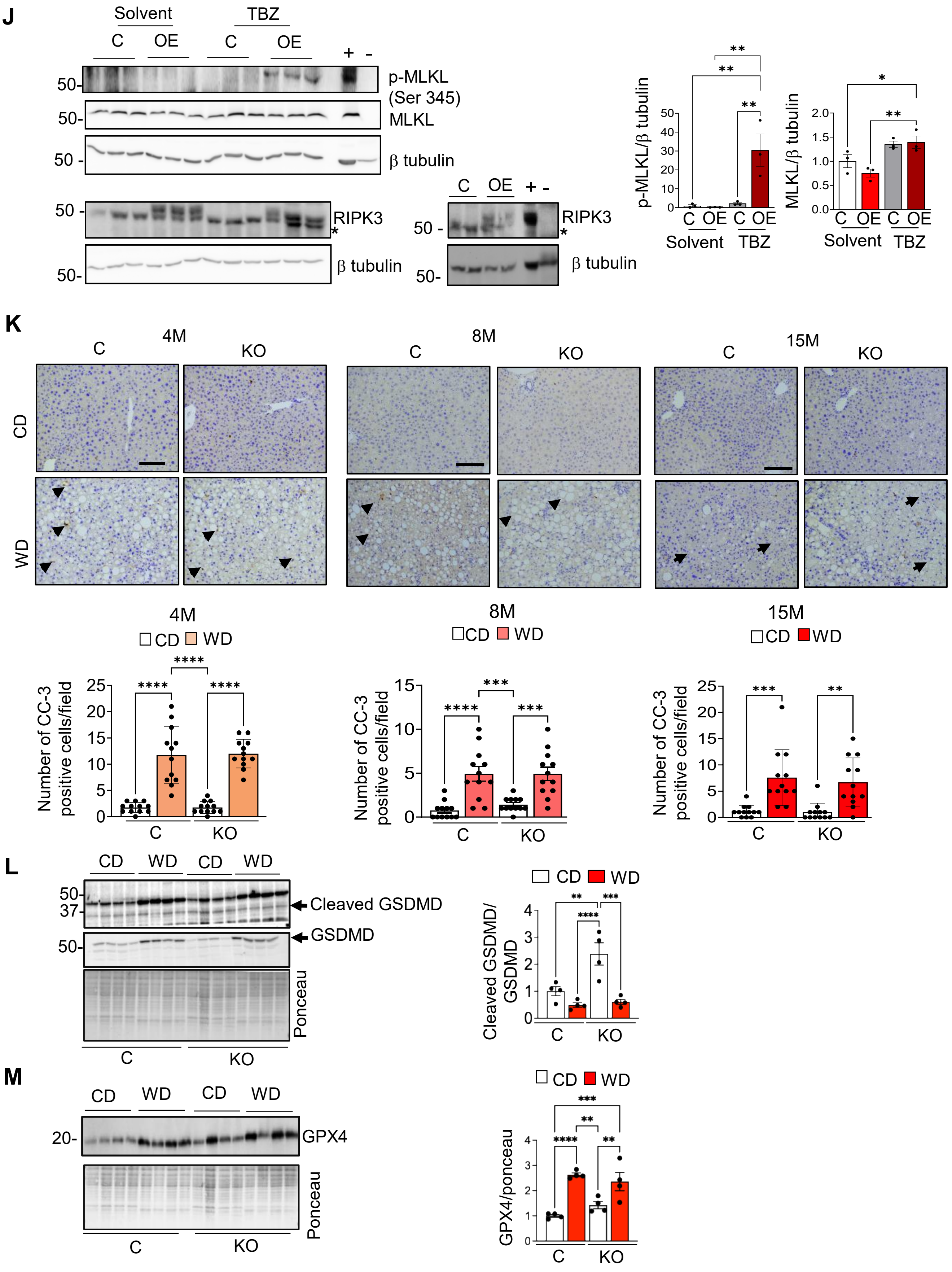


**Supplementary Figure 1 (Continued)**

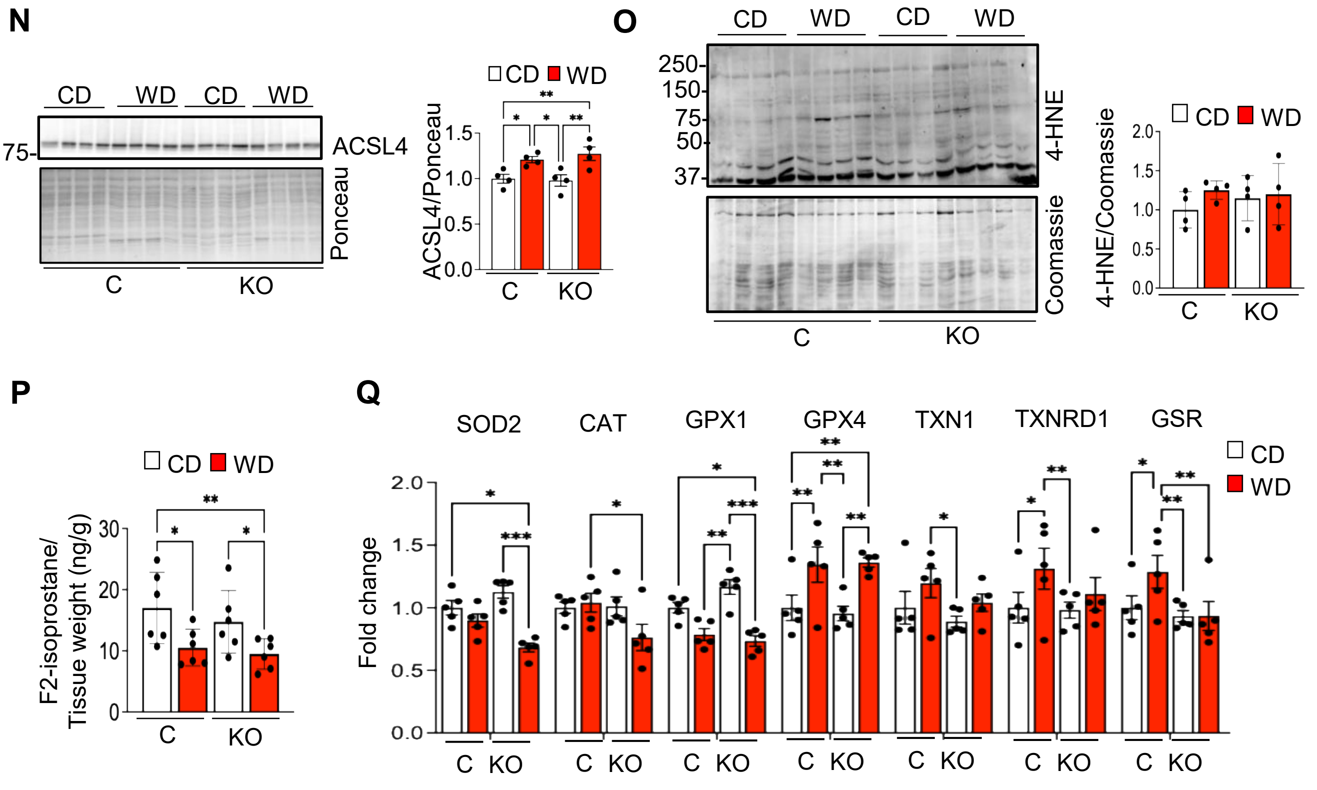


**Supplementary Figure 1 (Continued)**

**A**

| Mean Lipid droplet size (μm²) | | | |
|---|---|---|---|
| **Group** | **4M** | **8M** | **15M** |
| C-WD | 36.98 | 51.58 | 46.44 |
| KO-WD | 50.979 | 66.118 | 24.67 |

**B**

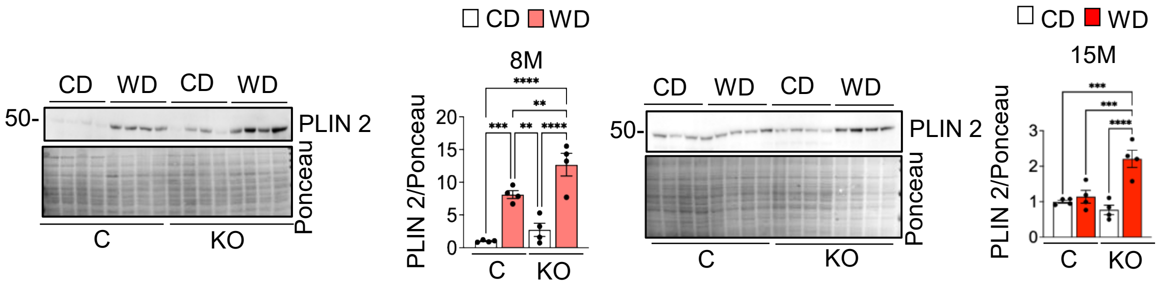


**C**

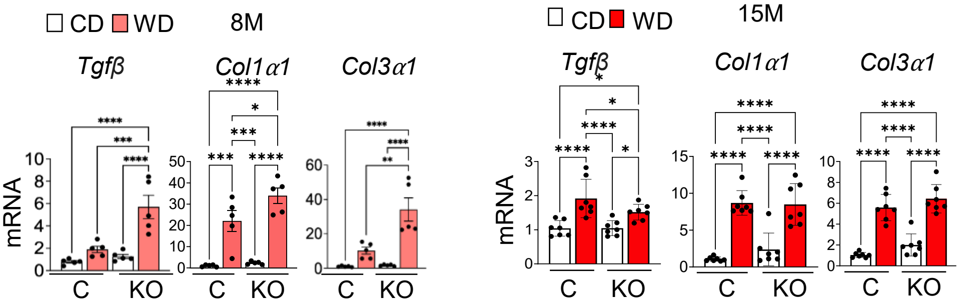


**Supplementary Figure 2**

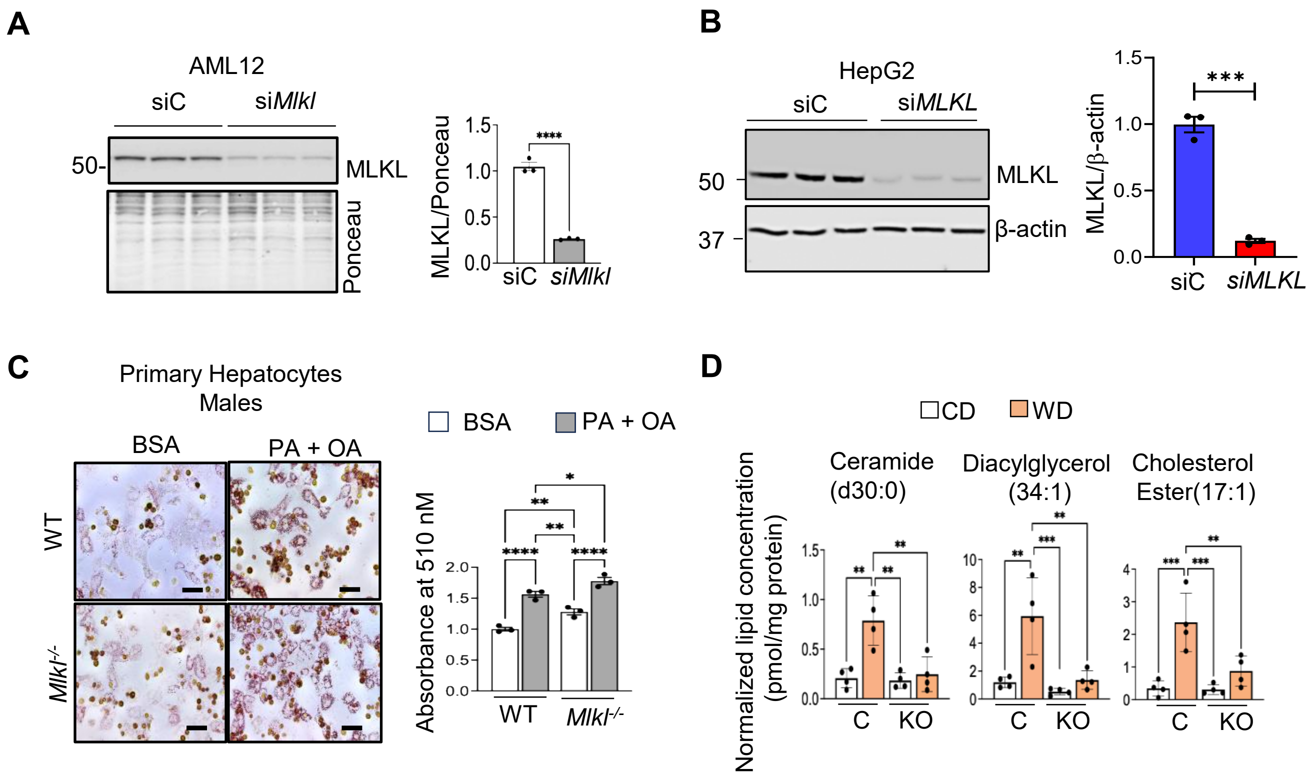


**Supplementary Figure 3**

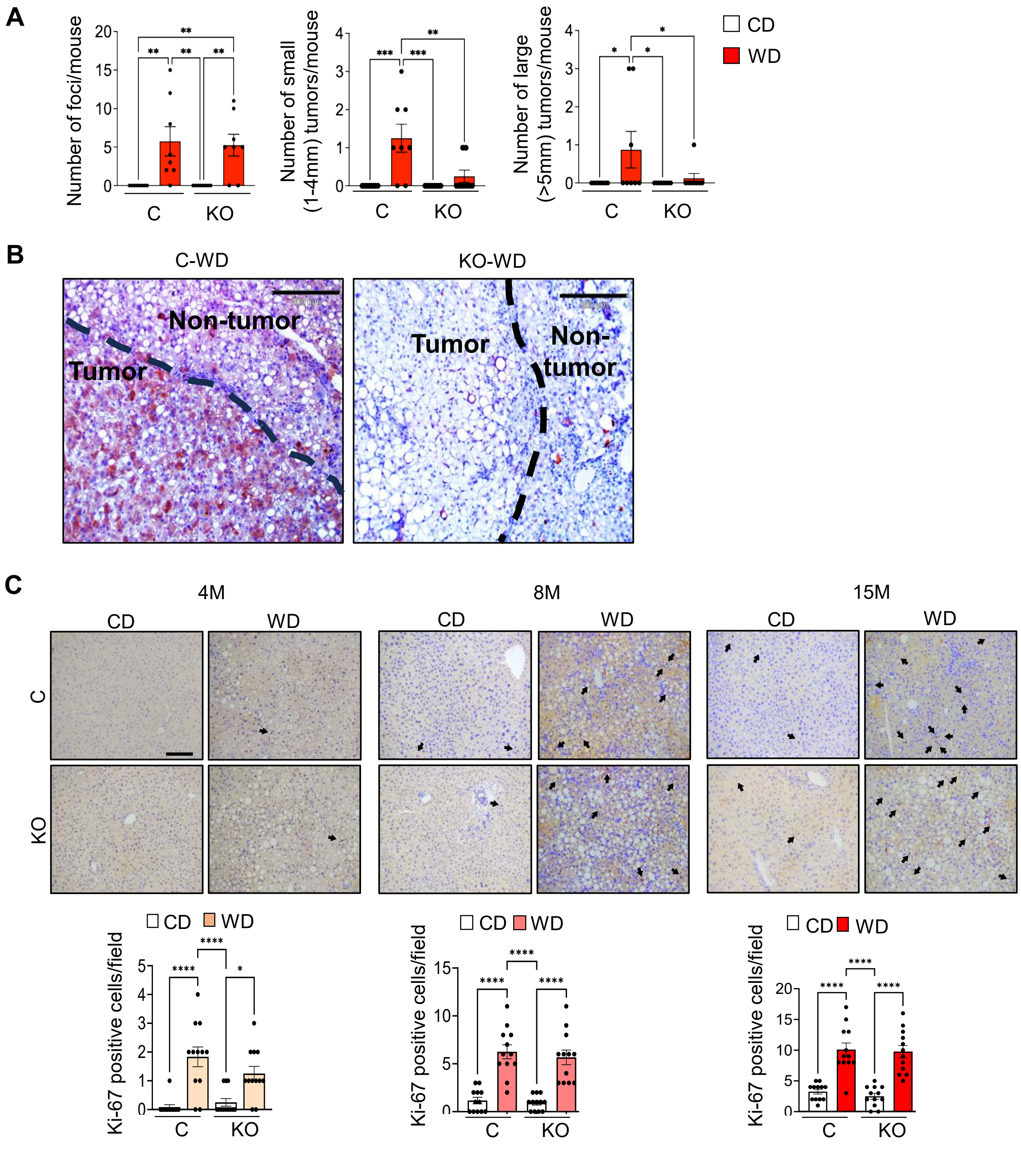


**Supplementary Figure 4**

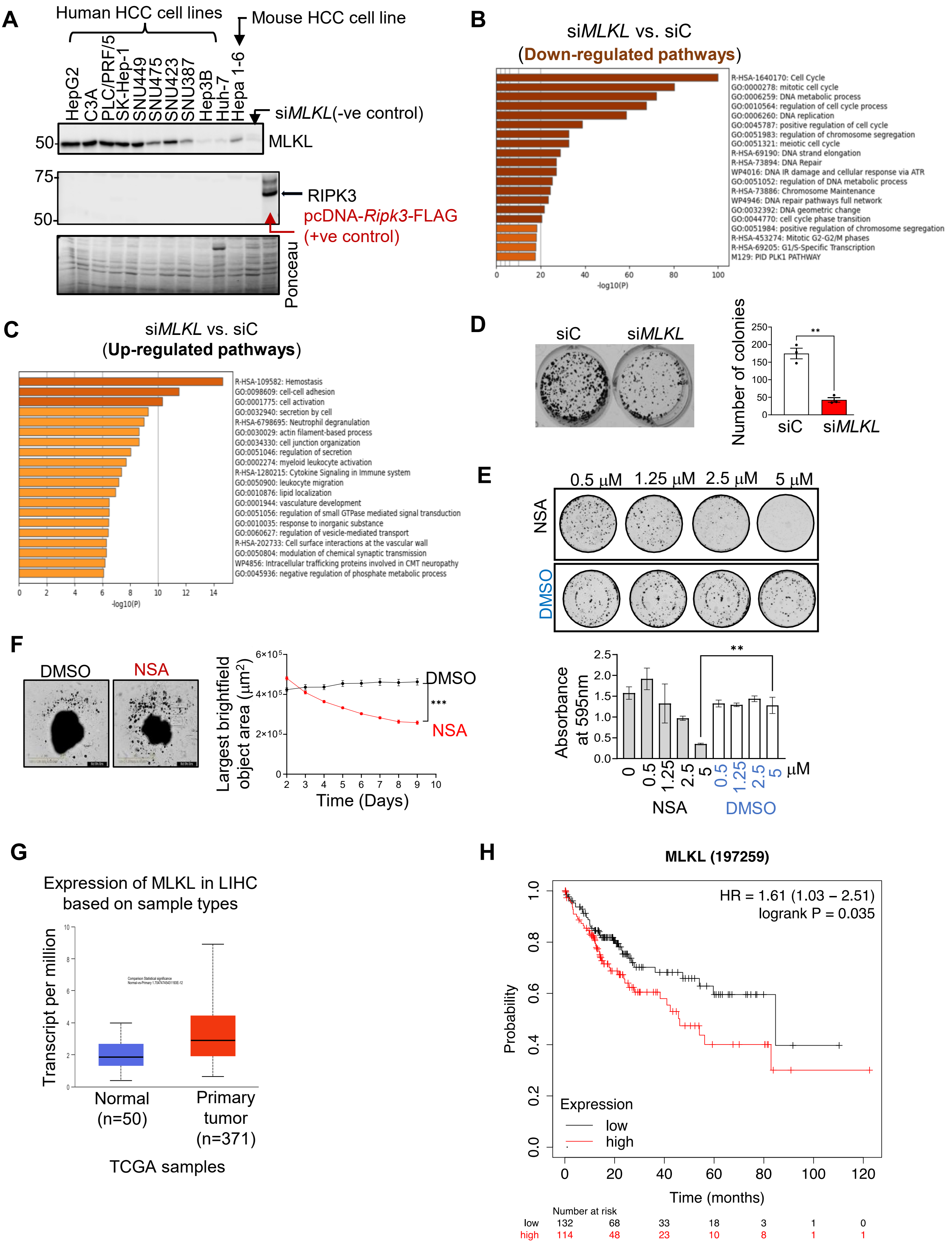


**Supplementary Figure 5**

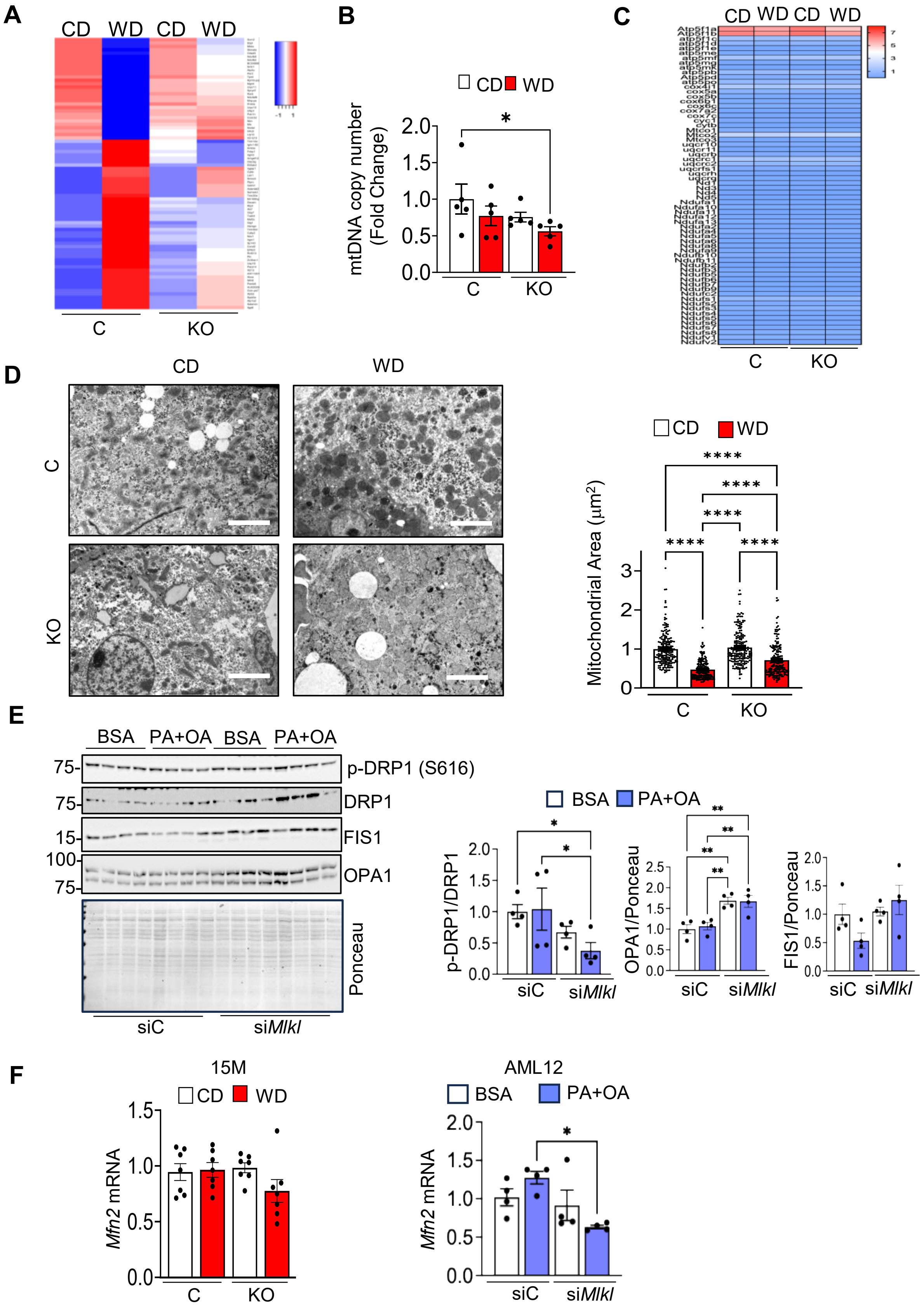


**Supplementary Figure 6**

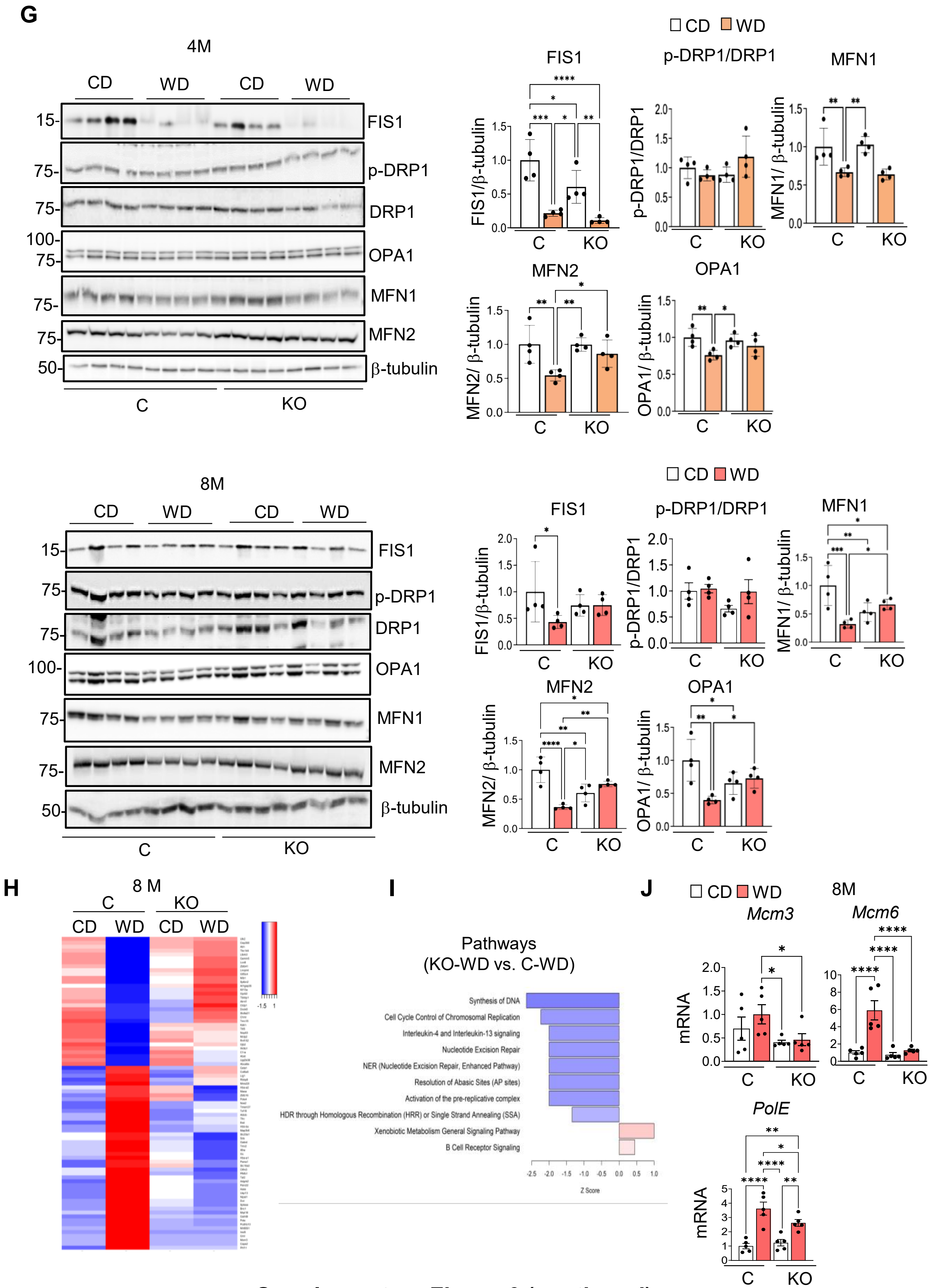


**Supplementary Figure 6 (continued)**

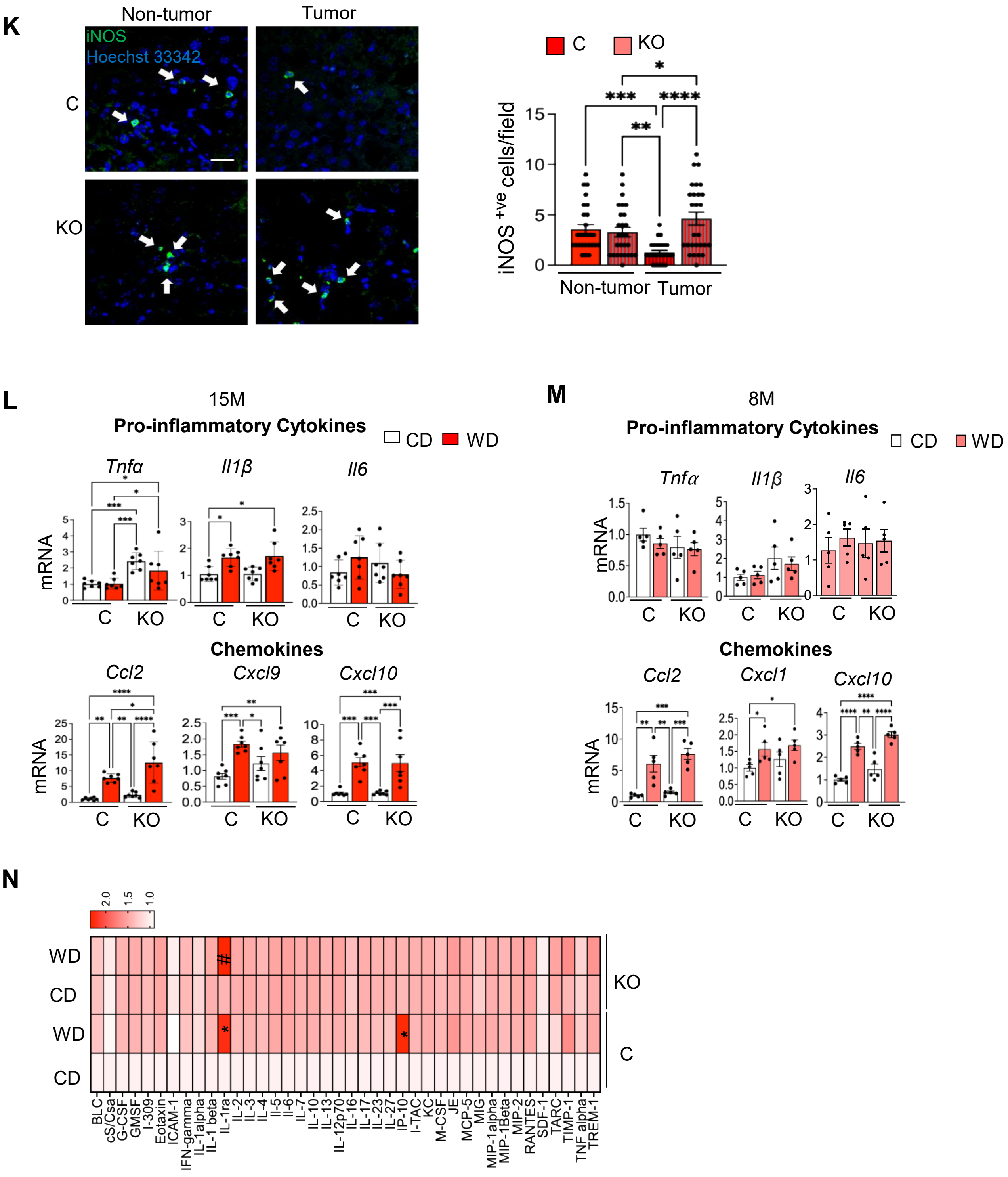


**Supplementary Figure 6 (continued)**

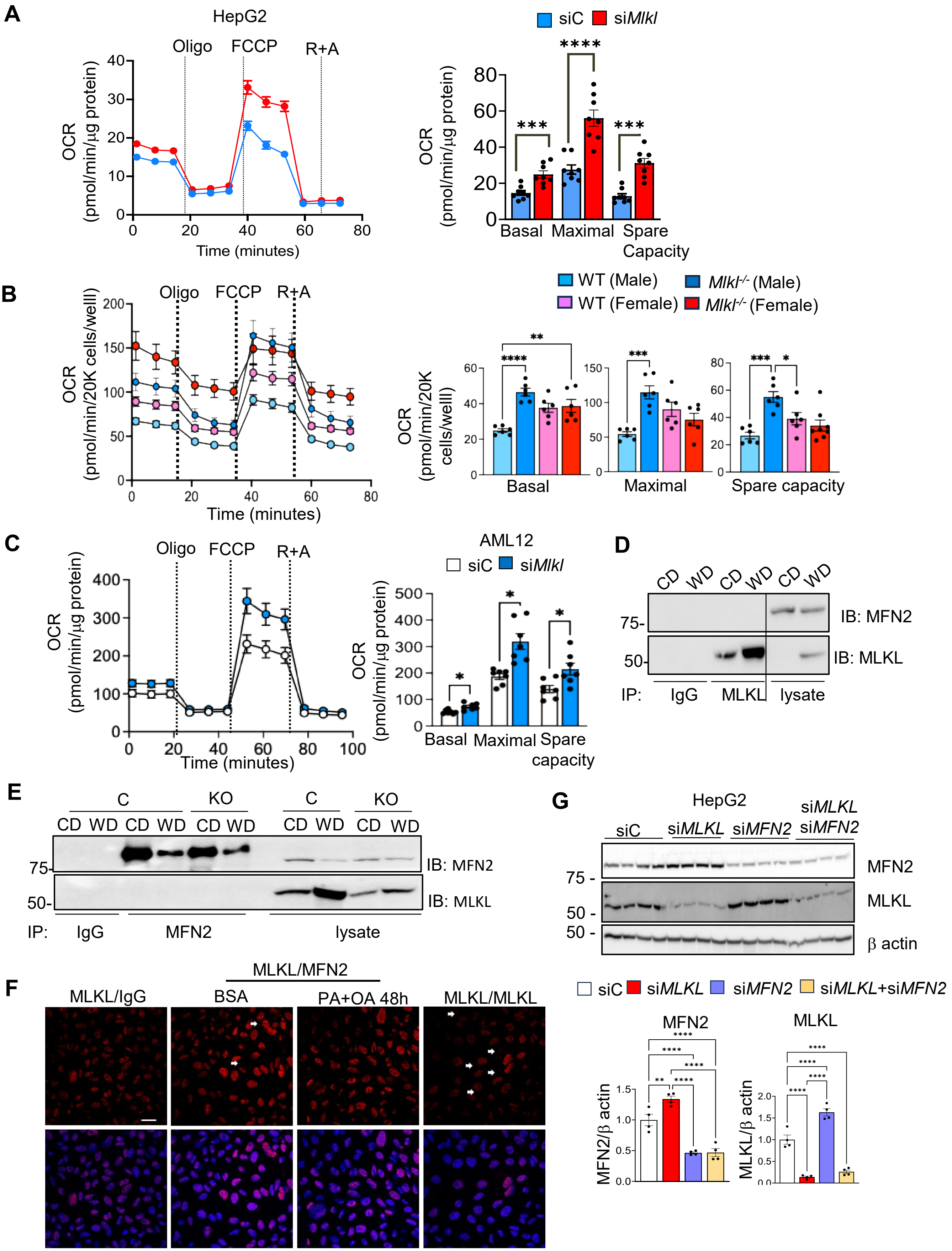


**Supplementary Figure 7**

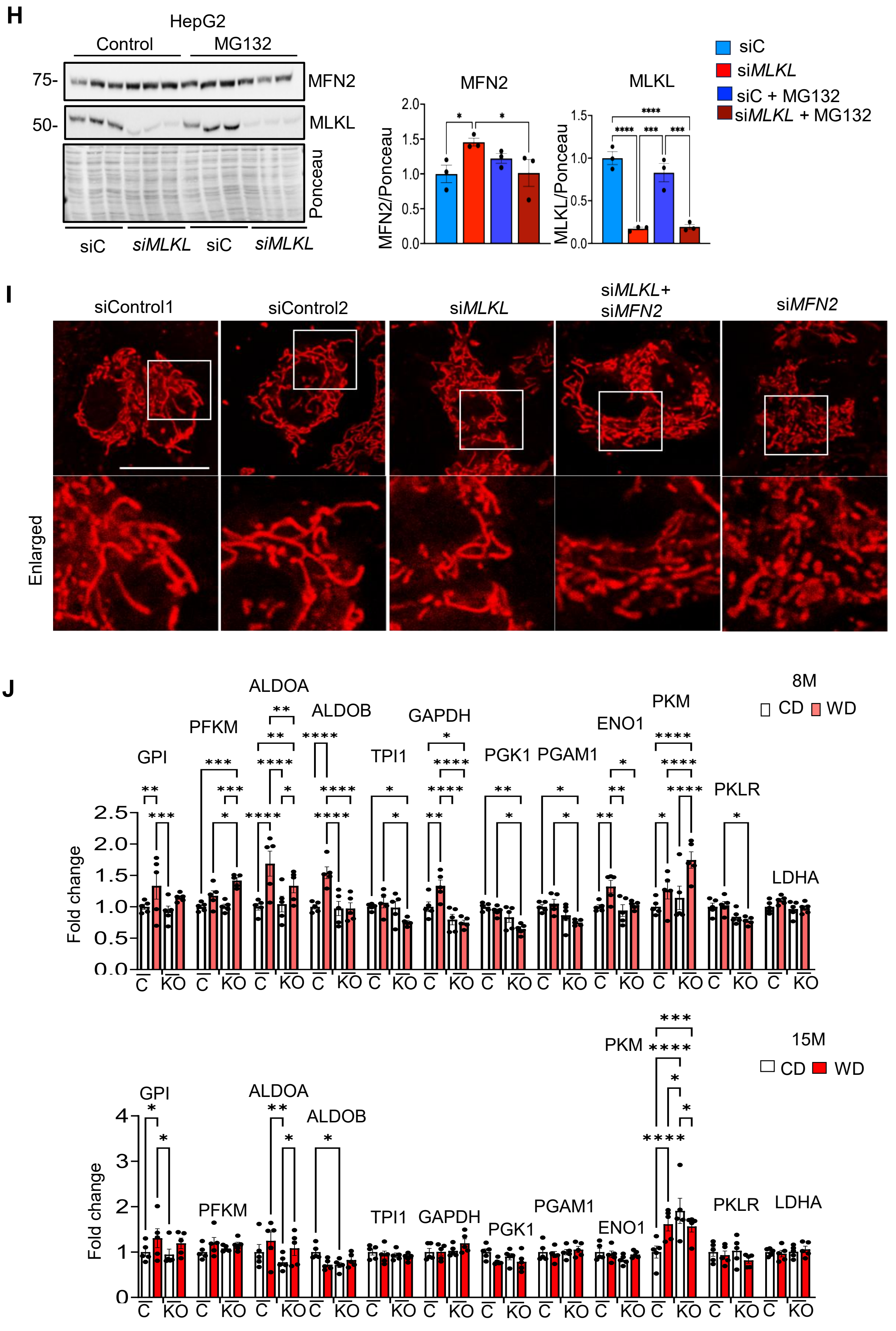


**Supplementary Figure 7 (continued)**

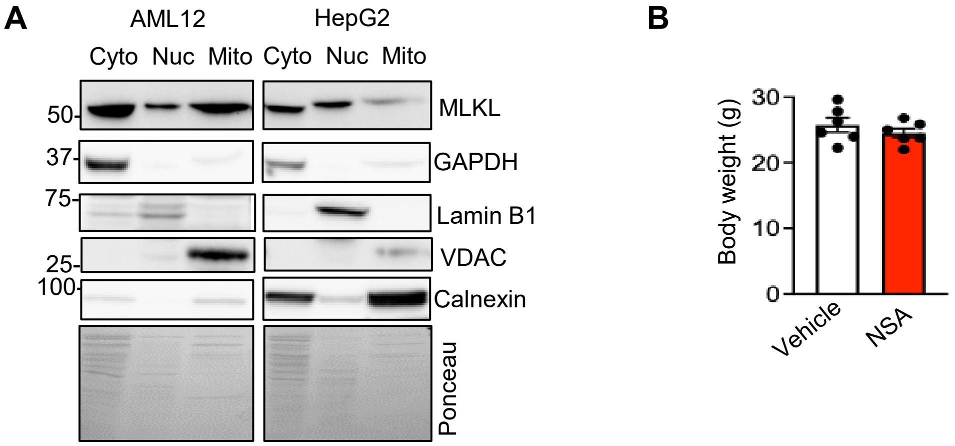


**Supplementary Figure 8**
