## Supplementary figure legends for "Hepatocyte MLKL Drives Obesity-Driven Hepatocellular Carcinoma Progression via Mitochondrial Dysfunction Independent of Necroptosis in MASLD"

**Figure S1.** Immunoblots showing the expression of MLKL and β-actin (loading control) two weeks after AAV8-TBG-Null (Control, Cre-) and AAV8-TBG-Cre (*Mlkl*$^{HepKO}$, Cre+) injection in the (A) whole liver extracts, and (B) various tissues from the two experimental groups to validate the *Mlkl*$^{HepKO}$ mouse model. (C) Transcript levels of *Albumin*, *F4/80*, *Clec4f*, and *Cd31* in isolated hepatocytes and non-parenchymal cell (NPC) fractions of control (C, white bars) or *Mlkl*$^{HepKO}$ mice (KO, green bars), normalized to β-microglobulin and expressed as fold change (n=5 mice/group). (D) Immunoblot showing the expression of MLKL with Ponceau staining as loading control in isolated hepatocytes and NPC of control mice fed either Control diet (CD) or Western diet (WD) for 4 months. (E) Western blots showing the expression of p-MLKL(Ser 345), MLKL and GAPDH in BV2 murine microglial cells treated with either vehicle (DMSO), pan caspase inhibitor, zVAD-fmk (20 μM) or zVAD + necroptosis inhibitor, Necrostatin-1s (20 μM) for 24 h. (F) Immunoblots showing expression of p-MLKL, MLKL, RIPK3, RIPK1 and Ponceau (loading control) from the insoluble fraction homogenates of the liver tissues of control (C) and *Mlkl*$^{HepKO}$ (KO) mice fed either Control (CD) or Western diet (WD) for 15 months, (n=4 mice/group). Positive controls: BV2 cells treated with pan caspase inhibitor, zVAD for p-MLKL, and *RIPK3*-overexpressing HepG2 cells for RIPK3 blots. Negative controls: Liver lysates from *Ripk3*$^{-/-}$ or *Mlkl*$^{-/-}$ mice for RIPK3 or p-MLKL blots. (G-H) The mRNA levels of *Ripk3* in: (G) isolated hepatocytes and non-parenchymal cell (NPC) fractions of control (C, white bars) or *Mlkl*$^{HepKO}$ mice (KO, green bars), normalized to β-microglobulin and expressed as fold change (n=5 mice/group), (H) Whole liver tissues of control (CD) or *Mlkl*$^{HepKO}$ (KO) mice fed Control diet (CD) or Western diet (WD) for 4-, 8- or 15-months, normalized to β-microglobulin (n=7, 5, and 7 respectively) and expressed as fold change. (I) *Top:* Western blots showing p-MLKL(S358), MLKL, RIPK3 and β-tubulin (loading control) in HepG2-*RIPK3* over expressing cells (RIPK3-OE) compared to control HepG2 cells *(left)*. Quantified immunoblots of p-MLKL and MLKL normalized to β-tubulin and expressed as fold change, (n=3) *(right)*. *Middle:* Representative images from Incucyte live-cell imaging showing changes in cell confluence over 4 days in HepG2 control (pcDNA) or *RIPK3* over expressing cells (pcDNA-*RIPK3*-FLAG) *(left)*. Proliferation curves representing cell confluency (in percentage) for 4 days *(right)*. *Bottom*: Extracellular lactate dehydrogenase (LDH) activity expressed as mean percentage of total LDH release as a measure of plasma membrane integrity.

(J) *Left:* Representative Western blot analysis p-MLKL(Ser345), MLKL, RIPK3, and β-tubulin (loading control) in control(C) and *Ripk3*-overexpressing AML12 cells (OE) treated with either solvent (DMSO) or a necroptosis-inducing cocktail (TBZ). Positive (+) control: necroptotic BV2 cell lysate for p-MLKL, and *RIPK3*-overexpressing HepG2 cell lysate for RIPK3 blots. Negative (-) controls: liver lysates from $Ripk3^{-/-}$ or $Mlkl^{-/-}$ for RIPK3 or p-MLKL blots respectively. Asterix (*) against blots indicate non-specific bands. *Right:* Quantitative of p-MLKL and MLKL blots normalized to β-tubulin and expressed as fold change. (K) *Top:* Representative IHC images for cleaved caspase 3 (CC3) staining in control (CD) or $Mlkl^{HepKO}$ (KO) mice fed Control diet (CD) or Western diet (WD) for 4, 8 or 15 months. Black arrows represent positive staining for CC3 (Scale Bar: 150 μm). *Bottom:* Quantified CC3 positive cells per field. Each dot represents an individual image taken. (n=3/group). (L-O) *Left:* Immunoblots showing the hepatic expression of : (L) Gasdermin D and Cleaved Gasdermin D, (M) GPX4 (N) ACSL4, and (O) 4-HNE protein adducts in the livers of control (C) and $Mlkl^{HepKO}$ (KO) mice fed either Control diet (CD) or Western diet (WD) for 15 months (n=4 mice/group). Ponceau or Coomassie stain (for 4-HNE) served as loading control. *Right:* Quantified immunoblots showing Cleaved GSDMD/GSDMD ratio, changes in protein expression of GPX4 or ACSL4 normalized to ponceau, and 4-HNE normalized to Coomassie, all expressed as fold change. (P) F2-isoprostane analysis, (n=6 mice/group). (Q) Proteomics analysis of antioxidant proteins in the livers of control (C) or $Mlkl^{HepKO}$ (KO) mice fed Control (CD) or Western diet (WD) for 15 months (n= 5 mice/group). Data represented as mean±SEM. Data points marked with asterisks (*) indicate statistically significant differences comparing the mean of each group with that of every other group. One-way or Two-way ANOVA (Q), P< 0.05; *p<0.05, **p< 0.01, ***p<0.001, ****p<0.0001.

**Figure S2**. Data from liver tissues of control (C) and $Mlkl^{HepKO}$ (KO) mice fed Control diet (CD) or Western diet (WD). (A) Table showing mean droplet size differences in $\mu m^2$ in the livers of control WD (C-WD) or $Mlkl^{HepKO}$ WD (KO-WD) mice after 4-, 8- or 15-month diet-feeding (n=3 per group). (B) Immunoblots showing the expression of PLIN2 and Ponceau stain (loading control) in liver tissues of mice after 8 months (*left*) or 15 months (*right*) diet feeding. Graphical representation of quantified PLIN2 blot normalized to Ponceau and expressed as fold change (n=4 mice/group). (C) Transcript levels of markers of fibrosis *Tgfβ*, *Col1α1* and *Col3α1* (normalized to β-microglobulin and expressed as fold change) in the livers of mice after 8-months (*left*) and 15-months (*right*) diet feeding, (n=5 or 7 mice/group for 8- or 15- month diet feeding respectively).

Data represented as mean ±SEM. Data points marked with asterisks (*) indicate statistically significant differences comparing the mean of each group with that of every other group. One-way ANOVA $p < 0.05$. *$p<0.05$, **$p< 0.01$, ***$p<0.001$, ****$p<0.0001$.

**Figure S3.** (A) *Left*: Immunoblot showing the expression of MLKL and Ponceau as loading control in control (siC) or *Mlkl* knockdown (si*Mlkl*) AML12 murine hepatocytes (n=3 per group). *Right:* Graphical representation of quantified MLKL for siC (white bar) or si*Mlkl* (grey bar), normalized to ponceau and expressed as fold change . (B) *Left*: Immunoblots of MLKL and β-actin (loading control) in siC or si*MLKL* HepG2 cells (n=3/group). *Right*: Graphical representation of quantified blot normalized to β-actin and expressed as fold change; siC (blue) or si*MLKL* (red). (C) *Left:* Oil red O staining in primary hepatocytes isolated from the livers of male wild type (WT) or *Mlkl* knockout ($Mlkl^{-/-}$) mice. Primary hepatocytes were treated with BSA (750 µM) or a mixture of BSA-conjugated palmitic acid (PA) and oleic acid (OA) (BSA-PA:OA, 250:500 µM) for 48 h (20X objective, Scale bar=100 µm). *Right:* Graphical representation of quantified total Oil red O extracted from primary cells and expressed as fold change (n=3/group). (D) Lipidomic quantification of lipid species: Ceramide (d30:0), Diacylglycerol (34:1) or Cholesterol ester (17:1) in the livers of control (C) and $Mlkl^{HepKO}$ (KO) mice fed either Control diet (CD) or Western diet (WD) for 4 months (n=4 mice/group). Total lipid extracts were analyzed using LC-MS/MS. Data are expressed as concentration in pmol/mg protein. Data represented as mean ± SEM. Data points marked with asterisks (*) indicate statistically significant differences comparing the mean of each group with that of every other group. (A) Two-tailed unpaired t-test, (B-D) One-way ANOVA $p < 0.05$; *$p<0.05$, **$p< 0.01$, ***$p<0.001$, ****$p<0.0001$.

**Figure S4**. (A) Number of foci/ mouse and sizes of tumors: small tumors (1-4 mm); large tumors (>5mm) from the livers of control (C) or $Mlkl^{HepKO}$ (KO) mice fed either Control diet (CD) or Western diet (WD) for 15 months. (B) Representative immunohistochemistry staining (IHC) images for Glypican 3 (GPC3) staining in tumors isolated from control WD (C-WD) vs. $Mlkl^{HepKO}$ WD (KO-WD) diet fed mice (n=3/group). Dark red color in the tumor-marked region indicates positive staining for GPC3, and a dotted line separates the tumor from the non-tumor region. (C) *Top:* Ki-67 IHC in livers of control (C) or $Mlkl^{HepKO}$ (KO) mice fed Control diet (CD) or Western diet (WD) for 4-, 8-, or 15-months. The arrows represent positive staining for Ki-67 (n=3 mice/group). *Bottom:* Quantified Ki-67 positive cells per field. Data represented as mean ± SEM. Data points marked with asterisks (*) indicate statistically significant differences comparing the mean of each group with that of every other group. One-way ANOVA $p < 0.05$. *$p<0.05$, **$p< 0.01$, ***$p<0.001$, ****$p<0.0001$.

**Figure S5.** (A) Immunoblots showing expression of MLKL and RIPK3 in various human and mouse HCC cell lines. Positive (+ve) control: RIPK3-overexpressing HepG2 cell lysates (pcDNA-*RIPK3*-FLAG) for RIPK3. Negative Control: MLKL knockdown (si*MLKL*) HepG2 cell lysate and liver lysates from *Ripk3*$^{-/-}$ mice for MLKL or RIPK3 blots respectively. (B-C) Schematic representation of pathways significantly: (B) downregulated and (C) upregulated obtained from RNA seq analysis in MLKL knockdown (si*MLKL*) compared to control (siC) HepG2 cells. (D) *Left*: Representative images of colony formation in control (siC) or *MLKL* knockdown (si*MLKL*) HepG2 cells . *Right*: Graphical representation of the number of colonies developed after colony formation assay. (E) *Top*: Representative images of colony formation in HepG2 cells treated with various concentrations (0-5 μM) of DMSO (control) or NSA (MLKL inhibition). *Bottom:* Spectrophotometric absorbance quantification of cell viability at 595 nm, (n=3/group). (F) *Left:* Representative images from the real-time spheroid growth monitoring of control (DMSO) or NSA treated HepG2 spheroids for 10 days, performed using the Incucyte live-cell imaging and analysis system. *Right:* Real-time spheroid growth monitoring curve representing spheroid area (in $\mu m^2$) over 10 days [control (black line) or *MLKL* knockdown (red line)] (n=3/group). (G) Expression of *MLKL* gene in either normal (non-cancer livers, blue bar) or primary tumor (HCC, red bar) obtained from the TCGA-LIHC dataset. (H) Kaplan-Meier curve showing the survival probability of liver cancer patients with either low (black line) or high (red line) MLKL expression level. Data represented as mean ± SEM. Data points marked with asterisks (*) indicate statistically significant differences comparing the mean of each group with that of every other group. (E) One-way ANOVA. (D &F) Two-tailed unpaired t-test; $p< 0.05$. **$p< 0.01$, ***$p<0.001$.

**Figure S6.** (A-C) Data from livers of control (C) or *Mlkl*$^{HepKO}$ (KO) mice fed either Control diet (CD) or Western diet (WD) for 15 months (n=5 mice/group). (A) Heat map showing differential gene expression; Red and blue colors represent genes that are significantly upregulated or downregulated. (B) Mitochondrial DNA copy number represented as fold change. (C) Heat map representing the expression of electron transport chain complex I-V proteins determined by targeted mass spectrometry. (D) *Left:* Transmission electron microscopy images of liver tissue from control (C) and *Mlkl*$^{HepKO}$ (KO) mice fed Control (CD) or Western diet (WD) for 4 months (Magnification: 1000X, Scale bar= 2 μm). *Right:* Graphical representation of the ultrastructure measurement of mitochondrial area (μm$^2$). Each dot represents an individual mitochondrion (n= 3 mice/group ). (E) *Left*: Immunoblots of proteins involved in mitochondrial dynamics in control (siC) or *Mlkl* knockdown (si*Mlkl)* AML12 cells treated with BSA (375 μM) or a mixture of BSA-PA:OA, 250:125 μM for 48 h. Fission proteins include p-DRP1, DRP1, and FIS1; the fusion protein is OPA1. *Right:* Graphical representation of quantified blots showing the ratio of p-DRP1(ser616) to DRP1; OPA1 and FIS1 expression normalized to Ponceau (loading control). All data are expressed as fold change (n=4/group). (F) Transcript levels of *Mfn2* normalized to HPRT in the livers of control (C) or *Mlkl*$^{HepKO}$ (KO) mice fed either control (CD) or Western diet (WD) for 15 months (*left*), (n=7 mice/group), and siC or si*Mlkl* AML12 cells treated with BSA (375 μM) or a mixture of BSA-PA:OA, 250:125 μM for 48 h *(right),* (n=4). (G) *Left:* Immunoblots showing changes in mitochondrial dynamics proteins FIS1, p-DRP1 (Ser616), DRP1, OPA1, MFN1, MFN2, and β−tubulin (loading control) in the livers of control (C) or *Mlkl*$^{HepKO}$ (KO) mice fed either control diet (CD) or Western diet (WD) for 4 months *(top)* or 8 months *(bottom)*. *Right*: Graphical representation of quantified blots showing the ratio of p-DRP1(ser616) to DRP1; OPA1 and FIS1 expression normalized to Ponceau. All data are expressed as fold change (n=4 mice/group). (H) Heat map representing hepatic differential gene changes in control (C) or *Mlkl*$^{HepKO}$ (KO) mice fed either control diet (CD) or Western diet (WD) for 8 months (n=5 mice/group). Red and blue colors represent genes that are significantly upregulated or downregulated. (I) Schematic representation of significantly downregulated or upregulated pathways obtained from RNA seq analysis of liver tissues from *Mlkl*$^{HepKO}$ WD (KO-WD) vs. Control WD (C-WD) fed mice for 8 months. Red and blue colors represent pathways that are significantly upregulated or downregulated. (J) Relative RNA expression of DNA replication factors: *Mcm3*, *Mcm6*, and *PolE* normalized to β-microglobulin and expressed as fold change (n=5 mice/group). (K) *Left:* Representative immunofluorescence images (IF) for iNOS staining (green) and Hoechst (blue) in tumor and non-tumor regions from control (C) or *Mlkl*$^{HepKO}$ (KO) mice fed WD for 15 months (n=3 mice/group). White arrows indicate iNOS positive cells. *Right:* Quantified iNOS positive cells per 63X field (Each dot represents an individual image taken; Scale Bar=25 μm). (L) *Top:* Representative confocal microscopy images for CD206 (green) and Hoechst (blue) staining within non-tumor and tumor regions of control (C) or *Mlkl*$^{HepKO}$ (KO) mice fed control diet (CD) or Western diet (WD) for 15 months. *Bottom:* Bar chart displaying the CD206 positive cells per 63X field; Scale Bar: 25 μm. (M) Transcript levels of inflammatory cytokines: *Tnfa, Il1β,* and *Il6,* and chemokines :*Ccl2, Cxcl9, Cxcl10* in the livers of control (C) and *Mlkl*$^{HepKO}$ (KO) mice 15 months post diet feeding, (n=7 mice/group). (M) The mRNA levels of inflammatory cytokine markers (*Tnfa, Il1β, Il6*) and chemokines (*Ccl2, Cxcl1,Cxcl10*) in the livers of control (C) or *Mlkl*$^{HepKO}$ (KO) fed either control diet (CD) or Western diet (WD) for 8 months, (n=5 mice/group). (N) Heat map showing the expression of pro-inflammatory protein panel in the livers of control (C) or *Mlkl*$^{HepKO}$ (KO) mice fed either Control diet (CD) or Western diet (WD) for 8 months. Results were obtained from proteome profiler analysis (n=4 mice/group). * or # represents statistical significance between control CD vs control WD or control CD vs *Mlkl*$^{HepKO}$ WD, respectively. Data represented as mean ± SEM. Data points marked with asterisks (*) indicate statistically significant differences comparing the mean of each group with that of every other group. One-way ANOVA $p < 0.05$. *$p<0.05$, **$p< 0.01$, ***$p<0.001$, ****$p<0.0001$.

**Figure S7.** (A-C) Oxygen consumption rate (OCR) measured by Seahorse XF Cell Mito Stress Test, with sequential addition of oligomycin, FCCP, and rotenone/antimycin A (R+A). *Left:* Representative oxygen consumption rate (OCR) curve for: (A) HepG2 siC or si*MLKL* cells, (B) Primary hepatocytes isolated from male or female wild type (WT) or *Mlkl* knockout (*Mlkl*$^{-/-}$) mice, and (C) AML12 siC or si*Mlkl* cells. *Right:* Bar graph summarizing the quantified respiratory parameters. Each dot represents an individual experiment. (D) Immunoblots showing the expression of MFN2 and MLKL after subjecting liver lysates of control (C) or *Mlkl*$^{HepKO}$ (KO) mice fed either Control diet (CD) or Western diet (WD) for 15 months to immunoprecipitation (IP) using anti-MLKL antibody alongside IgG negative control. (E) Reciprocal co-immunoprecipitation performed on liver lysates of control (C) or *Mlkl*$^{HepKO}$ (KO) mice fed either Control diet (CD) or Western diet (WD) for 15months using an anti-MFN2 antibody alongside IgG negative control. Western blot shows MLKL and MFN2 expression. (F) Representative immunofluorescence microscopy images demonstrating spatial distribution and proximity of MLKL and MFN2 in AML12 cells. Cells were treated with BSA control (375 μM) or a combination of BSA-PA: OA (250:125 μM) for 48 h. White arrows indicate areas of notable interaction (Scale bar=25 μm). Negative control: MLKL/IgG interaction, and positive control: MLKL/MLKL interaction. (G) *Top:* Immunoblots showing protein expression levels of MLKL,

MFN2 and β-actin (loading control) in control siRNA (siC), *MLKL* siRNA (si*MLKL*-50 nM), *MFN2* siRNA (50 nM) or *MLKL* + *MFN2* (50 nM+50 nM) siRNA transfected HepG2 cells. *Bottom:* Quantified immunoblots showing the fold change of MLKL and MFN2 normalized to β-actin. (H) *Left:* Western blots showing MFN2 and MLKL expression in siC or si*MLKL* HepG2 cells treated with BSA (375 μM) or BSA-PA:OA (250:125 μM), with (+) or without (-) 10 μM MG132 for 4 h. Ponceau staining served as the loading control. *Right:* Quantified MFN2 and MLKL immunoblots as fold change normalized to Ponceau. Each dot represents an individual experiment (n= 3/group). (I) High resolution fluorescence microscopy images showing mitochondrial networks (Mitotracker red stain) in HepG2 cells transfected with siC, si*MLKL*, si*MLKL*+ si*MFN2*, or si*MFN2*. (J) Proteomics analysis of glycolysis proteins in the livers of control (C) or $Mlkl^{HepKO}$ (KO) mice fed Control (CD) or Western diet (WD) for 8 months *(top)* or 15 months *(bottom)*, (n= 5 mice/group). Data represented as mean ± SEM. Data points marked with asterisks (*) indicate statistically significant differences comparing the mean of each group with that of every other group. One-way or Two-way ANOVA (J) ; $p < 0.05$. *$p<0.05$, **$p< 0.01$, ***$p<0.001$, ****$p<0.0001$.

**Figure S8.** (A) Representative immunoblot analysis of MLKL in isolated subcellular fractions from AML12 and HepG2 cell lines. Fractions include cytoplasmic (C), nuclear (N), and mitochondrial (M) compartments. Compartment-specific loading markers were used to verify fraction purity: GAPDH (cytoplasm), Lamin B1 (nucleus), Calnexin (endoplasmic reticulum), and VDAC (mitochondria). Ponceau staining served as the loading control. (B) Total body weight (in grams) of nude mice used in the *in vivo* tumor xenograft model. Measurements compare the vehicle control group (DMSO, n=6, white bar) against the human MLKL inhibitor, Necrosulfonamide treated group (NSA, n=6, red bar). Data represented as mean ± SEM. Data points marked with asterisks (*) indicate statistically significant differences. Two-tailed unpaired t-test; $P<0.05$.
