## Supplementary methodology for "Hepatocyte MLKL Drives Obesity-Driven Hepatocellular Carcinoma Progression via Mitochondrial Dysfunction Independent of Necroptosis in MASLD"

### SUPPLEMENTARY METHODS

#### S1. Cell Culture

AML12 mouse hepatocyte cell line (CRL2254) was obtained from the American Type Culture Collection (ATCC) and maintained with AML12 complete media: Dulbecco's modified eagle medium (DMEM; 30-2002, ATCC) supplemented with 1x insulin-transferrin selenium supplement (ITS, 41-400-045, Thermo Scientific), 40 ng/mL dexamethasone (D4902-25MG, Millipore Sigma), 1% Penicillin- Streptomycin (15070063, Fisher Scientific) and 10% fetal bovine serum (FBS, A5669801, Fisher Scientific). Cells were incubated at 37°C with 5% $CO_2$. HepG2 cells (HB-8065, ATCC) and Hep3B (HB-8064) from ATCC were maintained in Eagle's Minimum Essential Medium (EMEM, 30-2003, ATCC) and 10% FBS at 37°C with 5% $CO_2$. Liver cancer panel from ATCC (TCP-1011) that included different human liver cancer cell lines (C3A, SK-Hep-1, SNU-475, SNU-449, SNU-387, SNU-423) was purchased and cell lines maintained in the recommended culture media. Hepa1-6 cells (CRL1830, ATCC) and Huh-7 cells (JCRB0403, Sekisui Diagnostics) were maintained in DMEM supplemented with 10% fetal bovine serum and 1% Penicillin-Streptomycin at 37°C with 5% $CO_2$. BV-2 cells (ABC-TC212S, AcceGen Biotechnology, Fairfield, NJ) were maintained in DMEM and 10% FBS at 37°C with 5% $CO_2$. All cell culture studies were performed in triplicates.

#### S2. Western Blot Analysis

Western blot analysis was performed and quantification done using Image J software as described previously(1) . After western blotting, Blazin' Blue™ Protein Gel Stain (P-810-1, Gold Biotechnology) was used to stain the gels, and Ponceau S solution (P7170-1L, Sigma-Aldrich) was used to stain the blots as per manufacturer's instruction. Images were taken using the ChemiDoc imager (Bio-Rad) and quantified with ImageJ software (U.S. National Institutes of Health). Primary antibodies used in this study are listed in **Table S1**. Quantified blots are graphically represented alongside representative immunoblots.

#### S3. Quantitative RT-PCR

Isolation of total RNA and cDNA synthesis was performed as described previously(2). The primers used for RT-PCR are listed in **Table S2.** Results were computed by the comparative method ($2^{-\Delta\Delta}$Ct) using β-microglobulin, β-actin, or hypoxanthine phosphoribosyltransferase 1 (HPRT) as controls, as described previously. The expression of genes and heat map representation was performed using GraphPad Prism.

#### S4. Plasma ALT and Hepatic Triglyceride Assays

Plasma ALT and hepatic triglyceride content were quantified using ALT or triglyceride colorimetric assay kit (700260 and 10010303, Cayman Chemical Company, USA). Procedures for measurements were conducted following the manufacturer's instruction.

#### S5. Histology and Histomorphometry

Formalin-fixed liver was paraffin-embedded, and sections were stained with Hematoxylin and Eosin (H&E) according to standard protocol at the Stephenson Cancer Center Tissue Pathology core, University of Oklahoma Health Campus. Images were acquired using the ECHO Revolve Microscope (Discover Echo Inc, San Diego, CA) at 10X magnification for 3 random non-overlapping fields per sample. Lipid droplet number and area were measured using Adiposoft (v1.16) Fiji (v2.15.0) previously described (3)

#### S6. Picrosirius Red Staining

Sectioned paraffin-embedded liver sections (4 μM) were stained with Picrosirius Red following a standardized protocol at the Stephenson Cancer Center Tissue Pathology Core, University of Oklahoma Health Campus. Images were acquired using the ECHO Revolve Microscope (Discover Echo Inc) at 10X magnification for 3 random non-overlapping fields per sample. Quantification was done using the Image J software (U.S. National Institutes of Health).

#### S7. Immunohistochemistry for MLKL, GPC3, Cleaved Caspase-3, and Ki-67

IHC was carried out as described previously (4). Paraffin-embedded liver sections were incubated with primary antibodies against MLKL (21066-1-AP, Proteintech), Glypican 3 (PA5-102528, Thermo Scientific), cleaved-caspase-3 (9664, Cell Signaling), or Ki-67 (ab15580, Abcam) overnight at 4°C. Diaminobenzidine-based colorimetric method was used for the detection of target proteins in the sections. Nuclei were counter stained with Mayer's Hematoxylin (MHS16, Sigma-Aldrich). The human liver cancer tissue microarray is from US Biolab DLV1002. Images were taken with ECHO Revolve Microscope (Discover Echo Inc).

#### S8. Hydroxyproline Assay for Collagen Content

The liver collagen content was measured using hydroxyproline (OHP) assay as described previously(4).Absorbance measurements were determined at 558 nm and converted into μg units using the 4-parameter standard curve generated from standard concentrations of trans-4-hydroxy-L-proline (H5534, Sigma-Aldrich). Calculated OHP content was expressed as μg hydroxyproline/g of tissue.

**S9. siRNA mediated knockdown of *Mlkl, MLKL*, *MFN2*, or *MLKL*/*MFN2* double knockdown in AML12 or HepG2 cells**

AML12 or HepG2 cells were transfected with 100 nM of siControl (Silencer Select Negative Control No.1 siRNA, Cat. No. 4390843, Invitrogen, Fisher Scientific) or si*Mlkl/MLKL* (Silencer Select siRNA against mouse or human MLKL, Cat. No. 4390771 or 4392420, Invitrogen, Fisher Scientific) for 48 h. HepG2 cells were also transfected with 50 nM of si*MFN2* (Silencer Select siRNA against human MFN2, Cat. No. 4392420, Invitrogen, Fisher Scientific), or a combination of si*MLKL* and si*MFN2* (combined concentration of 100 nM) using Lipofectamine RNAiMAX (13778075, Invitrogen, Fisher Scientific) for 48 h (Cell mitostress test) or 6 days (Cell proliferation by Incucyte live cell analysis). The efficiency of knockdown was determined using Western blot analysis.

To determine changes in mitochondrial fission/fusion proteins, AML12 cells were seeded in 24-well plate (8 x $10^4$ cells per well) overnight. The cells were transfected with either 100 nM siControl or si*Mlkl* as described above for 48 h. The transfected cells were subsequently exposed to either BSA Control for Fatty Acid Complexes or a combination of 250 μM BSA-Palmitate and 125 μM BSA-Oleate (Cat. No. 29556, 29558, 29557; Cayman Chemical Company) in AML12 complete media with 2.5% Fetal Bovine Serum (FBS) for 48 h. Western blot analysis was performed to determine changes in protein expression of MFN1, MFN2, OPA1, p-DRP1 (S616), DRP1 and FIS1.

**S10. Oil Red O Staining**

AML12 cells, HepG2 cells, and primary hepatocytes isolated from male or female wild-type and *Mlkl-/-* mice were seeded in 24-well plates at a density of 8 × $10^4$ cells/well for AML12 cells and 1 × $10^5$ cells/well for HepG2 cells and primary hepatocytes. AML12 or HepG2 cells were transfected with siRNA (100nM siControl or *siMlkl/siMLKL*) for 48 h, whereas isolated primary hepatocytes were incubated for 36 h. The cells were then exposed to either BSA Control for Fatty Acid Complexes or a combination of 250 µM BSA-Palmitate and 500 µM BSA-Oleate (Cat. No. 29556, 29558, 29557; Cayman Chemical Company) in DMEM F12, EMEM or DMEM complete media with 2.5% Fetal Bovine Serum (FBS) for 48 h. The accumulation of lipids in cells was visualized using Oil red O staining and quantified by 100% isopropanol extraction as described previously (5). Briefly, cells were rinsed with distilled water and fixed with 4% paraformaldehyde for 30 minutes at room temperature. The cells were rinsed with distilled water, followed by rinsing with 60% isopropanol. Oil Red O was added for 15 minutes and then rinsed with distilled water before taking images using the ECHO Revolve Microscope (Discover Echo Inc) at 20X magnification (3 random non overlapping images for each sample). For quantification, Oil Red O dye was washed with water and then 500 µL 100% isopropanol was added to each well and incubated for 10 minutes with gentle shaking. Absorbance readings of extract were recorded in 96-well plate at 510 nM. Fold change of read absorbances were represented graphically.

**S11. Targeted Mitochondrial Proteomics**

Quantitative targeted proteomics analysis was carried out to evaluate changes in hepatic mitochondrial enzymes as previously described (6). The quantified protein abundances were represented graphically. The raw data obtained from this is provided in **Table S4, 6, 8**.

**S12. RNA Sequencing and Bioinformatic Processing**

Total RNA was isolated from the liver tissue using RNeasy kit (74106, Qiagen, Germantown, MD). RNA library construction and sequencing was carried out by the Institutional Research Core Facility at OUHSC. Briefly, RNA was checked for quality using Agilent's 4150 Tapestation, and concentrations from ThermoFisher's NanoDrop readings were used. Stranded RNA-seq libraries were constructed using NEBNext poly (A) mRNA isolation kit followed directly by IDT's XGen Broad Range RNA Library Prep Kit and the established protocols. The library construction was done using 1µg of RNA. Each of the libraries was indexed during library construction to multiplex for sequencing. Libraries were quantified using Invitrogen's Qubit 4 fluorometer and checked for size and quality on Agilent's 4150 Tapestation. Samples were normalized and pooled onto a 150 paired end run on Illumina's NextSeq 2000 Platform to obtain 20M reads per sample. Expression values quantile normalized with the voom function were analyzed for differential expression using the standard functions of the limma package. Moderate t-test p-values were adjusted for multiple testing using the false discovery rate (FDR) method. FDR (q.value) <0.05 and absolute log2 fold change above 1 were used as criteria to filter significantly differentiated genes. Genes significantly changed by WD diet in control mice, reverting to CD diet levels after knocking out hepatocyte *Mlkl* were identified by collapsing Control-CD mice with both $Mlkl^{HepKO}$ WD and $Mlkl^{HepKO}$ CD mice into a single group and contrasting it to Control-WD mice. Since the t-statistics consider the difference between group averages relative to within-group variation, a strong t-statistic would indicate both strong difference between Control-WD mice versus the other groups, and similar expression levels in the Control-CD and both groups of CRE treated mice. Gene Set Enrichment Analysis (GSEA) was conducted using specialized Bioconductor packages, including fgsea, ReactomePA, and viewPathway, to identify functionally related gene sets (e.g., GO terms, KEGG and Reactome pathways) that were significantly overrepresented among the differentially expressed genes. Each pathway GSEA edge set was used as basis of selection for genes included in the heatmaps showing the pathway activity variation among the four mice phenotypes. Ingenuity Pathway Analysis (IPA, QIAGEN, RedwoodCity CA, https://www.qiagenbioinformatics.com/products/ingenuitypathway-analysis) was used for further discovery and interactive exploration of significantly impacted statistics and causal gene networks, pathways, disease, upstream regulators, and regulatory effects.

**S13. Mitochondrial DNA (mtDNA) Copy Number Assay**

Genomic DNA was isolated from the liver tissue (25 mg) using the QIAamp DNA Kits (56304) for DNA extraction following manufacturer's instructions. Mitochondrial content was quantified using the QuantStudio Absolute Q Digital PCR System according to the protocol described (7). Briefly, for each sample, a mix of 2 µL of Absolute Q Universal DNA Digital PCR Master Mix and 0.5 µL of the appropriate primer mix (mtDNA or nucDNA) was made, with overage. DNA was diluted to a concentration that optimized the precision of the copy number then combined with 2.5 µL of the mix. A final reaction volume of 10 µL was achieved by adding an appropriate amount of water to each. To load the dPCR plate, 9 µL of the final reaction was added to a well. Following that, 15 µL of Absolute Q Isolation Buffer was overlaid on top of the 9 µL of the final reaction. The wells were covered with a Quant Studio Absolute Q MAP16 Digital PCR Gasket Strip. The plate was then loaded into the Quant Studio Absolute Q Digital PCR System. The digital PCR was conducted using the following protocol: 10 min. at 96°C, followed by 40 cycles of 5 seconds at

96°C and 15 seconds at 60°C. The ratio of mtDNA to nuclear DNA was then converted to fold change from control.

**S14. Cell Cycle Analysis**

Cell cycle distribution was assessed by propidium iodide staining as previously described (Lankadasari et al., 2018). HepG2 cells ($0.5 \times 10^6$) were treated with necrosulfonamide (NSA; 5 µM) or transfected with siControl or siMLKL (100 nM) for 48 h. Cells were fixed in ice-cold 70% ethanol, treated with RNase A (50 µg/mL) for 10 min, and stained with propidium iodide (50 µg/mL) for 30 min at room temperature in the dark. DNA content was analyzed on a Stratedigm S3 flow cytometer, and cell cycle distribution was determined using ModFit LT software.

**S15. Cell Proliferation and Spheroid Growth Assays**

Proliferation and spheroid growth assays were performed as described previously(8). Briefly, for proliferation assay, HepG2 cells were plated as 8000 cells/well and MLKL was inhibited pharmacologically using 5 µM NSA or DMSO control (0.05%), or genetically by siRNA mediated knockdown of MLKL (100 nM) for 4 days. For cell proliferation experiments involving double knockdowns, single knockdowns were performed using 50 nM of si*MLKL* or si*MFN2*. Double knockdowns were performed using a total siRNA concentration of 100 nM (50 nM of each siRNA) for 5 days. Corresponding siControls with 50 nM and 100 nM were maintained. Human *RIPK3* was overexpressed using 1µg pcDNA-*RIPK3*-Flag (78815, Addgene, Watertown, MA or control plasmid (pcDNA3.1+C-DYK, GenScript, Piscataway, NJ) for 4 days. Live cell imaging was done using Incucyte S3 live-cell analysis system (Sartorius, Ann Arbor, MI). Live cell images were acquired in standard scan type mode with a 10X objective, and 4 images were captured per well. The percentage confluence was analyzed and plotted against time.

For spheroid growth assays, HepG2 cells were plated as 3000 cells/well in a round bottom ultralow attachment plate (7007, Corning), centrifuged at 1500 rpm for 10 minutes. The cells were either pharmacologically or genetically inhibited for MLKL as described above. The formation of spheroids was then monitored using Incucyte live cell imaging system. Images were acquired in spheroid scan type mode for single spheroids with a 4X objective and single image per well. Spheroid formation was assessed by calculating the largest brightfield object area.

**S16. Measurement of Mitochondrial Respiration**

Mitochondrial respiration was measured using the Seahorse XF96 Cell Mito Stress Test (Agilent Technologies, Santa Clara, CA). AML12 cells transfected with siControl or si*Mlkl* for 48 h and treated with BSA-conjugated palmitate (50 μM) or BSA vehicle control (50 μM) for 48 h prior to analysis. For cell mito-stress test with HepG2 cells, cells were transfected with either 50 nM of si*MLKL* or si*MFN2* or double knockdown of si*MLKL* and si*MFN2* (100 nM total concentration) along with corresponding siControls (50 nM or 100 nM) for 48 h. For experiments with primary hepatocytes, cells were analyzed after 36 h of plating. Oxygen consumption rate (OCR) was determined following sequential injection of oligomycin (1.5 μM), FCCP (2 μM), and rotenone/antimycin A (0.5 μM each), and respiratory parameters were calculated using Seahorse Wave software (Agilent Technologies)(9).

**S17. Glycolysis stress test**

HepG2 cells were transfected with siControl (50 or 100 nM), si*MFN2* (50 nM), si*MLKL* (50 nM), or a combination of si*MFN2* (50 nM) and si*MLKL* (50 nM). Twenty-four hours after transfection, glycolytic function was assessed using a Seahorse XF Glycolysis Stress Test (Agilent Technologies). Cells were incubated in Seahorse XF assay medium (103575-100, Agilent) containing 2 mM L-glutamine (103579-100, Agilent) and lacking glucose and pyruvate (180 μL/well) for 1 h at 37C in a non-carbon dioxide incubator. The extracellular acidification rate (ECAR) was measured following sequential injections of glucose (10 mM, Port A), oligomycin (1 μM, Port B), and 2-deoxy-D-glucose (2-DG; 50 mM, Port C). Glycolytic parameters, including glycolysis, glycolytic capacity, and glycolytic reserve, were calculated using Seahorse Wave software according to the manufacturer's instructions.

**S18. Transmission Electron Microscopy**

The TEM experiment was performed at the Imaging core facility, Oklahoma Medical Research Foundation (OMRF) imaging facility using procedures as described before(6). TEM micrographs were analyzed using ImageJ (National Institutes of Health, Bethesda, MD, USA). Images were calibrated using the microscope scale bar prior to analysis Individual mitochondria were manually outlined in ImageJ, and mitochondrial area was quantified and expressed as $\mu m^2$. Lipid droplets (LDs) and mitochondria were identified based on established ultrastructural characteristics. The distance between each LD and the nearest mitochondrion was measured manually using the ImageJ straight-line tool, recording the shortest edge-to-edge distance between the LD surface and the outer mitochondrial membrane. Distances were expressed in nanometers (nm). Measurements were performed on multiple representative fields from each sample, and data were averaged for each animal. Analyses were conducted using three biological replicates per group (n = 3/group), and results are presented as mean ± SEM.

#### S19. Immunofluorescence Staining for MLKL, TOM20, iNOS, and CD206

Immunofluorescence staining was done as described (10). Briefly, paraffin sections (4 µm) were deparaffinized and rehydrated. Heat-induced antigen retrieval was performed, followed by permeabilization with 0.1% Triton X-100. Sections were blocked and then incubated with primary antibody MLKL (21066-1-AP, Proteintech, Rosemont, IL), TOM20 (66777-1-Ig, Proteintech), iNOS (18985-1-AP, Proteintech), or CD206 (AF2535, R&D Systems, Minneapolis, MN) at 4°C overnight. Coresponding Alexa Fluor-conjugated secondary antibody was applied. The nuclei were stained with DAPI (62248, Thermo Scientific) or Hoechst 33342 (5117, R&D Systems). Images were taken with Leica SP8 Confocal microscope.

#### S20. Proximity Ligation Assay (PLA)

PLA was carried out with Duolink PLA kit (DUO92008, Sigma-Aldrich) and Probes (DUO92002 and DUO92004, Sigma-Aldrich) as described (10). Briefly, cells were cultured on coverslips. After fixation, permeabilization, and blocking, primary antibody mouse anti-MLKL (66675-1-Ig, Proteintech), and rabbit anti-MFN2 (12186-1-AP, Proteintech), or Normal Rabbit IgG (2729, Cell Signaling Technology, Danvers, MA), or rabbit anti-MLKL (MA553520, Invitrogen, Carlsbad, CA) were used for incubation at 4°C overnight. Duolink PLA probe incubation, ligation, and amplification were performed according to the manufacturer's instructions at 37°C for 1 h, 30 min, and 100 min, respectively. Nuclei were counterstained with DAPI. Fluorescence images were acquired using a Leica SP8 confocal microscope (Leica Biosystems, Deer Park, IL).

#### S21. Immunoprecipitation

For immunoprecipitation, liver lysates were pre-cleared with Protein A/G magnetic beads (88802, Thermo Scientific) and then incubated with 1 µg of control IgG, anti-MLKL antibody (MABC604, Millipore), or anti-MFN2 antibody (12186-1-AP, Proteintech) per 1 mg of protein at 4°C overnight. The lysates were subsequently incubated with Protein A/G magnetic beads at 4°C for 1 h. After washing with NP-40 wash buffer [50 mM Tris-HCl (pH 7.5), 150 mM NaCl, 2 mM EDTA, and 1% NP-40], the immunoprecipitates were resuspended in 2× Laemmli sample buffer, heated at 95°C for 5 min, chilled on ice, and subjected to immunoblotting.

**S22. Mitotracker Staining**

Cells were seeded on Nunc™ Lab-Tek™ II Chambered Coverglass (12-565-338, Thermo Scientific). At 50-60% confluency, cells were stained with pre-warmed (37°C) 500 nM MitoTracker® probe (M22425, Invitrogen) for 30 min. Images were taken with Leica SP8 Confocal microscope.

**S23. Untargeted Lipidomics Analysis**

Untargeted lipidomics analysis was performed by Creative Proteomics (Shirley, NY, USA) using UPLC–MS to profile the liver lipid composition. The resulting raw data are provided in Table S5.

**S25. Proteasome inhibition assay**

AML12 or HepG2 cells (5 x $10^5$) were seeded in 6-well plates. The cells were transfected with siRNA (100 nM siControl or *siMlkl/siMLKL*) for 48 h. These cells were subsequently treated with either 375 µM BSA Control or a combination of 250 µM BSA-Palmitate and 125 µM BSA-Oleate (Cat. No. 29556, 29558, 29557; Cayman Chemical Company) in DMEM F12 or EMEM complete media with 2.5% Fetal Bovine Serum (FBS) and incubated for 48 h. 10 µM MG132 in DMSO (Cat. No. 474790, EMD Millipore) or DMSO (0.05%) (Cat. No. D2650, Sigma Aldrich) was added to the cells 4 h towards the end of 48 hours incubation with fatty acids. The cells were lysed and supernatant collected for Western blot analysis as described in Mohammed et al, 2021. Changes in protein expression of MFN2 (12186-1-AP, Proteintech) were quantified by western blotting.

**S26. NADH Oxidase/Complex I Activity Assay**

Mitochondrial functions were assessed by electron transport chain (ETC) enzyme activities. As previously described (11) mitoplasts were isolated from frozen livers. Briefly, 25 mg of thawed tissues were homogenized in 400 µL of isolation buffer containing 1.0 mM EDTA and 25 mM MOPS pH 7.4 by a motor-driven Potter-Elvehjem tissue grinder. The homogenate was spun at 750 g for 10 min at 4 °C, and the supernatant was then spun again at 10000 g for 15 min. The resulting mitoplasts pellet was resuspended in 25 mM MOPS pH 7.4 and immediately snap frozen. NADH oxidase, an overall assessment of electron transport chain activity through complexes I–III–IV was measured as follows. Twice frozen-thawed mitoplasts were diluted into 25 mM MOPS pH 7.4 at roughly 250 µg/mL and the rotenone-sensitive rate of NADH oxidation was monitored spectrophotometrically utilizing an Agilent 8453 diode array UV−Vis spectrophotometer. Activity was measured as the rate of NADH oxidation (340 nm, ε = 6200 M–1 cm–1) following addition of 10 mM KCl and 150 µM NADH. Similarly, complex I activity was measured spectrophotometrically as the rate of NADH oxidation in the presence of 50 nM antimycin A and 100 µM ubiquinone-1 following the addition of 150 µM NADH. The specificity for complex I activity was confirmed by inhibition with 1.25 µM rotenone. The protein concentration of each sample was determined by the Bradford method with a BSA as the standard.

**S27. F2-Isoprostane Analysis**

F2-isoprostanes were quantified by gas chromatography/mass spectrometry (GC/MS). Briefly, 30–50 mg of snap-frozen liver tissue was collected for each sample, and tissue weights were recorded. Each sample was placed in a 2 ml round-bottom tube containing 1 ml of cold methanol supplemented with 50µg/ml butylated hydroxytoluene (BHT). Sterile beads were added to each tube, and samples were homogenized in a shaking tissue lyser at 25 rpm for 30 sec. To release esterified isoprostanes, 500µl of 4 M potassium hydroxide was added to each homogenate and incubated at 37°C for 30 min. Homogenates were then transferred to 15 ml polypropylene conical tubes and acidified to pH 2-3 using 400-500 µl of 4 M hydrochloric acid. An internal standard consisting of 2 ng tetradeuterated 15-F2t-isoprostane in 10 µl ethanol was added to each sample, followed by equilibration at 4°C for 15 min. For ethyl acetate extraction, 10 ml of ethyl acetate was added to each sample to achieve a 5:1 ratio of ethyl acetate to the methanol/aqueous solution. Samples were vortexed for 1 min, and the phases were separated by centrifugation at 4,000 × g for 5 min.

For aminopropyl solid-phase extraction (SPE), an aminopropyl SPE column fitted with a 12-cc cartridge was activated with 5 ml of heptane using a vacuum manifold. The upper organic layer from each sample was transferred onto the column by pipetting, while the lower aqueous phase was retained for subsequent protein estimation. After sample loading, each column was washed with 5 ml of ethyl acetate and eluted into a scintillation vial with 5 ml of a methanol/ethyl acetate/acetic acid (85:10:5, v/v/v) solution. Eluted samples were completely dried under nitrogen in an evaporator at 37°C. The residue was reconstituted in 200 µl methanol and stored overnight at −20°C.

Negative-ion chemical ionization gas chromatography/mass spectrometry (NICI-GC/MS) was performed using an initial oven temperature of 190°C and a final temperature of 300°C with a ramp rate of 15°C/min. Methane was used as the reagent gas. The *m/z* 569.4 anion was monitored for TMS-derivatized F2-prostaglandin isomers, whereas the *m/z* 573.4 anion was monitored for the tetradeuterated internal standard. The chromatographic peak height of the lighter ion that eluted at the same retention time as the 15-F2t-isoprostane internal standard was measured. F2-isoprostane abundance was calculated as the ratio of analyte peak height to internal standard peak height, multiplied by the 2 ng internal standard, and normalized to tissue weight according to the following equation:

$$\frac{analyte\ peak\ height}{internal\ standard\ peak\ height} \times 2\ ng \Big/ tissue\ mass\ (g)$$

The retained aqueous phase was used for protein estimation by precipitating proteins with cold acetone, centrifuging to pellet the proteins, and removing the supernatant until no aqueous phase remained. Residual acetone was completely evaporated under nitrogen, and the protein pellet was saponified with 1 ml of 1 M sodium hydroxide. Protein concentration was determined using the bicinchoninic acid (BCA) assay and expressed as mg protein/ml NaOH solution. Isoprostane concentration normalized to protein content was expressed as pg isoprostane/mg protein.

**S28. Detection of 4-Hydroxynonenal (4-HNE) Adducts**

4-HNE modified proteins were detected by western blotting as described previously (1, 12).

**S29. Soluble and Insoluble Protein Extraction**

The soluble and insoluble fractions form liver lysates were isolated as described previously with modifications(13). Briefly, approximately 50 mg of snap-frozen liver tissue was homogenized in ice-cold radioimmunoprecipitation assay buffer (RIPA buffer; PI89901, Fisher Scientific) supplemented with Protease inhibitor (GB-331-5, Goldbio, St. Louis, MO), Phosphatase inhibitor (GB-451-5, Goldbio) and Phenylmethanesulphonyl fluoride (77510-384, VWR Chemicals, Radnor, PA) cocktail. Homogenates were centrifuged at 14000 rpm for 20 min at 4°C, and the supernatant was collected as the detergent-soluble protein fraction. The supernatant was centrifuged a second time under the same conditions to remove any remaining insoluble material. The resulting pellet was resuspended in RIPA buffer containing 8 M urea and disrupted using stainless steel bead–based mechanical homogenization to solubilize detergent-insoluble proteins. Protein concentrations were determined using the Bradford protein assay (500006, Bio-Rad Protein Assay Dye Reagent Concentrate, Bio-Rad Laboratories, Hercules, CA) according to the manufacturer's instructions. Equal amounts of protein (40 μg) from the soluble and insoluble fractions were resolved by SDS-PAGE, transferred to PVDF membranes, and subjected to immunoblotting using antibodies against RIPK1, RIPK3, p-RIPK3, MLKL, and p-MLKL. Liver lysates from *Ripk3*$^{-/-}$ mice and *RIPK3*-overexpressing HepG2 cells were used as negative and positive controls, respectively, for RIPK3 detection, while lysates from BV2 cells treated with zVAD-fmk served as a positive control for p-MLKL.

**S30. Subcellular Fractionation and Immunoblotting**

Cytoplasmic and nuclear protein extraction was performed using the Boster Cytoplasmic and Nuclear Protein Extraction Kit (AR0106, Boster Bio, Pleasanton, CA) according to the manufacturer's protocol. Briefly, AML12 and HepG2 cells ($2 \times 10^6$ cells per condition) were harvested and washed with 1X PBS. The cell pellet was lysed with Cytoplasmic Extraction Reagent A/B and centrifuged to isolate the cytoplasm. The supernatant containing the cytoplasmic fraction was collected. The remaining pellet was lysed with Nuclear extraction reagent and centrifuged to yield the nuclear protein fraction. Both fractions were stored at -80 °C until usage.

Intact mitochondria were isolated from AML12 and HepG2 cells ($2 \times 10^6$ cells per condition) using the Mitochondria Isolation Kit for Cultured Cells (HY-K1060-100T, MedChemExpress, Monmouth Junction, NJ) according to the manufacturer's protocol. Briefly, cells were harvested, washed with ice-cold PBS, and lysed in the kit's isolation buffer supplemented with PMSF. Differential centrifugation was performed to remove nuclei and unbroken cells, followed by pelleting of the crude mitochondrial fraction, with the resulting supernatant retained as the cytosolic fraction. Mitochondrial fraction was resuspended in HEPES lysis buffer.

Protein concentrations were determined by Bradford assay. Equal amounts of protein from mitochondrial, nuclear and cytosolic fractions were resolved by SDS-PAGE and immunoblotted for MLKL. Fractionation purity was confirmed using VDAC1 (mitochondrial marker) and GAPDH (cytosolic marker), Calnexin (ER marker) and Lamin B1 (nuclear marker).

**S31. Enzyme-Linked Immunosorbent Assay (ELISA)**

Mouse alpha-fetoprotein (AFP) plasma level was detected with Mouse alpha-fetoprotein (AFP) quantikine ELISA kit (MAFP00; R&D Systems).

**S32. LDH Release Assay**

Cell death was assayed by measuring the release of lactate dehydrogenase (LDH) into cell culture medium (14) using CyQUANT™ LDH Cytotoxicity Assay kit (C20300, Thermo Scientific) by following manufacturer's instructions.

**S33. *In vivo* Xenograft Studies with NSA Treatment**

All animal procedures were approved by the Institutional Animal Care and Use Committee of the University of Oklahoma Health Sciences Center (IACUC, OUHSC) and performed in accordance with institutional and NIH guidelines for the care and use of laboratory animals.

Male athymic nude mice (Strain #:002019, 4 weeks old, Jackson Laboratory, Bar Harbor, ME) were housed under specific-pathogen-free conditions with a 12-h light/dark cycle and ad libitum access to food and water. HepG2 cells ($7 \times 10^6$ cells in 50 µl of [sterile PBS:Matrigel(356234, Corning) 1: 1 ratio]) were subcutaneously injected into the right flank of each mouse. Tumors were allowed to establish for 3 weeks, at which point mice with palpable, visible tumors were randomized into treatment groups (n = 6/group) to ensure comparable baseline tumor volumes between groups. Mice received intraperitoneal injections every other day for 4 weeks of either vehicle control (0.1% DMSO in PBS) or necrosulfonamide (NSA; 10 mg/kg body weight, dissolved in 0.1% DMSO/PBS) (480073, Sigma Aldrich). Tumor volume was measured every 3 days using digital calipers and calculated using the modified ellipsoid formula: $V = (L \times W^2) / 2$ where L is the longest diameter and W is the perpendicular width. Mice were monitored daily for signs of distress and were euthanized if tumor volume reached 1.5 $cm^3$, as per IACUC-OUHSC humane endpoint criteria, or at the conclusion of the 4-week treatment period, whichever occurred first. At the study endpoint, mice were euthanized by $CO_2$ asphyxiation followed by cervical dislocation, and tumors were excised, weighed, and either snap-frozen in liquid nitrogen or fixed in 10% neutral-buffered formalin for downstream molecular and histological analyses. Blood was collected in EDTA-coated tubes and centrifuged at 2000 x g for 10 minutes at 4°C, the plasma was collected. Mouse plasma "Preventive Care Profile Plus" profile analysis was done by the Division of Comparative Medicine at University of Oklahoma Health Campus.

**S34. Colony Formation Assay**

HepG2 cells (5000 cells/well) transfected with 100 nM siControl or siMLKL, or treated with 5 μM NSA or DMSO (0.01%) were plated in 6-well plate and left to form colonies for 10 days. The colonies were then fixed with 4% PFA and stained with 0.2% crystal violet (V5265, Sigma-Aldrich). The number of colonies were counted and represented graphically. Further, the crystal violet was dissolved using 10% acetic acid and the absorbance was measured at 595 nm (15).

**S35. Human Dataset Analysis**

The TCGA-LIHC (The Cancer Genome Atlas -Liver Hepatocellular Carcinoma) data set was used to compare the expression level of MLKL between normal liver tissue (n=50) and primary hepatocellular carcinoma (n=371) samples, the result is expressed as MLKL transcripts per million (TPM). Based on this, the patients were stratified into low and high expression groups and correlation between expression level of MLKL and patient survival was examined. The prognosis of the high and low expressing groups was examined by Kaplan-Meier analysis using Kaplan-Meier Plotter, and the survival outcome was compared with log-rank test. A p-value of $<0.05$ is considered statistically significant.

**S36. Necroptosis Induction in BV2 and AML 12 Cells**

BV2 murine microglial cells were treated with the pan-caspase inhibitor z-VAD-fmk (20 μM) for 24 h in the presence or absence of the RIPK1 inhibitor Necrostatin-1 (20 μM). For AML12 cells, $0.5 \times 10^6$ cells were seeded per well and transfected with either pcDNA or RIPK3 expression plasmids as described in Supplementary Methods S15. Following transfection, cells were pretreated with z-VAD-fmk (20 μM) and subsequently stimulated with TNF-α (20 ng/mL) and BV6 (4 μM) for 4 h to induce necroptosis. After treatment, cells were washed with ice-cold PBS, harvested, and lysed in HEPES lysis buffer supplemented with protease and phosphatase inhibitor cocktails. Protein lysates were then subjected to immunoblot analysis of necroptosis pathway markers(16).

**S37: Liver Non-Parenchymal Cell (NPC) Isolation**

The NPC fraction was isolated from the supernatant remaining after hepatocyte isolation that is described in the main methodology section. Briefly, the supernatant was centrifuged at 500 × g for

10 minutes to pellet the NPCs. The resulting pellet was lysed in TRIzol Reagent (Cat. No. 15596018, Invitrogen) and processed for subsequent RNA isolation.

**AI use statement:** Microsoft Copilot was used solely for language editing. All scientific content was reviewed and verified by the authors, who take full responsibility for the accuracy and integrity of the manuscript.
