## Supplementary Tables for "Hepatocyte MLKL Drives Obesity-Driven Hepatocellular Carcinoma Progression via Mitochondrial Dysfunction Independent of Necroptosis in MASLD"

**Table S1** Antibody list

| Antibody | Clone | Catalog Number | Vendor |
|---|---|---|---|
| MLKL | 3H1 | MABC604 | Millipore |
| p-MLKL(S345) | EPR9515(2) | ab196436 | Abcam |
| RIPK3 | | NBP1-77299 | Novus |
| β-tubulin | TUB 2.1 | T5201 | Sigma |
| GPX4 | E-12 | sc-166570 | Santa Cruz |
| Gasdermin D | L60 | 93709 | Cell Signaling |
| Cleaved Gasdermin D | | 50928S | Cell Signaling |
| MFN1 | | 13798-1-AP | Proteintech |
| MFN2 | | 12186-1-AP | Proteintech |
| OPA1 | | ab42364 | Abcam |
| DRP1 | | 12957-1-AP | Proteintech |
| pDRP1 (S616) | | PA5-64821 | Thermo Scientific |
| FIS1 | | 10956-1-AP | Proteintech |
| PERILIPIN-2 | | 15294-1-AP | Proteintech |
| MLKL | 23GB2895 | MA553520 | Invitrogen |
| β-actin | AC-15 | A5441 | Sigma |
| GAPDH | GAPDH-71.1 | G8795 | Sigma |
| ACSL4 | | 22401-1-AP | Proteintech |
| LAMIN B1 | | ab16048 | Abcam |
| CALNEXIN | | 10427-2-AP | Proteintech |
| VDAC | D73D12 | 4661S | Cell Signaling |
| DCLK1 | | 21699-1-AP | Proteintech |
| OCT4 | | 11263-1-AP | Proteintech |
| p-RB (Ser807/811) | D20B12 | 8516 | Cell Signaling |
| p-21 | EPR18021 | ab188224 | Abcam |
| p-CDC2 (Tyr15) | 10A11 | 4539 | Cell Signaling |
| p-CHK1 (Ser345) | 133D3 | 2348 | Cell Signaling |
| p-CHK2 (Thr68) | C13C1 | 2197 | Cell Signaling |
| Glypican 3 | | PA5-102528 | Thermo Scientific |
| cleaved-caspase-3 (Asp175) | 5A1E | 9664 | Cell Signaling |
| Ki67 | | ab15580 | Abcam |
| MLKL | | 21066-1-AP | Proteintech |
| Tom20 | 1D6F5 | 66777-1-Ig | Proteintech |
| iNOS | | 18985-1-AP | Proteintech |
| CD206 | | AF2535 | R&D |
| Normal Rabbit IgG | | 2729 | Cell Signaling |
| p-MLKL(S358) | EPR9514 | ab187091 | Abcam |
| RIPK3 | | 63-216 | ProSci |
| RIPK3 | | 17563-1-AP | Proteintech |
| RIPK1 | | NBP1-77077 | Novus |
| p-RIPK(S166) | | 28252-1-AP | Proteintech |
| 4HNE: gift from Dr. Luke Szweda, Oklahoma Medical Research Foundation (PMID: 8554315) | | | |

**Table S2.** List of primers used for quantitative RT PCR

| Primer name | Forward primer sequence | Reverse primer sequence |
|---|---|---|
| Mlkl | 5'-CTGAGGGAACTGCTGGATAGAG-3' | 5'-CGAGGAAACTGGAGCTGCTGAT-3' |
| β-microglobulin | 5′-CACTGACCGGCCTGTATGC-3′ | 5′-GGGTGGCGTGAGTATACTTGAAT-3′ |
| Albumin | 5’-GCGCAGATGACAGGGCGGAA-3’ | 5’-GTGCCGTAGCATGCGGGAGG-3’ |
| Ccl2 | 5’-TTAAAAACCTGGATCGGAACCAA-3’ | 5’-GCATTAGCTTCAGATTTACGGGT-3’ |
| HPRT | 5’-CTGGTGAAAAGGACCTCTCG-3’ | 5’-TGAAGTACTCATTATAGTCAAGGGCA-3’ |
| Pparγ | 5’-GTACTGTCGGTTTCAGAAGTGCC-3’ | 5′-ATCTCCGCCAACAGCTTCTCCT-3’ |
| FATP2 | 5’-TCCTCCAAGATGTGCGGTACT-3’ | 5'-TAGGTGAGCGTCTCGTCTCG -3' |
| FATP5 | 5’-GCAGCATGGGTCCTGAAAG-3’ | 5’-ACGGGAGAACTAAGATAGCAGC-3’ |
| Plin2 | 5’-GACCTTGTGTCCTCCGCTTAT-3’ | 5’-CAACCGCAATTTGTGGCTC-3’ |
| Cidec | 5’-ATGGACTACGCCATGAAGTCT-3’ | 5’-CGGTGCTAACACGACAGGG-3’ |
| PNPLA2 | 5’-ATGGACTACGCCATGAAGTCT-3’ | 5’-CGGTGCTAACACGACAGGG-3’ |
| LIPE | 5’-CCAGCCTGAGGGCTTACTG-3’ | 5’-CTCCATTGACTGTGACATCTCG-3’ |
| CPT1a | 5’-CTCCGCCTGAGCCATGAAG-3’ | 5’-CACCAGTGATGATGCCATTCT-3’ |
| Col1a1 | 5’-GCTCCTCTTAGGGGCCACT-3’ | 5’-CCACGTCTCACCATTGGGG-3’ |
| Col3a1 | 5’-CTGTAACATGGAAACTGGGGAAA-3’ | 5’- CCATAGCTGAACTGAAAACCACC-3’ |
| CXCL1 | 5’- ACCGAAGTCATAGCCACACTC-3’ | 5’- CTCCGTTACTTGGGGACACC-3’ |
| CXCL9 | 5’-CCTAGTGATAAGGAATGCACGATG -3’ | 5’-CTAGGCAGGTTTGATCTCCGTTC -3’ |
| CXCL10 | 5’- ATCATCCCTGCGAGCCTATCCT-3’ | 5’-GACCTTTTTTGGCTAAACGCTTTC -3’ |
| IL7 | 5’-CAGGAACTGATAGTAATTGCCCG -3’ | 5’- CTTCAACTTGCGAGCAGCACGA-3’ |
| IFNα | 5’-GGATGTGACCTTCCTCAGACTC-3’ | 5’- ACCTTCTCCTGCGGGAATCCAA-3’ |
| F4/80 | 5'-CCCCAGTGTCCTTACAGAGTG-3' | 5'-GTGCCCAGAGTGGATGTCT-3' |
| ACC1 | 5'-GTTCTGTTGGACAACGCCTTCAC-3' | 5'-GGAGTCACAGAAGCAGCCCATT-3' |
| SCD1 | 5'-GCAAGCTCTACACCTGCCTCTT-3' | 5'-ACGTGCCTTGTAAGTTCTGTGGC-3' |
| IL1β | 5’-AGGTCAAAGGTTTGGAAGCA-3’ | 5’-TGAAGCAGCTATGGCAACTG-3’ |
| IL-6 | 5’-TGGTACTCCAGAAGACCAGAGG-3’ | 5’-AACGATGATGCACTTGCAGA-3’ |
| CD36 | 5’- GGACATTGAGATTCTTTTCCTCTG -3’ | 5’- GCAAAGGCATTGGCTGGAAGAAC-3’ |
| CD31 | 5’-CTGGTGCTCTATGCAAGCCT-3’ | 5’-AGTTGCTGCCCATTCATCAC-3’ |
| Clec4f | 5′-CTTCGGGGAAGCAACAACTC-3 | 5′-CAAGCAACTGCACCAGAGAAC-3′ |
| TNFα | 5’-CACAGAAAGCATGATCCGCGACGT-3 | 5’- CGGCAGAGAGGAGGTTGACTTTCT-3’ |
| Mfn2 | 5’-GTGGAATACGCCAGTGAGAAGC-3’ | 5’-CAACTTGCTGGCACAGATGAGC-3’ |
| β-actin | 5’-CATTGCTGACAGGATGCAGAAGG-3’ | 5’-TGCTGGAAGGTGGACAGTGAGG-3' |
| Ripk3 | 5'-GAAGACACGGCACTCCTTGGTA-3' | 5'-CTTGAGGCAGTAGTTCTTGGTGG-3' |
| CXCL8 | 5'-CATCCAGAGCTTGAGTGTGACG-3' | 5'-GGCTTCAGGGTCAAGGCAAACT-3' |
| TGF-β | 5’-ACCATGCCAACTTCTGTCTGGGAC-3’ | 5’-ACAACTGCTCCACCTTGGGCTTG-3’ |
| Mcm3 | 5’-CTGCTGTCACTACAGACCAGGA-3’ | 5’-ATCACCTCGTGGATGGCTGTGC-3’ |
| Mcm6 | 5’-CGACAGCTTGAGAGCATGATCC-3’ | 5’-TGACATCAGGCGTCTCTACACG-3’ |
| PolE | 5’-CTGAGTTCCTGGGAGACCAGAT-3’ | 5’-CTCACTGTAGGCTCTGCTTGGA-3’ |

**Table S3.** Composition of diet used in the study.

| Product | Control Diet (g) | Control Diet (kcal %) | Western Diet (g) | Western Diet (kcal %) |
|---|---|---|---|---|
| **Protein** | 20 | 20 | 20 | 17 |
| **Carbohydrate** | 65 | 64 | 48 | 40 |
| **Fat** | 7 | 16 | 23 | 43 |
| **Total** | | 100 | | 100 |
| **kcal/gram** | 4 | | 4.8 | |
| **Composition** | **Control Diet (g)** | **Control Diet (kcal)** | **Western Diet (g)** | **Western Diet (kcal)** |
| **Protein** | | | | |
| Casein, Lactic | 200 | 800 | 200 | 800 |
| L-Cystine | 3 | 12 | 0 | 0 |
| DL-Methionine | 0 | 0 | 3 | 12 |
| **Carbohydrate** | | | | |
| Corn Starch | 397.486 | 1590 | 0 | 0 |
| Maltodextrin 10 | 132 | 528 | 0 | 0 |
| Wheat Starch | 0 | 0 | 50 | 200 |
| Sucrose | 107.0777 | 428 | 424 | 1696 |
| Cellulose, BW200 | 50 | 0 | 50 | 0 |
| **Fat** | | | | |
| Soybean Oil | 70 | 630 | 0 | 0 |
| Canola Oil | 0 | 0 | 50 | 450 |
| Lard | 0 | 0 | 181 | 1629 |
| t-Butylhydroquinone | 0.014 | 0 | 0 | 0 |
| Mineral Mix S10026 | 3.5 | 0 | 3.5 | 0 |
| Sodium Chloride | 2.59 | 0 | 2.59 | 0 |
| Calcium Carbonate | 12.495 | 0 | 12.495 | 0 |
| Potassium Phosphate | 6.86 | 0 | 6.86 | 0 |

| | | | | |
|---|---|---|---|---|
| Potassium Citrate | 2.4773 | 0 | 2.4773 | 0 |
| Vitamin Mix V10037 | 10 | 40 | 10 | 40 |
| Choline Bitartrate | 2.5 | 0 | 2.5 | 0 |
| Choline Chloride | 0 | 0 | 2.5 | 0 |
| Cholesterol | 0 | 0 | 1.9 | 0 |
| FD&C Yellow Dye #5 | 0 | 0 | 0 | 0 |
| FD&C Red Dye #40 | 0 | 0 | 0.1 | 0 |
| FD&C Blue Dye #1 | 0 | 0 | 0 | 0 |
| **Total** | 1000 | 4028 | 1000.4223 | 4827 |

**Table S4**: Targeted mitochondrial proteomics data for antioxidant proteins in the livers of Control or *Mlkl*$^{HepKO}$ mice fed either CD or WD for 8 or 15 months

| **Genotype** | | **Control** | **Control** | **Control** | **Control** | **Control** | **Control** | **Control** | **Control** | **Control** | **Control** | ***Mlkl HepKO*** | ***Mlkl HepKO*** | ***Mlkl HepKO*** | ***Mlkl HepKO*** | ***Mlkl HepKO*** | ***Mlkl HepKO*** | ***Mlkl HepKO*** | ***Mlkl HepKO*** | ***Mlkl HepKO*** | ***Mlkl HepKO*** |
|---|---|---|---|---|---|---|---|---|---|---|---|---|---|---|---|---|---|---|---|---|---|
| **Diet** | | **CD** | **CD** | **CD** | **CD** | **CD** | **WD** | **WD** | **WD** | **WD** | **WD** | **CD** | **CD** | **CD** | **CD** | **CD** | **WD** | **WD** | **WD** | **WD** | **WD** |
| **Protein** | **Peptide** | **CD_ 1** | **CD_ 2** | **CD_ 3** | **CD_ 4** | **CD_ 5** | **WD_ 1** | **WD_ 2** | **WD_ 3** | **WD_ 4** | **WD_ 5** | **CD_ 1** | **CD_ 2** | **CD_ 3** | **CD_ 4** | **CD_ 4** | **WD_ 1** | **WD_ 2** | **WD_ 3** | **WD_ 4** | **WD_ 5** |
| **8 Months** | | | | | | | | | | | | | | | | | | | | | |
| Cat | pmol/100ug | 7.223 | 6.251 | 6.657 | 6.678 | 5.993 | 5.172 | 5.223 | 5.477 | 6.456 | 6.278 | 5.602 | 6.856 | 6.085 | 3.675 | 3.154 | 3.374 | 2.987 | 2.954 | 3.309 | 2.694 |
| Gpx1 | pmol/100ug | 1.191 | 1.050 | 1.011 | 1.450 | 1.337 | 0.759 | 0.918 | 0.878 | 0.912 | 0.922 | 1.266 | 0.935 | 1.137 | 0.892 | 0.666 | 0.568 | 0.382 | 0.664 | 0.526 | 0.429 |
| Gpx4 | pmol/100ug | 0.202 | 0.200 | 0.200 | 0.218 | 0.176 | 0.344 | 0.243 | 0.365 | 0.346 | 0.336 | 0.171 | 0.233 | 0.215 | 0.187 | 0.175 | 0.313 | 0.267 | 0.337 | 0.344 | 0.277 |
| Gsr | pmol/100ug | 0.162 | 0.167 | 0.212 | 0.185 | 0.164 | 0.247 | 0.179 | 0.247 | 0.212 | 0.202 | 0.195 | 0.204 | 0.186 | 0.166 | 0.145 | 0.200 | 0.189 | 0.158 | 0.199 | 0.191 |
| Prdx1 | pmol/100ug | 9.947 | 9.557 | 9.828 | 10.010 | 8.944 | 9.183 | 8.712 | 10.082 | 11.324 | 9.193 | 10.564 | 10.802 | 10.834 | 8.225 | 6.282 | 8.310 | 6.434 | 6.948 | 7.288 | 6.032 |
| Prdx2 | pmol/100ug | 1.113 | 1.027 | 1.043 | 0.973 | 0.956 | 1.041 | 0.892 | 1.091 | 1.129 | 1.049 | 0.969 | 0.962 | 1.017 | 0.905 | 0.859 | 1.082 | 0.888 | 0.973 | 1.098 | 0.875 |
| Prdx3 | pmol/100ug | 0.905 | 0.829 | 0.883 | 0.918 | 0.803 | 1.020 | 0.786 | 1.244 | 1.062 | 1.080 | 0.905 | 0.873 | 0.924 | 0.668 | 0.600 | 0.925 | 0.733 | 0.920 | 0.715 | 0.665 |
| Prdx4 | pmol/100ug | 0.269 | 0.246 | 0.235 | 0.310 | 0.246 | 0.232 | 0.183 | 0.303 | 0.238 | 0.208 | 0.328 | 0.285 | 0.237 | 0.255 | 0.208 | 0.221 | 0.240 | 0.224 | 0.227 | 0.163 |
| Prdx5 | pmol/100ug | 1.502 | 1.341 | 1.309 | 1.508 | 1.428 | 1.220 | 0.951 | 1.673 | 1.583 | 1.082 | 1.495 | 1.455 | 1.418 | 1.220 | 1.010 | 1.163 | 0.909 | 1.261 | 1.032 | 0.942 |
| Prdx6 | pmol/100ug | 3.453 | 3.195 | 3.712 | 3.481 | 3.249 | 3.001 | 3.402 | 3.370 | 3.324 | 2.756 | 3.583 | 3.492 | 3.467 | 2.239 | 1.287 | 1.518 | 1.197 | 1.600 | 1.774 | 1.374 |
| Sod1 | pmol/100ug | 8.552 | 7.182 | 6.555 | 7.887 | 7.286 | 3.578 | 5.405 | 4.845 | 4.673 | 4.521 | 7.582 | 7.429 | 7.946 | 4.404 | 3.956 | 3.809 | 3.195 | 3.611 | 3.678 | 2.585 |
| Sod2 | pmol/100ug | 1.120 | 1.111 | 1.054 | 1.275 | 1.089 | 0.831 | 0.778 | 0.811 | 0.902 | 0.805 | 1.103 | 1.027 | 1.167 | 0.966 | 0.609 | 0.623 | 0.376 | 0.513 | 0.547 | 0.463 |
| Txn1 | pmol/100ug | 1.622 | 1.960 | 2.087 | 1.934 | 2.007 | 2.057 | 1.967 | 2.583 | 1.955 | 2.095 | 2.065 | 2.061 | 1.959 | 2.072 | 2.117 | 2.631 | 2.052 | 3.068 | 2.360 | 1.947 |
| Txnrd1 | pmol/100ug | 0.328 | 0.309 | 0.390 | 0.413 | 0.336 | 0.365 | 0.329 | 0.381 | 0.363 | 0.420 | 0.338 | 0.353 | 0.376 | 0.358 | 0.313 | 0.374 | 0.399 | 0.370 | 0.360 | 0.366 |
| **15 months** | | | | | | | | | | | | | | | | | | | | | |
| Cat | pmol/100ug | 4.051 | 4.085 | 4.201 | 3.257 | 3.664 | 4.171 | 4.797 | 3.508 | 4.350 | 3.210 | 4.076 | 4.956 | 3.576 | 3.469 | 3.395 | 2.978 | 4.410 | 2.754 | 2.566 | 1.978 |
| Gpx1 | pmol/100ug | 0.523 | 0.630 | 0.653 | 0.661 | 0.592 | 0.555 | 0.447 | 0.449 | 0.408 | 0.546 | 0.796 | 0.728 | 0.579 | 0.714 | 0.748 | 0.373 | 0.403 | 0.503 | 0.475 | 0.491 |
| Gpx4 | pmol/100ug | 0.218 | 0.236 | 0.189 | 0.230 | 0.335 | 0.326 | 0.322 | 0.405 | 0.367 | 0.203 | 0.281 | 0.238 | 0.222 | 0.202 | 0.209 | 0.316 | 0.339 | 0.340 | 0.348 | 0.301 |
| Gsr | pmol/100ug | 0.178 | 0.195 | 0.157 | 0.202 | 0.270 | 0.228 | 0.254 | 0.305 | 0.324 | 0.178 | 0.205 | 0.176 | 0.168 | 0.174 | 0.212 | 0.174 | 0.277 | 0.145 | 0.180 | 0.160 |
| Prdx1 | pmol/100ug | 6.654 | 7.853 | 7.126 | 6.458 | 7.872 | 8.675 | 6.402 | 8.054 | 6.908 | 6.550 | 10.519 | 8.736 | 8.133 | 9.837 | 7.358 | 7.753 | 7.427 | 7.967 | 6.328 | 6.831 |
| Prdx2 | pmol/100ug | 0.837 | 0.981 | 0.985 | 0.931 | 1.045 | 1.270 | 0.986 | 1.325 | 0.949 | 0.990 | 1.301 | 1.177 | 1.026 | 1.177 | 1.000 | 1.361 | 1.026 | 1.222 | 1.004 | 1.041 |
| Prdx3 | pmol/100ug | 0.627 | 0.777 | 0.591 | 0.617 | 0.882 | 1.151 | 0.841 | 1.007 | 0.919 | 0.584 | 0.848 | 0.708 | 0.721 | 0.781 | 0.527 | 0.924 | 0.855 | 0.803 | 0.685 | 0.684 |
| Prdx4 | pmol/100ug | 0.146 | 0.224 | 0.177 | 0.203 | 0.268 | 0.178 | 0.197 | 0.240 | 0.176 | 0.177 | 0.323 | 0.251 | 0.228 | 0.230 | 0.213 | 0.212 | 0.187 | 0.228 | 0.189 | 0.258 |
| Prdx5 | pmol/100ug | 0.880 | 1.199 | 1.183 | 1.063 | 1.428 | 1.240 | 1.073 | 1.065 | 0.748 | 0.700 | 1.018 | 0.974 | 0.789 | 0.995 | 0.964 | 0.798 | 0.898 | 1.003 | 0.730 | 0.702 |
| Prdx6 | pmol/100ug | 1.584 | 2.137 | 1.860 | 1.708 | 2.061 | 1.836 | 1.656 | 1.276 | 1.120 | 1.117 | 1.586 | 1.602 | 1.472 | 1.391 | 1.404 | 1.323 | 1.322 | 1.379 | 0.936 | 1.157 |
| Sod1 | pmol/100ug | 5.353 | 4.626 | 5.279 | 4.931 | 3.909 | 4.601 | 3.524 | 2.554 | 1.784 | 2.276 | 3.394 | 2.392 | 3.134 | 3.119 | 2.882 | 1.984 | 1.887 | 2.507 | 2.048 | 2.012 |
| Sod2 | pmol/100ug | 0.608 | 0.687 | 0.655 | 0.770 | 0.547 | 0.688 | 0.610 | 0.604 | 0.478 | 0.554 | 0.790 | 0.787 | 0.636 | 0.682 | 0.785 | 0.452 | 0.498 | 0.469 | 0.362 | 0.450 |
| Txn1 | pmol/100ug | 1.756 | 1.942 | 1.953 | 1.752 | 3.228 | 3.054 | 2.582 | 2.626 | 2.840 | 1.614 | 2.127 | 2.099 | 1.770 | 1.712 | 1.740 | 2.269 | 2.633 | 2.332 | 2.105 | 1.724 |

Table S5. Untargeted Lipidomic profiling of hepatic tissues (Mlkl$^{HepKO}$ WD vs. Control WD)

| Genotype | | Control | Control | Control | Control | Control | Control | Control | Control | *Mlkl HepKO* | *Mlkl HepKO* | *Mlkl HepKO* | *Mlkl HepKO* | *Mlkl HepKO* | *Mlkl HepKO* | *Mlkl HepKO* | *Mlkl HepKO* | | | |
|---|---|---|---|---|---|---|---|---|---|---|---|---|---|---|---|---|---|---|---|---|
| **Diet** | | **CD** | **CD** | **CD** | **CD** | **WD** | **WD** | **WD** | **WD** | **CD** | **CD** | **CD** | **CD** | **WD** | **WD** | **WD** | **WD** | | | |
| **Lipid Species** | **Class** | **CD_1** | **CD_2** | **CD_3** | **CD_4** | **WD_1** | **WD_2** | **WD_3** | **WD_4** | **CD_1** | **CD_2** | **CD_3** | **CD_4** | **WD_1** | **WD_2** | **WD_3** | **WD_4** | **lgFold** | **p_value** | **q_value** |
| PE(34:3e)-H | PE | 51.62302276 | 51.66611253 | 58.70645469 | 53.86685419 | 15.76987028 | 11.73105255 | 8.838234859 | 9.94825536 | 44.35793173 | 49.8750799 | 57.48872029 | 44.59531904 | 18.26085601 | 17.26017451 | 22.65781558 | 17.43861211 | -1.49 | 3.40E-09 | 1.29E-06 |
| PE(44:12)-H | PE | 133.0463408 | 161.2433868 | 165.9729704 | 168.2081033 | 16.14587682 | 12.76109834 | 12.20227099 | 37.85171742 | 139.0986822 | 177.8326233 | 223.2980131 | 158.7908145 | 52.20364762 | 45.78521032 | 40.60953239 | 42.14838513 | -2.13 | 3.60E-08 | 3.89E-06 |
| PE(36:6)+H | PE | 9.584591791 | 12.92321762 | 15.96014204 | 22.09168654 | 2.11634266 | 1.164804217 | 1.623166055 | 0.508095514 | 9.720631659 | 8.806066815 | 10.36085118 | 11.81595905 | 1.976851938 | 4.416920067 | 3.153016283 | 2.399494328 | -1.92 | 7.40E-08 | 5.74E-06 |
| MePC(34:7)+Na | MePC | 8.846285926 | 10.1275863 | 17.3784816 | 12.08766113 | 4.049197314 | 3.373816632 | 3.205189826 | 3.875637539 | 9.446386445 | 11.09287311 | 9.71788639 | 11.26985182 | 4.927034668 | 6.572149372 | 5.371205575 | 4.009185443 | -1.02 | 1.60E-06 | 3.22E-05 |
| PE(38:7)+Na | PE | 8.846285926 | 10.1275863 | 17.3784816 | 12.08766113 | 4.049197314 | 3.373816632 | 3.205189826 | 3.875637539 | 9.446386445 | 11.09287311 | 9.71788639 | 11.26985182 | 4.927034668 | 6.572149372 | 5.371205575 | 4.009185443 | -1.02 | 1.60E-06 | 3.22E-05 |
| PI(36:5)-H | PI | 53.51180737 | 64.66055464 | 47.87645018 | 72.47275534 | 10.66374373 | 4.875567086 | 0.512855586 | 5.111920378 | 67.48600203 | 28.5332034 | 60.32638734 | 69.95364955 | 23.50761141 | 18.92578268 | 14.53398721 | 12.38816035 | -2.48 | 4.10E-06 | 5.69E-05 |
| MePC(33:2)+NH4 | MePC | 20.83862565 | 20.84202135 | 28.28020221 | 39.98403464 | 7.829264466 | 2.924684341 | 5.064056753 | 8.104873476 | 14.80638401 | 28.49248254 | 20.21707829 | 22.56637713 | 9.988642683 | 11.15055624 | 10.84384461 | 7.581331724 | -1.43 | 5.80E-06 | 7.14E-05 |
| SM(t42:1)+H | SM | 25.99413609 | 24.97690318 | 22.6669013 | 27.31477295 | 9.030044495 | 13.97298349 | 12.42694708 | 12.85611428 | 18.34677964 | 24.98725004 | 23.32458296 | 19.07366434 | 15.91002199 | 17.75857306 | 14.15990446 | 14.14181887 | -0.71 | 9.60E-06 | 9.89E-05 |
| PC(32:1)+H | PC | 14.00475879 | 15.38038749 | 10.90643017 | 16.1240586 | 4.366156326 | 5.891713586 | 4.073280292 | 6.136492549 | 11.3645031 | 11.25030228 | 9.764132035 | 11.91710503 | 5.598053187 | 10.74108402 | 10.05557089 | 6.595187386 | -0.91 | 1.30E-05 | 0.000126 |
| PC(30:0)+H | PC | 123.2288284 | 144.87695 | 132.3465378 | 136.2213752 | 73.8413461 | 87.10297341 | 82.90333619 | 82.80023587 | 111.2927809 | 109.1172361 | 123.3861647 | 120.7502663 | 109.8956855 | 99.07779795 | 108.1263002 | 89.41669181 | -0.49 | 1.60E-05 | 0.000143 |
| PC(33:3)+H | PC | 823.3358137 | 876.0924001 | 1304.509588 | 1227.741015 | 505.4401958 | 594.0662536 | 469.0516944 | 469.8441802 | 799.8295507 | 984.1402922 | 882.7017849 | 978.0143664 | 641.6238307 | 835.2150356 | 726.2315516 | 608.1411527 | -0.71 | 3.20E-05 | 0.000246 |
| PE(34:0)+Na | PE | 823.3358137 | 876.0924001 | 1304.509588 | 1227.741015 | 505.4401958 | 594.0662536 | 469.0516944 | 469.8441802 | 799.8295507 | 984.1402922 | 882.7017849 | 978.0143664 | 641.6238307 | 835.2150356 | 726.2315516 | 608.1411527 | -0.71 | 3.20E-05 | 0.000246 |
| PE(34:0)+Na | PE | 823.3358137 | 876.0924001 | 1304.509588 | 1227.741015 | 505.4401958 | 594.0662536 | 469.0516944 | 469.8441802 | 799.8295507 | 984.1402922 | 882.7017849 | 978.0143664 | 641.6238307 | 835.2150356 | 726.2315516 | 608.1411527 | -0.71 | 3.20E-05 | 0.000246 |
| PE(36:3)+H | PE | 823.3358137 | 876.0924001 | 1304.509588 | 1227.741015 | 505.4401958 | 594.0662536 | 469.0516944 | 469.8441802 | 799.8295507 | 984.1402922 | 882.7017849 | 978.0143664 | 641.6238307 | 835.2150356 | 726.2315516 | 608.1411527 | -0.71 | 3.20E-05 | 0.000246 |
| DG(34:1)+NH4 | DG | 0.057770475 | 0.632403385 | 0.315366937 | 0.370638724 | 3.610202386 | 2.307882673 | 2.063998423 | 1.48818107 | 0.199946947 | 0.526142877 | 0.20487833 | 0.300699658 | 1.15452621 | 1.370563974 | 0.372005887 | 0.616003713 | 0.91 | 3.20E-05 | 0.00025 |
| PC(31:1)+H | PC | 64.52101716 | 41.63174973 | 83.66380459 | 72.31451029 | 19.85429344 | 34.01288824 | 19.48605749 | 15.38833149 | 60.3800157 | 58.78080911 | 58.84104034 | 45.10600199 | 26.38363389 | 44.59139979 | 30.34590914 | 32.41142877 | -1.06 | 3.40E-05 | 0.000259 |
| PE(34:1)+H | PE | 64.52101716 | 41.63174973 | 83.66380459 | 72.31451029 | 19.85429344 | 34.01288824 | 19.48605749 | 15.38833149 | 60.3800157 | 58.78080911 | 58.84104034 | 45.10600199 | 26.38363389 | 44.59139979 | 30.34590914 | 32.41142877 | -1.06 | 3.40E-05 | 0.000259 |
| PE(34:2)+Na | PE | 19.94286621 | 23.2536852 | 36.00005192 | 25.97136088 | 13.38942766 | 14.40810484 | 9.957363701 | 12.89447559 | 20.63589337 | 22.18657596 | 24.42744243 | 24.95031022 | 19.04342215 | 21.85691571 | 17.70058716 | 16.74442957 | -0.68 | 6.10E-05 | 0.000415 |
| PEt(36:4)-H | PEt | 38.39297334 | 14.9805004 | 39.63405035 | 41.42260565 | 6.590268996 | 15.32000578 | 6.239258224 | 4.178234155 | 31.01080411 | 28.8408389 | 34.49022023 | 22.97108647 | 17.70298088 | 17.57406829 | 11.53208308 | 19.08827428 | -1.36 | 8.20E-05 | 0.000523 |
| PG(36:2)-H | PG | 0.213118943 | 0 | 0 | 0 | 0.540638751 | 2.027371518 | 1.304624395 | 0.98332672 | 0.184649674 | 0.082644473 | 0.104844663 | 0.075685887 | 0.223517934 | 0.365846293 | 0.855805969 | 0.203934014 | 0.72 | 1.00E-04 | 0.000639 |
| PE(36:2)-H | PE | 249.9692176 | 232.2001135 | 238.6439227 | 598.626681 | 14.31903738 | 53.63757032 | 86.4590465 | 101.7917136 | 225.8099759 | 177.4024864 | 202.2175182 | 209.5167698 | 126.1708125 | 94.99554995 | 114.5445655 | 101.5817788 | -1.75 | 0.00012 | 0.000729 |
| dMePE(34:2)-H | dMePE | 249.9692176 | 232.2001135 | 238.6439227 | 598.626681 | 14.31903738 | 53.63757032 | 86.4590465 | 101.7917136 | 225.8099759 | 177.4024864 | 202.2175182 | 209.5167698 | 126.1708125 | 94.99554995 | 114.5445655 | 101.5817788 | -1.75 | 0.00012 | 0.000729 |
| ChE(17:1)+NH4 | ChE | 0.827111024 | 0.923165933 | 1.426694622 | 1.611972994 | 2.306623564 | 5.708585817 | 6.857314786 | 8.889528459 | 0.663723764 | 0.210315934 | 0.735379063 | 0.468749906 | 0.670884432 | 2.280360885 | 1.276744287 | 1.221634296 | 1.09 | 0.00012 | 0.000729 |
| PA(38:4)-H | PA | 38.39297334 | 14.9805004 | 39.63405035 | 41.42260565 | 5.030923794 | 15.32000578 | 6.239258224 | 2.026416825 | 31.01080411 | 28.8408389 | 34.49022023 | 22.97108647 | 17.70298088 | 17.57406829 | 11.53208308 | 19.08827428 | -1.56 | 0.00013 | 0.000772 |
| PG(44:11)+NH4 | PG | 5.467538976 | 5.376336992 | 5.886661265 | 7.567907879 | 1.063264312 | 2.384687277 | 2.301462432 | 3.695703282 | 3.775846336 | 3.872412663 | 5.563697634 | 4.525313201 | 4.673056958 | 3.897756085 | 4.793254821 | 3.19023785 | -0.78 | 0.00015 | 0.000871 |
| PS(42:6)+Na | PS | 5.467538976 | 5.376336992 | 5.886661265 | 7.567907879 | 1.063264312 | 2.384687277 | 2.301462432 | 3.695703282 | 3.775846336 | 3.872412663 | 5.563697634 | 4.525313201 | 4.673056958 | 3.897756085 | 4.793254821 | 3.19023785 | -0.78 | 0.00015 | 0.000871 |
| PA(36:3)-H | PA | 180.7232773 | 165.8423776 | 174.3347517 | 175.5171289 | 32.48970182 | 47.04607019 | 65.99751252 | 22.52264433 | 171.5179658 | 190.3824911 | 147.5202295 | 115.7796416 | 194.178241 | 66.15167776 | 196.3686653 | 47.82840357 | -1.48 | 0.00021 | 0.00114 |
| TG(34:2e)+NH4 | TG | 0.257429454 | 0.556301031 | 0.325567261 | 0.15163945 | 1.990852549 | 2.421236124 | 1.810308616 | 2.029919568 | 0.018014434 | 0.489154592 | 0.067303399 | 0.208497359 | 0.175466191 | 2.01325823 | 0.244295754 | 0.541651899 | 0.84 | 0.00024 | 0.00126 |
| PG(36:2)-H | PG | 0 | 0 | 0 | 0.357006654 | 2.9145686 | 1.164959337 | 6.546147427 | 0.87216442 | 0 | 0 | 0 | 0 | 0 | 0.449836163 | 0.85954187 | 0.71495022 | 1.12 | 4.00E-04 | 0.00197 |

| | | | | | | | | | | | | | | | | | | | | |
|---|---|---|---|---|---|---|---|---|---|---|---|---|---|---|---|---|---|---|---|---|
| TG(56:6)+H | TG | 2.00684484 | 0.793909145 | 0.77288178 | 0.317196441 | 15.80039259 | 5.821985839 | 10.53620945 | 5.709805331 | 1.346231704 | 2.43234687 | 1.867737459 | 0.722360742 | 5.869734821 | 1.666461195 | 0.237819188 | 6.710493915 | 1.64 | 0.00045 | 0.00217 |
| PE(38:5p)+H | PE | 10.79502696 | 13.62827814 | 11.32086009 | 15.43962049 | 3.732822559 | 4.415326002 | 4.09803982 | 8.708250931 | 6.592766034 | 7.727824563 | 12.94017745 | 9.727679284 | 11.76849268 | 9.364455866 | 15.28336735 | 7.795971184 | -0.83 | 0.00047 | 0.00226 |
| PE(38:4)-H | PE | 3.335761652 | 11.17171316 | 10.26153505 | 2.919838025 | 20.24843082 | 17.72162915 | 25.1197973 | 35.84802391 | 11.63631773 | 6.541357099 | 9.023095287 | 8.78709439 | 5.305225029 | 6.780615737 | 15.39201457 | 12.05807824 | 1.28 | 0.00049 | 0.00233 |
| PE(36:3e)+Na | PE | 14.19204794 | 8.566638371 | 9.791495506 | 11.60717333 | 3.732822559 | 4.415326002 | 4.09803982 | 8.708250931 | 6.592766034 | 7.727824563 | 9.694221131 | 9.9792845 | 11.76849268 | 9.364455866 | 9.499009787 | 7.795971184 | -0.69 | 0.00076 | 0.00335 |
| TG(38:4e)+H | TG | 13.60381497 | 11.60552276 | 18.44231367 | 15.160062 | 5.406516332 | 3.737520538 | 5.605012982 | 10.27722152 | 15.10581703 | 17.56751053 | 15.11479004 | 14.42076799 | 17.34104956 | 18.16223887 | 16.20161936 | 8.049931816 | -0.81 | 0.00076 | 0.00335 |
| Cer(d30:0)+H | Cer | 0.090035582 | 0.26764828 | 0.300869063 | 0.17115532 | 0.897485788 | 1.064429023 | 0.702861659 | 0.486065759 | 0.288083816 | 0.126230216 | 0.200758638 | 0.119061944 | 0.087392733 | 0.491216701 | 0.24610762 | 0.163386208 | 0.39 | 0.0011 | 0.00446 |
| TG(50:3)+NH4 | TG | 0.387900004 | 0.604024402 | 0.650337522 | 0.365661622 | 1.253439762 | 1.114567376 | 4.190181762 | 3.141153322 | 0.256319559 | 0.451464472 | 0.084697728 | 0.973571385 | 0.357724533 | 0.443589013 | 0.687352609 | 0.229111697 | 0.75 | 0.0013 | 0.00537 |
| PA(43:3)-H | PA | 7.523289145 | 2.692812434 | 2.91997756 | 15.24742346 | 9.349657456 | 55.49924127 | 50.46940692 | 36.2897104 | 15.62845835 | 7.584983022 | 8.634002947 | 5.195084741 | 11.71809492 | 19.04677878 | 12.23925818 | 7.870226487 | 1.58 | 0.0013 | 0.00543 |
| MG(29:1)+NH4 | MG | 1.437875916 | 0.274738318 | 1.540207248 | 0.74589285 | 2.331500041 | 3.894359387 | 3.200554633 | 3.508630557 | 1.201811069 | 1.499533965 | 2.886333855 | 0.524427439 | 1.407992569 | 1.170352507 | 0.705914785 | 1.631561027 | 0.78 | 0.0014 | 0.00561 |
| PC(36:3)+H | PC | 4.928266118 | 3.835251143 | 2.409280969 | 12.78321458 | 16.36653452 | 143.8354 | 12.62199602 | 333.537277 | 5.233395101 | 4.912558353 | 3.635065108 | 15.88044877 | 8.883097822 | 8.799022341 | 14.53693886 | 4.849047011 | 2.26 | 0.0023 | 0.00864 |
| LPC(16:0)+H | LPC | 2.834934912 | 2.056728645 | 4.626686733 | 3.101261385 | 0.690917099 | 0.17254213 | 0.234973507 | 1.250460825 | 0.673070409 | 0.723142713 | 3.979399185 | 2.783871292 | 1.010349907 | 3.311282852 | 3.411005015 | 1.410081103 | -0.97 | 0.0027 | 0.0102 |
| MG(28:1)+H | MG | 3.016066122 | 1.991707839 | 1.864676307 | 4.997405466 | 1.291443194 | 1.066408774 | 1.355069165 | 1.342278414 | 2.751442129 | 1.697823405 | 1.514320785 | 2.357069556 | 2.050979823 | 2.084579372 | 2.394335327 | 2.68142569 | -0.52 | 0.0032 | 0.0118 |
| TG(46:2)+NH4 | TG | 0.515536101 | 1.098575294 | 0.703665535 | 0.460748907 | 3.772562553 | 31.23728919 | 3.81429339 | 1.765898315 | 1.588643851 | 0.46698233 | 0.732813565 | 0.636804335 | 0.480628679 | 1.080160269 | 2.220665645 | 0.58868884 | 1.39 | 0.0033 | 0.0119 |
| TG(46:7)+H | TG | 26.39800501 | 27.26548478 | 23.50353375 | 20.67976211 | 16.15493769 | 16.61391385 | 18.79050019 | 10.97551901 | 17.15261198 | 22.08770221 | 22.22429766 | 20.01765352 | 18.11222296 | 22.16167344 | 23.97371069 | 29.16936979 | -0.44 | 0.0035 | 0.0126 |
| PE(44:11)-H | PE | 8.393872165 | 4.587490066 | 6.307410513 | 17.1884377 | 2.416437492 | 3.230013011 | 4.80762189 | 1.550018403 | 6.122541973 | 3.975234093 | 2.103212813 | 6.273419382 | 9.896532217 | 7.458538049 | 10.34352691 | 10.09869308 | -0.87 | 0.004 | 0.0141 |
| MG(29:1)+H | MG | 8.829985499 | 7.549028235 | 5.45331666 | 1.778645881 | 1.415714691 | 0.438338361 | 2.534845556 | 1.493283115 | 2.632804999 | 8.471005671 | 3.424016098 | 2.623235043 | 4.208422215 | 2.997578864 | 2.543765127 | 4.497727194 | -0.97 | 0.0042 | 0.0146 |
| PG(38:4)-H | PG | 10.7235598 | 12.6141562 | 20.82970524 | 19.40965785 | 3.053279214 | 6.356929069 | 4.413157271 | 14.78569071 | 12.2372279 | 15.55183135 | 9.532918324 | 10.19557775 | 15.7663948 | 17.83678066 | 24.98789403 | 16.97708021 | -0.83 | 0.0043 | 0.0148 |
| MG(18:0)+H | MG | 2.435910874 | 1.594193335 | 1.97431186 | 2.64087715 | 8.805022736 | 10.13274037 | 6.448571236 | 4.787637132 | 8.546111794 | 6.008094879 | 4.177514067 | 0.773715002 | 2.478276833 | 3.085164663 | 1.135167209 | 2.417075107 | 0.97 | 0.0045 | 0.0155 |
| MePC(32:3e)+Na | MePC | 13.46825453 | 8.566638371 | 9.791495506 | 12.81815982 | 3.732822559 | 4.415326002 | 8.190797983 | 8.708250931 | 6.592766034 | 7.727824563 | 9.694221131 | 9.9792845 | 11.76849268 | 9.364455866 | 9.499009787 | 7.795971184 | -0.55 | 0.0056 | 0.0185 |
| TG(34:1e)+Na | TG | 1.538920413 | 0.08539772 | 0.466392527 | 0.13590475 | 33.33276263 | 49.55021932 | 1.738057574 | 1.526804532 | 0.160318039 | 0.529878626 | 0.822282047 | 4.705130475 | 1.211140576 | 2.271437353 | 0.872165042 | 1.102561572 | 1.97 | 0.0056 | 0.0186 |
| PC(31:1)+H | PC | 147.3975522 | 192.9298492 | 219.8194 | 202.9968997 | 125.1312238 | 151.9749952 | 122.5850449 | 150.8496546 | 169.8536325 | 191.3535753 | 174.1260458 | 164.339259 | 167.8963404 | 168.649437 | 197.9483151 | 155.816036 | -0.32 | 0.006 | 0.0196 |
| PE(34:1)+H | PE | 147.3975522 | 192.9298492 | 219.8194 | 202.9968997 | 125.1312238 | 151.9749952 | 122.5850449 | 150.8496546 | 169.8536325 | 191.3535753 | 174.1260458 | 164.339259 | 167.8963404 | 168.649437 | 197.9483151 | 155.816036 | -0.32 | 0.006 | 0.0196 |
| PE(38:6e)+H | PE | 5.930461219 | 8.566638371 | 9.791495506 | 12.50149428 | 3.732822559 | 4.415326002 | 4.09803982 | 8.708250931 | 6.592766034 | 7.727824563 | 9.694221131 | 8.890522912 | 9.417298952 | 9.364455866 | 9.499009787 | 7.795971184 | -0.51 | 0.0081 | 0.0253 |
| dMePE(34:2)-H | dMePE | 1.574282109 | 1.029864282 | 1.730451579 | 1.21375507 | 0.451782721 | 0.942010762 | 0.220417203 | 0.485279598 | 0.649635066 | 1.820182164 | 1.505924905 | 0.36624995 | 1.246576173 | 1.311713572 | 1.516215192 | 1.770880945 | -0.46 | 0.0099 | 0.03 |
| MePC(33:0)+Na | MePC | 66.61017536 | 43.39451304 | 40.32286053 | 78.08318317 | 20.05377115 | 42.71451206 | 29.7614339 | 25.71839147 | 36.66485485 | 63.15530746 | 34.671235 | 54.36727073 | 35.037067 | 76.03730825 | 75.42221174 | 55.15431386 | -0.64 | 0.011 | 0.0328 |
| PMe(19:1e)-H | PMe | 10.37772347 | 15.14599773 | 9.588891279 | 10.56125116 | 14.4946688 | 15.98446075 | 17.32157489 | 15.17613089 | 13.99694929 | 10.74905233 | 11.10872949 | 11.90036741 | 10.65627952 | 13.92710545 | 9.691626272 | 10.99856282 | 0.31 | 0.011 | 0.0336 |
| dMePE(36:4)-H | dMePE | 3.335761652 | 9.062757172 | 9.183897332 | 8.949337027 | 12.27227068 | 12.57723775 | 26.74877699 | 32.96614264 | 16.31800256 | 11.35356723 | 9.302973912 | 11.74091634 | 13.15105608 | 5.971833848 | 19.91112132 | 4.189939609 | 0.91 | 0.012 | 0.0341 |
| TG(52:2)+Na | TG | 2.846564157 | 2.458023272 | 0.755858496 | 1.078116856 | 5.712381874 | 1.732106662 | 5.973085728 | 7.32704252 | 4.774039093 | 1.297196452 | 3.018186077 | 1.083253555 | 1.417579891 | 1.742852352 | 2.254938035 | 1.713167409 | 0.77 | 0.013 | 0.0381 |
| TG(48:1)+NH4 | TG | 1.18820522 | 1.782765851 | 1.283463117 | 0.890355123 | 3.026161225 | 5.220879387 | 4.658123028 | 1.58678857 | 2.30798387 | 1.849004853 | 1.206624792 | 0.98976059 | 3.076929974 | 0.419663768 | 0.616974607 | 1.383877221 | 0.66 | 0.013 | 0.0383 |
| PE(40:6p)+H | PE | 9.495308348 | 6.361404482 | 7.43960419 | 7.633715766 | 2.74582096 | 2.890780942 | 4.024148831 | 2.458185197 | 17.19848691 | 4.392379041 | 2.620061363 | 1.907880326 | 6.558918506 | 10.835478 | 8.945972461 | 5.992971343 | -0.78 | 0.021 | 0.0551 |
| dMePE(38:6)-H | dMePE | 1.30359684 | 1.620161873 | 13.94392918 | 5.040237617 | 5.268687444 | 193.4773713 | 178.775502 | 6.240268222 | 4.551701024 | 9.587044286 | 2.031091841 | 23.50376887 | 5.15219641 | 3.778834535 | 4.655731163 | 1.672822514 | 1.99 | 0.021 | 0.0566 |
| MePC(19:4e)+Na | MePC | 17.45331536 | 5.753338134 | 15.20988766 | 8.041965927 | 1.490367634 | 3.704576909 | 1.732893102 | 8.416523418 | 3.520948112 | 2.458130986 | 19.33904161 | 3.432702121 | 7.036157721 | 21.62293266 | 9.433519345 | 10.57224548 | -1.03 | 0.023 | 0.0606 |

| | | | | | | | | | | | | | | | | | | | | |
|---|---|---|---|---|---|---|---|---|---|---|---|---|---|---|---|---|---|---|---|---|
| MG(18:0)+Na | MG | 92.02569228 | 216.4393233 | 256.8216233 | 222.3548563 | 113.6625911 | 31.5708872 | 62.49013004 | 112.4428931 | 272.1793665 | 49.54301141 | 290.3595688 | 102.4534675 | 330.7191092 | 339.012437 | 282.4707644 | 369.8579333 | -0.94 | 0.027 | 0.0679 |
| PE(45:7)+Na | PE | 5.820830587 | 5.126357335 | 6.920940327 | 8.095701835 | 4.431672725 | 4.641412993 | 4.599667978 | 4.605273874 | 4.749740688 | 5.981000823 | 6.277957908 | 3.224913629 | 5.733200174 | 7.04774569 | 6.767706533 | 5.785738803 | -0.29 | 0.032 | 0.0794 |
| PC(38:6)+H | PC | 136.1800384 | 296.8301176 | 367.9914295 | 257.1988236 | 111.973374 | 135.5611506 | 118.2929506 | 197.1688786 | 198.7326449 | 262.7589189 | 251.5616341 | 226.1161596 | 532.0039446 | 504.4725992 | 423.726778 | 163.0506494 | -0.59 | 0.036 | 0.0877 |
| MG(27:1)+H | MG | 0.781373267 | 0.916886048 | 0.7539837 | 2.38047204 | 0.205080758 | 0.120602357 | 0.093132098 | 0.210266936 | 0.615403722 | 0.544291652 | 0.385034804 | 0.193070584 | 0.560395522 | 0.196711892 | 4.850994017 | 1.813798356 | -0.61 | 0.039 | 0.0927 |
| TG(61:1)+NH4 | TG | 10.52872251 | 8.852227172 | 10.08996327 | 10.89668369 | 7.648858649 | 7.626704535 | 8.749499667 | 8.803955047 | 9.598039797 | 8.783677427 | 8.976685802 | 10.2543569 | 11.59501312 | 11.93781964 | 9.095051801 | 9.958357766 | -0.19 | 0.039 | 0.0937 |
| dMePE(18:0e)-H | dMePE | 2130.913567 | 2129.795787 | 1507.491238 | 1716.954887 | 3880.56007 | 4329.181054 | 4381.89243 | 3992.065133 | 4336.801001 | 3321.468307 | 3603.13008 | 3320.361799 | 3281.293699 | 3638.90531 | 3016.147576 | 3060.262058 | 0.8 | 1.10E-06 | 2.55E-05 |
| dMePE(40:4)-H | dMePE | 102.9769213 | 122.3501689 | 129.1536173 | 150.5768307 | 12.06420861 | 25.38465521 | 6.661448596 | 12.12180619 | 51.96745078 | 105.1396687 | 22.43979716 | 36.65095163 | 54.27709003 | 48.36103426 | 75.64488757 | 31.38341338 | -2.22 | 4.10E-06 | 5.73E-05 |
| TG(50:1)+Na | TG | 21.63077057 | 19.87615048 | 25.63431928 | 19.72648855 | 1.544545506 | 1.512157527 | 2.34857102 | 1.16546371 | 2.927357142 | 9.922192608 | 4.447063728 | 8.19841061 | 1.465726402 | 14.21050427 | 5.123565606 | 4.407298618 | -2.16 | 6.20E-06 | 7.41E-05 |
| TG(50:2)+NH4 | TG | 9.91420223 | 13.71709207 | 12.42734878 | 12.85528111 | 52.95448857 | 36.42229152 | 121.9286819 | 60.38781854 | 48.51993884 | 34.81502512 | 26.8093525 | 36.07905707 | 48.70825154 | 37.88564988 | 48.20926198 | 12.38831738 | 1.56 | 8.60E-05 | 0.00055 |
| PE(38:7)-H | PE | 305.6703028 | 247.3345113 | 417.3784326 | 295.9131368 | 6.240596481 | 27.97667423 | 43.78550717 | 11.82338717 | 7.231167375 | 83.13534494 | 278.71606 | 25.68977355 | 150.5660982 | 151.9337692 | 162.6014754 | 172.6149707 | -2.82 | 0.00025 | 0.00132 |
| PC(36:3)+H | PC | 1.332572444 | 0.22774027 | 0.516204592 | 0.424985229 | 0.871124378 | 2.086161385 | 4.79156751 | 1.700463673 | 1.439210057 | 1.676021197 | 2.333296048 | 2.278744594 | 0.808440904 | 0.05403066 | 0.516054106 | 0.152060367 | 0.67 | 0.0072 | 0.0229 |
| PC(34:2)+H | PC | 3.419656861 | 7.336618525 | 2.253334961 | 6.50086323 | 0.737486157 | 1.141462981 | 2.821006295 | 0.820610019 | 0.387796258 | 1.662875639 | 3.509314401 | 0.523369185 | 3.549121334 | 2.416793241 | 6.822907627 | 2.007401859 | -0.89 | 0.011 | 0.0332 |
| PS(28:1e)+H | PS | 2.813909167 | 6.162642779 | 5.029386683 | 7.964156643 | 0.678785185 | 3.760564717 | 2.584802066 | 1.681161634 | 1.20251668 | 1.290268661 | 0.685841576 | 1.47429344 | 2.275697477 | 6.340896536 | 10.81465485 | 9.039206359 | -0.74 | 0.019 | 0.0518 |
| PC(38:4)+H | PC | 2.990830486 | 21.95735448 | 4.470779225 | 6.710669704 | 2.596334855 | 2.476186564 | 1.434933616 | 2.019383669 | 1.235832245 | 0.735818507 | 2.289665229 | 9.486345673 | 4.313720996 | 6.504699945 | 7.35180642 | 11.59710506 | -0.93 | 0.032 | 0.0787 |
| MG(28:1)+H | MG | 0.733103718 | 0.67672533 | 1.206671493 | 2.249657125 | 0.053981071 | 0.480921116 | 0.297265675 | 0.74003375 | 0.289725719 | 0.150481418 | 0.524744949 | 0.631748259 | 1.000510632 | 1.143440398 | 3.581625434 | 2.283463112 | -0.44 | 0.038 | 0.0923 |
| PC(40:4)+HCOO | PC | 156.3586547 | 23.19901389 | 4.678759286 | 21.32863878 | 181.509774 | 118.9278589 | 175.1654886 | 44.45683568 | 134.716715 | 120.0332774 | 101.8583965 | 119.6884203 | 17.86605308 | 106.8616166 | 7.02940141 | 6.71023292 | 1.47 | 0.04 | 0.0944 |

**Note:** Values represent absolute abundance metrics. Adjusted p-values (q- values) less than 0.05 denote statistical significance.

**Table S6**: Targeted mitochondrial proteomics data for electron transport chain complex I-V proteins in the livers of Control or *Mlkl*$^{HepKO}$ mice fed either CD or WD for 15 months

| Protein | Peptide | Control - Control diet | | | | | Control - Western diet | | | | | *MlklHepKO* - Control diet | | | | | *MlklHepKO* - Western diet | | | | |
|---|---|---|---|---|---|---|---|---|---|---|---|---|---|---|---|---|---|---|---|---|---|
| Atp5f1a | pmol/100ug | 4.86 | 7.45 | 6.68 | 4.51 | 6.79 | 5.26 | 4.95 | 5.65 | 6.39 | 6.27 | 5.90 | 7.40 | 7.31 | 5.42 | 6.41 | 4.65 | 5.90 | 5.57 | 4.74 | 4.50 |
| Atp5f1b | pmol/100ug | 5.21 | 7.47 | 7.38 | 5.16 | 6.27 | 4.67 | 6.05 | 6.16 | 6.67 | 6.20 | 5.85 | 7.09 | 8.57 | 4.33 | 6.87 | 5.18 | 6.61 | 6.30 | 5.59 | 5.33 |
| atp5f1c | pmol/100ug | 1.20 | 1.20 | 1.42 | 1.03 | 1.35 | 0.99 | 1.02 | 1.21 | 1.11 | 1.20 | 1.46 | 1.50 | 1.31 | 1.13 | 1.48 | 1.11 | 1.06 | 1.18 | 1.16 | 1.04 |
| atp5f1d | pmol/100ug | 0.73 | 0.63 | 0.74 | 0.56 | 0.63 | 0.63 | 0.61 | 0.63 | 0.55 | 0.56 | 0.77 | 0.78 | 0.64 | 0.59 | 0.62 | 0.56 | 0.61 | 0.53 | 0.51 | 0.50 |
| atp5f1e | pmol/100ug | 0.14 | 0.17 | 0.17 | 0.12 | 0.16 | 0.16 | 0.14 | 0.09 | 0.14 | 0.16 | 0.19 | 0.19 | 0.21 | 0.15 | 0.15 | 0.15 | 0.18 | 0.16 | 0.15 | 0.11 |
| atp5me | pmol/100ug | 0.14 | 0.11 | 0.19 | 0.17 | 0.22 | 0.13 | 0.15 | 0.12 | 0.15 | 0.16 | 0.16 | 0.21 | 0.25 | 0.18 | 0.19 | 0.14 | 0.20 | 0.21 | 0.24 | 0.13 |
| atp5mf | pmol/100ug | 1.19 | 1.32 | 1.36 | 1.19 | 1.54 | 1.16 | 1.30 | 1.39 | 1.33 | 1.21 | 1.32 | 1.44 | 1.79 | 1.19 | 1.36 | 1.33 | 1.55 | 1.53 | 1.67 | 1.22 |
| atp5mg | pmol/100ug | 0.13 | 0.14 | 0.17 | 0.13 | 0.19 | 0.14 | 0.17 | 0.18 | 0.13 | 0.15 | 0.18 | 0.22 | 0.21 | 0.18 | 0.18 | 0.19 | 0.18 | 0.20 | 0.25 | 0.15 |
| atp5mk | pmol/100ug | 0.08 | 0.10 | 0.11 | 0.07 | 0.07 | 0.10 | 0.06 | 0.09 | 0.10 | 0.07 | 0.07 | 0.09 | 0.09 | 0.06 | 0.07 | 0.06 | 0.06 | 0.08 | 0.08 | 0.06 |
| atp5pb | pmol/100ug | 0.96 | 0.89 | 1.04 | 0.84 | 0.82 | 0.87 | 0.83 | 0.97 | 0.95 | 1.02 | 1.26 | 1.08 | 1.13 | 0.80 | 1.10 | 1.00 | 0.85 | 1.20 | 0.89 | 0.84 |
| Atp5pd | pmol/100ug | 0.46 | 0.43 | 0.48 | 0.41 | 0.35 | 0.37 | 0.37 | 0.40 | 0.38 | 0.37 | 0.49 | 0.41 | 0.51 | 0.29 | 0.40 | 0.36 | 0.38 | 0.43 | 0.33 | 0.31 |
| atp5po | pmol/100ug | 0.88 | 0.85 | 0.98 | 0.82 | 0.66 | 0.69 | 0.70 | 0.87 | 0.87 | 0.78 | 0.91 | 0.80 | 1.22 | 0.60 | 0.82 | 0.74 | 0.73 | 1.03 | 0.71 | 0.65 |
| cox4i1 | pmol/100ug | 1.36 | 1.54 | 1.72 | 1.59 | 1.68 | 1.31 | 1.56 | 1.42 | 1.40 | 1.44 | 1.66 | 1.52 | 1.63 | 1.36 | 1.71 | 1.50 | 1.49 | 1.79 | 1.74 | 1.22 |
| cox5a | pmol/100ug | 0.68 | 0.53 | 0.68 | 0.57 | 0.42 | 0.74 | 0.41 | 0.60 | 0.51 | 0.34 | 0.45 | 0.57 | 0.51 | 0.40 | 0.43 | 0.43 | 0.46 | 0.51 | 0.44 | 0.36 |
| cox5b | pmol/100ug | 0.27 | 0.29 | 0.30 | 0.30 | 0.19 | 0.26 | 0.21 | 0.27 | 0.26 | 0.17 | 0.20 | 0.20 | 0.21 | 0.18 | 0.24 | 0.19 | 0.19 | 0.24 | 0.23 | 0.14 |
| cox6b1 | pmol/100ug | 0.18 | 0.22 | 0.26 | 0.24 | 0.23 | 0.23 | 0.23 | 0.22 | 0.23 | 0.21 | 0.25 | 0.25 | 0.25 | 0.25 | 0.22 | 0.21 | 0.30 | 0.28 | 0.27 | 0.16 |
| cox6c | pmol/100ug | 0.30 | 0.28 | 0.41 | 0.27 | 0.43 | 0.37 | 0.32 | 0.32 | 0.39 | 0.28 | 0.31 | 0.36 | 0.45 | 0.35 | 0.34 | 0.40 | 0.38 | 0.38 | 0.44 | 0.21 |
| cox7a2 | pmol/100ug | 0.09 | 0.10 | 0.12 | 0.09 | 0.09 | 0.13 | 0.08 | 0.04 | 0.12 | 0.07 | 0.10 | 0.09 | 0.08 | 0.06 | 0.09 | 0.08 | 0.11 | 0.10 | 0.06 | 0.06 |
| cox7c | pmol/100ug | 0.12 | 0.12 | 0.16 | 0.10 | 0.16 | 0.12 | 0.14 | 0.14 | 0.16 | 0.14 | 0.14 | 0.17 | 0.16 | 0.14 | 0.12 | 0.13 | 0.13 | 0.16 | 0.13 | 0.15 |
| cyc1 | pmol/100ug | 0.14 | 0.17 | 0.23 | 0.22 | 0.11 | 0.18 | 0.13 | 0.17 | 0.15 | 0.16 | 0.14 | 0.16 | 0.19 | 0.11 | 0.17 | 0.14 | 0.15 | 0.20 | 0.14 | 0.09 |
| cytb | pmol/100ug | 0.07 | 0.09 | 0.09 | 0.07 | 0.10 | 0.07 | 0.09 | 0.08 | 0.08 | 0.10 | 0.09 | 0.09 | 0.12 | 0.08 | 0.09 | 0.08 | 0.08 | 0.10 | 0.08 | 0.07 |
| Mtco1 | pmol/100ug | 0.16 | 0.28 | 0.36 | 0.22 | 0.34 | 0.18 | 0.29 | 0.21 | 0.30 | 0.34 | 0.30 | 0.24 | 0.29 | 0.28 | 0.32 | 0.28 | 0.27 | 0.31 | 0.27 | 0.21 |
| Mtco2 | pmol/100ug | 1.77 | 2.37 | 2.47 | 2.09 | 1.88 | 1.28 | 2.04 | 1.83 | 2.15 | 2.23 | 2.88 | 1.67 | 2.51 | 1.74 | 2.38 | 2.06 | 2.06 | 3.08 | 2.23 | 1.39 |
| Mtco3 | pmol/100ug | 0.11 | 0.10 | 0.15 | 0.12 | 0.14 | 0.10 | 0.11 | 0.06 | 0.16 | 0.13 | 0.11 | 0.13 | 0.08 | 0.12 | 0.15 | 0.11 | 0.12 | 0.17 | 0.13 | 0.10 |

| | | | | | | | | | | | | | | | | | | | | | |
|---|---|---|---|---|---|---|---|---|---|---|---|---|---|---|---|---|---|---|---|---|---|
| uqcr10 | pmol/100ug | 0.31 | 0.27 | 0.31 | 0.25 | 0.39 | 0.32 | 0.30 | 0.33 | 0.30 | 0.29 | 0.24 | 0.34 | 0.44 | 0.26 | 0.29 | 0.36 | 0.40 | 0.33 | 0.40 | 0.23 |
| uqcr11 | pmol/100ug | 0.07 | 0.08 | 0.09 | 0.07 | 0.07 | 0.09 | 0.06 | 0.08 | 0.07 | 0.05 | 0.07 | 0.10 | 0.12 | 0.05 | 0.07 | 0.07 | 0.08 | 0.12 | 0.09 | 0.05 |
| uqcrb | pmol/100ug | 0.18 | 0.27 | 0.30 | 0.37 | 0.33 | 0.31 | 0.27 | 0.37 | 0.26 | 0.22 | 0.22 | 0.24 | 0.31 | 0.25 | 0.24 | 0.29 | 0.31 | 0.31 | 0.38 | 0.22 |
| uqcrc1 | pmol/100ug | 1.35 | 1.71 | 1.65 | 1.42 | 1.41 | 1.18 | 1.20 | 1.20 | 1.59 | 1.32 | 1.25 | 1.55 | 1.59 | 1.06 | 1.42 | 1.17 | 1.39 | 1.32 | 1.10 | 1.06 |
| uqcrc2 | pmol/100ug | 0.76 | 0.72 | 1.08 | 0.83 | 0.96 | 0.72 | 0.81 | 0.78 | 0.90 | 0.78 | 0.78 | 0.94 | 1.22 | 0.63 | 0.98 | 0.84 | 0.84 | 0.94 | 0.90 | 0.58 |
| uqcrfs1 | pmol/100ug | 0.34 | 0.31 | 0.41 | 0.32 | 0.26 | 0.22 | 0.34 | 0.34 | 0.35 | 0.36 | 0.43 | 0.33 | 0.51 | 0.29 | 0.37 | 0.31 | 0.29 | 0.43 | 0.31 | 0.31 |
| uqcrh | pmol/100ug | 0.06 | 0.11 | 0.11 | 0.08 | 0.06 | 0.07 | 0.06 | 0.08 | 0.11 | 0.05 | 0.05 | 0.05 | 0.04 | 0.06 | 0.05 | 0.05 | 0.07 | 0.06 | 0.09 | 0.05 |
| uqcrq | pmol/100ug | 0.19 | 0.18 | 0.22 | 0.17 | 0.19 | 0.20 | 0.19 | 0.21 | 0.24 | 0.16 | 0.12 | 0.16 | 0.26 | 0.16 | 0.17 | 0.19 | 0.18 | 0.19 | 0.25 | 0.11 |
| Nd1 | pmol/100ug | 0.18 | 0.21 | 0.19 | 0.16 | 0.17 | 0.16 | 0.16 | 0.19 | 0.21 | 0.18 | 0.17 | 0.18 | 0.23 | 0.16 | 0.18 | 0.15 | 0.15 | 0.20 | 0.18 | 0.15 |
| Nd3 | pmol/100ug | 0.08 | 0.13 | 0.12 | 0.08 | 0.12 | 0.08 | 0.12 | 0.08 | 0.12 | 0.13 | 0.12 | 0.12 | 0.14 | 0.10 | 0.12 | 0.10 | 0.12 | 0.11 | 0.13 | 0.08 |
| Nd4 | pmol/100ug | 0.04 | 0.04 | 0.04 | 0.03 | 0.04 | 0.04 | 0.03 | 0.04 | 0.05 | 0.03 | 0.03 | 0.04 | 0.05 | 0.04 | 0.04 | 0.03 | 0.03 | 0.04 | 0.04 | 0.03 |
| Nd5 | pmol/100ug | 0.03 | 0.04 | 0.05 | 0.03 | 0.05 | 0.02 | 0.05 | 0.04 | 0.04 | 0.05 | 0.04 | 0.04 | 0.05 | 0.03 | 0.06 | 0.03 | 0.05 | 0.05 | 0.04 | 0.03 |
| Ndufa1 | pmol/100ug | 0.11 | 0.10 | 0.13 | 0.09 | 0.08 | 0.11 | 0.07 | 0.10 | 0.13 | 0.09 | 0.08 | 0.09 | 0.09 | 0.10 | 0.10 | 0.15 | 0.10 | 0.10 | 0.09 | 0.10 |
| Ndufa10 | pmol/100ug | 0.23 | 0.27 | 0.33 | 0.23 | 0.21 | 0.24 | 0.19 | 0.27 | 0.26 | 0.19 | 0.19 | 0.23 | 0.30 | 0.16 | 0.25 | 0.22 | 0.20 | 0.25 | 0.26 | 0.17 |
| Ndufa11 | pmol/100ug | 0.05 | 0.06 | 0.06 | 0.05 | 0.07 | 0.06 | 0.07 | 0.07 | 0.06 | 0.06 | 0.07 | 0.07 | 0.07 | 0.06 | 0.06 | 0.07 | 0.07 | 0.06 | 0.06 | 0.04 |
| Ndufa12 | pmol/100ug | 0.15 | 0.14 | 0.17 | 0.13 | 0.14 | 0.14 | 0.14 | 0.15 | 0.11 | 0.13 | 0.15 | 0.14 | 0.13 | 0.12 | 0.12 | 0.14 | 0.13 | 0.12 | 0.14 | 0.11 |
| Ndufa13 | pmol/100ug | 0.30 | 0.28 | 0.33 | 0.18 | 0.27 | 0.26 | 0.31 | 0.32 | 0.24 | 0.30 | 0.30 | 0.29 | 0.32 | 0.30 | 0.27 | 0.33 | 0.30 | 0.34 | 0.40 | 0.10 |
| Ndufa2 | pmol/100ug | 0.10 | 0.11 | 0.12 | 0.11 | 0.08 | 0.10 | 0.08 | 0.09 | 0.10 | 0.07 | 0.08 | 0.06 | 0.09 | 0.08 | 0.09 | 0.08 | 0.08 | 0.09 | 0.08 | 0.06 |
| Ndufa4 | pmol/100ug | 0.31 | 0.38 | 0.40 | 0.41 | 0.36 | 0.40 | 0.38 | 0.39 | 0.33 | 0.31 | 0.36 | 0.36 | 0.47 | 0.33 | 0.35 | 0.38 | 0.36 | 0.41 | 0.46 | 0.28 |
| Ndufa5 | pmol/100ug | 0.07 | 0.07 | 0.08 | 0.06 | 0.06 | 0.08 | 0.06 | 0.07 | 0.07 | 0.05 | 0.06 | 0.07 | 0.08 | 0.06 | 0.07 | 0.07 | 0.06 | 0.08 | 0.07 | 0.05 |
| Ndufa6 | pmol/100ug | 0.13 | 0.11 | 0.13 | 0.13 | 0.14 | 0.14 | 0.14 | 0.15 | 0.12 | 0.10 | 0.10 | 0.11 | 0.15 | 0.14 | 0.12 | 0.12 | 0.14 | 0.13 | 0.16 | 0.13 |
| Ndufa8 | pmol/100ug | 0.16 | 0.14 | 0.16 | 0.11 | 0.13 | 0.14 | 0.15 | 0.16 | 0.14 | 0.14 | 0.16 | 0.17 | 0.14 | 0.15 | 0.14 | 0.12 | 0.13 | 0.14 | 0.13 | 0.16 |
| Ndufa9 | pmol/100ug | 0.12 | 0.15 | 0.18 | 0.14 | 0.12 | 0.13 | 0.12 | 0.16 | 0.15 | 0.11 | 0.12 | 0.13 | 0.20 | 0.11 | 0.14 | 0.14 | 0.12 | 0.15 | 0.16 | 0.12 |
| Ndufb10 | pmol/100ug | 0.38 | 0.30 | 0.39 | 0.27 | 0.26 | 0.25 | 0.25 | 0.33 | 0.34 | 0.29 | 0.27 | 0.30 | 0.37 | 0.26 | 0.30 | 0.28 | 0.31 | 0.38 | 0.29 | 0.21 |
| Ndufb11 | pmol/100ug | 0.11 | 0.07 | 0.08 | 0.05 | 0.05 | 0.11 | 0.05 | 0.07 | 0.08 | 0.07 | 0.05 | 0.09 | 0.05 | 0.07 | 0.07 | 0.07 | 0.05 | 0.05 | 0.04 | 0.06 |
| Ndufb2 | pmol/100ug | 0.05 | 0.04 | 0.04 | 0.04 | 0.05 | 0.04 | 0.05 | 0.05 | 0.04 | 0.04 | 0.04 | 0.04 | 0.05 | 0.03 | 0.04 | 0.04 | 0.05 | 0.04 | 0.04 | 0.03 |
| Ndufb3 | pmol/100ug | 0.14 | 0.10 | 0.16 | 0.12 | 0.17 | 0.12 | 0.16 | 0.16 | 0.12 | 0.12 | 0.13 | 0.15 | 0.13 | 0.13 | 0.13 | 0.13 | 0.14 | 0.14 | 0.16 | 0.12 |

| | | | | | | | | | | | | | | | | | | | | | |
|---|---|---|---|---|---|---|---|---|---|---|---|---|---|---|---|---|---|---|---|---|---|
| Ndufb5 | pmol/100ug | 0.14 | 0.14 | 0.24 | 0.14 | 0.13 | 0.12 | 0.17 | 0.17 | 0.12 | 0.17 | 0.19 | 0.11 | 0.21 | 0.14 | 0.18 | 0.14 | 0.18 | 0.15 | 0.23 | 0.12 |
| Ndufb6 | pmol/100ug | 0.07 | 0.06 | 0.07 | 0.05 | 0.05 | 0.06 | 0.06 | 0.07 | 0.05 | 0.07 | 0.05 | 0.05 | 0.08 | 0.06 | 0.06 | 0.06 | 0.05 | 0.06 | 0.06 | 0.05 |
| Ndufb7 | pmol/100ug | 0.12 | 0.18 | 0.18 | 0.12 | 0.15 | 0.13 | 0.19 | 0.14 | 0.18 | 0.17 | 0.15 | 0.16 | 0.17 | 0.16 | 0.15 | 0.15 | 0.16 | 0.17 | 0.17 | 0.14 |
| Ndufb9 | pmol/100ug | 0.12 | 0.15 | 0.16 | 0.13 | 0.07 | 0.12 | 0.07 | 0.10 | 0.15 | 0.06 | 0.08 | 0.08 | 0.10 | 0.07 | 0.06 | 0.09 | 0.07 | 0.08 | 0.07 | 0.06 |
| Ndufc2 | pmol/100ug | 0.16 | 0.15 | 0.18 | 0.15 | 0.17 | 0.16 | 0.18 | 0.19 | 0.16 | 0.13 | 0.16 | 0.13 | 0.13 | 0.16 | 0.15 | 0.13 | 0.14 | 0.18 | 0.16 | 0.10 |
| Ndufs1 | pmol/100ug | 0.83 | 0.96 | 0.95 | 0.68 | 0.97 | 0.75 | 0.85 | 0.77 | 0.96 | 0.99 | 0.91 | 0.93 | 1.03 | 0.87 | 0.95 | 0.81 | 0.92 | 1.04 | 0.90 | 0.74 |
| Ndufs2 | pmol/100ug | 0.24 | 0.30 | 0.32 | 0.23 | 0.21 | 0.22 | 0.25 | 0.26 | 0.30 | 0.23 | 0.19 | 0.24 | 0.28 | 0.21 | 0.23 | 0.20 | 0.19 | 0.21 | 0.19 | 0.20 |
| Ndufs3 | pmol/100ug | 0.44 | 0.44 | 0.50 | 0.37 | 0.45 | 0.44 | 0.43 | 0.50 | 0.40 | 0.48 | 0.53 | 0.48 | 0.61 | 0.39 | 0.48 | 0.49 | 0.45 | 0.63 | 0.53 | 0.43 |
| Ndufs4 | pmol/100ug | 0.12 | 0.11 | 0.15 | 0.12 | 0.13 | 0.11 | 0.15 | 0.13 | 0.11 | 0.13 | 0.16 | 0.14 | 0.18 | 0.15 | 0.14 | 0.16 | 0.13 | 0.14 | 0.15 | 0.11 |
| Ndufs5 | pmol/100ug | 0.18 | 0.17 | 0.18 | 0.17 | 0.19 | 0.17 | 0.18 | 0.16 | 0.16 | 0.17 | 0.17 | 0.17 | 0.18 | 0.20 | 0.18 | 0.17 | 0.15 | 0.21 | 0.20 | 0.17 |
| Ndufs6 | pmol/100ug | 0.05 | 0.03 | 0.05 | 0.04 | 0.05 | 0.05 | 0.03 | 0.03 | 0.04 | 0.04 | 0.05 | 0.05 | 0.03 | 0.05 | 0.03 | 0.04 | 0.04 | 0.04 | 0.04 | 0.04 |
| Ndufs7 | pmol/100ug | 0.21 | 0.20 | 0.24 | 0.19 | 0.17 | 0.17 | 0.20 | 0.23 | 0.19 | 0.19 | 0.21 | 0.19 | 0.20 | 0.18 | 0.21 | 0.20 | 0.19 | 0.25 | 0.20 | 0.16 |
| Ndufs8 | pmol/100ug | 0.33 | 0.39 | 0.38 | 0.41 | 0.37 | 0.34 | 0.39 | 0.41 | 0.40 | 0.41 | 0.47 | 0.35 | 0.49 | 0.35 | 0.41 | 0.37 | 0.41 | 0.48 | 0.43 | 0.35 |
| Ndufv1 | pmol/100ug | 0.46 | 0.45 | 0.53 | 0.40 | 0.41 | 0.41 | 0.40 | 0.41 | 0.45 | 0.43 | 0.40 | 0.43 | 0.38 | 0.41 | 0.40 | 0.38 | 0.41 | 0.34 | 0.35 | 0.33 |
| Ndufv2 | pmol/100ug | 0.17 | 0.18 | 0.21 | 0.15 | 0.08 | 0.15 | 0.10 | 0.19 | 0.18 | 0.10 | 0.13 | 0.10 | 0.13 | 0.09 | 0.13 | 0.11 | 0.10 | 0.16 | 0.12 | 0.09 |

**Table S7**. Core glycolysis/gluconeogenesis genes significantly downregulated in si*MLKL* HepG2 cells

| Gene | Fold Change (siMLKL vs siControl) | Gene Name | Significance (padj) |
|---|---|---|---|
| HK2 | log2FC = −0.674 (≈0.63-fold, ~37% decrease) | Hexokinase 2 | $5.07 \times 10^{-21}$ |
| PDK1 | log2FC = −0.624 (≈0.65-fold, ~35% decrease) | Pyruvate Dehydrogenase Kinase 1 | $3.40 \times 10^{-14}$ |
| PFKFB3 | log2FC = −0.643 (≈0.64-fold, ~36% decrease) | 6-Phosphofructo-2-Kinase/Fructose-2,6-Biphosphatase 3 | $1.62 \times 10^{-7}$ |

**Table S8**: Targeted mitochondrial proteomics data for glycolysis proteins in the livers of Control or *Mlkl*$^{HepKO}$ mice fed either CD or WD for 8 or 15 months

| **Genotype** | | **Control** | **Control** | **Control** | **Control** | **Control** | **Control** | **Control** | **Control** | **Control** | **Control** | ***Mlkl HepKO*** | ***Mlkl HepKO*** | ***Mlkl HepKO*** | ***Mlkl HepKO*** | ***Mlkl HepKO*** | ***Mlkl HepKO*** | ***Mlkl HepKO*** | ***Mlkl HepKO*** | ***Mlkl HepKO*** | ***Mlkl HepKO*** |
|---|---|---|---|---|---|---|---|---|---|---|---|---|---|---|---|---|---|---|---|---|---|
| **Diet** | | **CD** | **CD** | **CD** | **CD** | **CD** | **WD** | **WD** | **WD** | **WD** | **WD** | **CD** | **CD** | **CD** | **CD** | **CD** | **WD** | **WD** | **WD** | **WD** | **WD** |
| **Protein** | **Peptide** | **CD_ 1** | **CD_ 2** | **CD_ 3** | **CD_ 4** | **CD_ 5** | **WD_ 1** | **WD_ 2** | **WD_ 3** | **WD_ 4** | **WD_ 5** | **CD_ 1** | **CD_ 2** | **CD_ 3** | **CD_ 4** | **CD_ 4** | **WD_ 1** | **WD_ 2** | **WD_ 3** | **WD_ 4** | **WD_ 5** |
| **8 Months** | | | | | | | | | | | | | | | | | | | | | |
| Gpi | pmol/100ug | 0.426 | 0.414 | 0.486 | 0.516 | 0.468 | 0.776 | 0.445 | 0.822 | 0.323 | 0.716 | 0.439 | 0.554 | 0.436 | 0.399 | 0.318 | 0.538 | 0.536 | 0.493 | 0.511 | 0.572 |
| Pfkm | pmol/100ug | 0.168 | 0.157 | 0.142 | 0.160 | 0.146 | 0.185 | 0.143 | 0.219 | 0.191 | 0.171 | 0.155 | 0.160 | 0.149 | 0.139 | 0.174 | 0.233 | 0.232 | 0.199 | 0.207 | 0.226 |
| Aldoa | pmol/100ug | 0.207 | 0.200 | 0.157 | 0.217 | 0.193 | 0.383 | 0.208 | 0.380 | 0.407 | 0.266 | 0.221 | 0.280 | 0.210 | 0.138 | 0.167 | 0.294 | 0.304 | 0.218 | 0.198 | 0.286 |
| Aldob | pmol/100ug | 16.986 | 19.453 | 19.880 | 18.388 | 16.468 | 27.053 | 23.557 | 28.172 | 34.226 | 27.696 | 17.892 | 23.212 | 21.303 | 13.739 | 12.560 | 22.680 | 16.453 | 14.139 | 20.626 | 14.786 |
| Tpi1 | pmol/100ug | 1.585 | 1.641 | 1.550 | 1.619 | 1.433 | 1.850 | 1.247 | 2.008 | 1.648 | 1.591 | 1.752 | 1.895 | 1.692 | 1.385 | 1.002 | 1.321 | 1.156 | 1.162 | 1.203 | 1.063 |
| Gapdh | pmol/100ug | 14.290 | 13.959 | 19.009 | 15.224 | 12.005 | 19.516 | 15.498 | 23.711 | 22.028 | 18.478 | 12.380 | 14.327 | 14.713 | 9.003 | 8.892 | 10.917 | 12.133 | 9.378 | 12.682 | 10.642 |
| Pgk1 | pmol/100ug | 1.410 | 1.268 | 1.475 | 1.441 | 1.317 | 1.313 | 1.130 | 1.319 | 1.416 | 1.261 | 1.354 | 1.376 | 1.267 | 0.904 | 0.861 | 1.023 | 0.918 | 0.725 | 0.961 | 0.840 |
| Pgam1 | pmol/100ug | 0.200 | 0.193 | 0.219 | 0.218 | 0.182 | 0.195 | 0.172 | 0.246 | 0.243 | 0.207 | 0.189 | 0.210 | 0.201 | 0.165 | 0.111 | 0.162 | 0.143 | 0.160 | 0.141 | 0.152 |
| Eno1 | pmol/100ug | 4.043 | 4.239 | 4.817 | 4.357 | 4.022 | 6.324 | 4.299 | 6.384 | 6.320 | 5.092 | 3.897 | 4.888 | 5.032 | 3.363 | 3.022 | 4.565 | 4.506 | 4.001 | 4.279 | 4.710 |
| Pkm | pmol/100ug | 0.535 | 0.421 | 0.391 | 0.475 | 0.433 | 0.574 | 0.350 | 0.707 | 0.641 | 0.567 | 0.393 | 0.405 | 0.394 | 0.555 | 0.829 | 0.901 | 0.927 | 0.747 | 0.598 | 0.764 |
| Pklr | pmol/100ug | 1.676 | 1.517 | 1.728 | 1.234 | 1.451 | 1.576 | 1.259 | 1.616 | 1.832 | 1.500 | 1.110 | 1.340 | 1.533 | 1.205 | 1.198 | 1.251 | 1.182 | 1.291 | 1.144 | 1.044 |
| Ldha | pmol/100ug | 10.581 | 8.700 | 9.881 | 9.201 | 8.315 | 10.599 | 9.901 | 9.525 | 11.140 | 10.380 | 9.318 | 9.948 | 10.061 | 7.169 | 8.544 | 10.227 | 9.335 | 8.513 | 9.385 | 8.237 |
| **15 Months** | | | | | | | | | | | | | | | | | | | | | |
| Gpi | pmol/100ug | 0.405 | 0.489 | 0.398 | 0.347 | 0.576 | 0.427 | 0.631 | 0.891 | 0.585 | 0.358 | 0.376 | 0.632 | 0.361 | 0.367 | 0.357 | 0.637 | 0.627 | 0.556 | 0.382 | 0.437 |
| Pfkm | pmol/100ug | 0.157 | 0.179 | 0.198 | 0.173 | 0.234 | 0.195 | 0.226 | 0.310 | 0.223 | 0.171 | 0.218 | 0.212 | 0.194 | 0.191 | 0.198 | 0.256 | 0.205 | 0.220 | 0.218 | 0.190 |
| Aldoa | pmol/100ug | 0.157 | 0.204 | 0.177 | 0.133 | 0.321 | 0.326 | 0.293 | 0.291 | 0.217 | 0.110 | 0.199 | 0.170 | 0.141 | 0.115 | 0.143 | 0.294 | 0.239 | 0.259 | 0.137 | 0.144 |
| Aldob | pmol/100ug | 15.056 | 20.410 | 17.647 | 14.698 | 14.520 | 11.301 | 14.524 | 9.963 | 13.126 | 10.669 | 8.497 | 14.232 | 11.762 | 11.678 | 12.682 | 15.429 | 10.845 | 16.846 | 12.916 | 11.864 |
| Tpi1 | pmol/100ug | 1.138 | 1.534 | 1.105 | 1.473 | 1.426 | 1.266 | 1.267 | 1.646 | 1.076 | 0.983 | 1.476 | 1.146 | 1.230 | 1.256 | 1.093 | 1.315 | 1.219 | 1.219 | 1.026 | 1.142 |
| Gapdh | pmol/100ug | 8.381 | 11.964 | 8.965 | 8.758 | 11.974 | 8.636 | 10.682 | 11.690 | 11.366 | 7.927 | 9.350 | 12.191 | 10.729 | 9.264 | 9.835 | 13.443 | 11.164 | 14.933 | 9.875 | 10.334 |
| Pgk1 | pmol/100ug | 1.100 | 1.255 | 0.955 | 0.826 | 1.066 | 0.819 | 0.789 | 0.934 | 0.769 | 0.785 | 0.938 | 1.113 | 0.962 | 0.661 | 0.995 | 0.888 | 0.804 | 1.130 | 0.675 | 0.587 |
| Pgam1 | pmol/100ug | 0.120 | 0.153 | 0.120 | 0.126 | 0.186 | 0.130 | 0.131 | 0.164 | 0.137 | 0.104 | 0.168 | 0.139 | 0.145 | 0.135 | 0.116 | 0.168 | 0.147 | 0.173 | 0.124 | 0.132 |
| Eno1 | pmol/100ug | 3.706 | 4.752 | 3.743 | 3.490 | 5.247 | 3.702 | 4.035 | 5.054 | 3.918 | 3.189 | 2.891 | 4.028 | 3.726 | 3.435 | 3.289 | 4.181 | 4.383 | 3.862 | 3.720 | 3.638 |
| Pkm | pmol/100ug | 0.556 | 0.617 | 0.599 | 0.298 | 0.765 | 0.729 | 0.695 | 1.091 | 1.052 | 1.006 | 1.656 | 0.984 | 0.970 | 1.095 | 0.701 | 0.974 | 0.955 | 0.966 | 0.929 | 0.610 |
| Pklr | pmol/100ug | 1.490 | 1.751 | 1.636 | 1.044 | 1.168 | 0.979 | 1.161 | 1.666 | 1.228 | 1.575 | 0.868 | 1.955 | 1.542 | 1.696 | 1.219 | 1.328 | 1.306 | 1.273 | 0.939 | 0.946 |
| Ldha | pmol/100ug | 8.230 | 9.290 | 7.760 | 9.453 | 9.003 | 8.510 | 8.978 | 10.468 | 7.323 | 6.889 | 9.509 | 10.154 | 8.962 | 8.130 | 7.252 | 10.609 | 8.468 | 10.772 | 7.523 | 8.935 |

**Table S9**. Plasma biochemistry of xenograft mice with control or NSA treatment

| Parameters (Unit) | Control | NSA |
|---|---|---|
| BUN ( mg/dL) | 17.75 ± 3.95 | 19.60 ± 3.78 |
| ALT (U/L) | 30.50 ± 5.00 | 32.20 ± 2.95 |
| AST (U/L) | 105.25 ± 31.27 | 80.60 ± 14.24 |
| GLU (mg/dL) | 246.50 ± 22.88 | 265.60 ± 26.83 |
| TP (g/dL) | 5.25 ± 0.21 | 4.98 ± 0.27 |
| ALB (g/dL) | 4.13 ± 0.25 | 3.94 ± 0.13 |
| GLOB (g/dL) | 1.10 ± 0.24 | 1.06 ± 0.19 |

The Creatinine (CRE) showed values below the detection limits (<0.2 mg/dL).

Abbreviations: BUN, blood urea nitrogen; ALT, alanine transaminase; AST, aspartate transaminase; GLU, glucose; TP, total protein; ALB, albumin; GLOB, globulin.
